# H2A.Z-dependent and -independent recruitment of metabolic enzymes to chromatin required for histone modifications

**DOI:** 10.1101/553297

**Authors:** Sujung Choi, Yong Heui Jeon, Zhi Yang, Minzhen He, Hyewon Shin, Jessica Pfleger, Danish Sayed, Sophie Astrof, Maha Abdellatif

## Abstract

H2A.Z plays a fundamental role in the regulation of transcription and epigenetics, however, the mechanisms that underlie its functions are not fully understood. Using rapid chromatin immunoprecipitation-mass spectrometry, we uncovered the association of H2A.Z-bound chromatin with an array of tricarboxylic acid cycle and beta-oxidation enzymes in the mouse heart. Recombinant green florescence fusion proteins combined with mutations of putative nuclear localization signals of select enzymes, including acetyl-CoA acyltransferase 2 (ACAA2), oxoglutarate dehydrogenase (OGDH), and isocitrate dehydrogenase 2 confirmed their nuclear localization and chromatin binding in both rodent and human cells. Conclusively, chromatin immunoprecipitation-deep sequencing, confirmed the selective association of ACAA2 and OGDH with H2A.Z-occupied transcription start sites. Finally, human H2A.Z-deficient HAP1 cells exhibited reduced chromatin-bound metabolic enzymes, with the exception of pyruvate dehydrogenase, accompanied with reduced posttranslational histone modifications. Thus, the data show that metabolic enzymes are recruited to active promoters for potential site-directed epigenetic modifications.

## INTRODUCTION

The highly conserved, histone variant, H2A.Z gene is unique in many ways when contrasted with those of core histones. Mainly, it is a single copy gene that does not exist within the histone clusters known in human and mouse genomes, it includes introns, and is polyadenylated ^1^, all of which underscore its specialized nature. We currently know that it selectively associates with transcriptionally active, as well as, inactive genes. For example, in yeast, Htz1 has been shown to suppress the spread of heterochromatin into transcriptionally active genes near the telomeres ^2^, while, in contrast, its abundance was shown to negatively correlate with transcriptional rates ^3^. Furthermore, changes in growth conditions induced translocation of Htz1 from transcriptionally active to inactive genes ^4^. More precisely, Htz1 is found at the transcription start site (TSS) of nearly all genes in euchromatin, in the −1 and +1 nucleosomes flanking a nucleosome free region in the active genes, while present mainly in the −1 nucleosome in inactive genes ^5^. Likewise, in *Drosophila*, H2Av is present at thousands of both transcriptionally active and inactive genes in euchromatin, as well as, in heterochromatic chromocenter of polytene chromosomes ^6^, whereas its density negatively correlates with that of RNA polymerase II (pol II). Conversely, other studies have shown that H2A.Z vs. H2A at the +1 nucleosome facilitates pol II progression ^7^. In murine embryonic stem cells, we also see this bifunctionality, where in the undifferentiated state, H2A.Z, in conjunction with the polycomb subunit Suz12, is present at silenced homeodomain genes involved in differentiation, whereas in committed neuronal progenitor cells, its associates with highly expressed genes ^8^. Other than destabilizing nucleosomes at the +1 position, we have little understanding of how H2A.Z selectively regulates transcriptional activation v. deactivation, or if it has any role in metabolism-induced transcriptional remodeling.

Generally, organisms respond to metabolic cues by exacting a change in gene transcription that influences their development and growth, or homeostasis. These signals include ATP:ADP:AMP and NAD^+^:NADH ratios, and the availability of metabolites that are involved in histone and DNA modifications - e.g. acetyl-CoA (ac-CoA), α-ketoglutarate (αKG), and succinyl-CoA (suc-CoA), not discounting other acyl-CoAs ^9^. In the case of acetyl-CoA, we know that during substrate abundance, citrate is exported from the mitochondria and into the cytosol and nucleus, where ATP citrate lyase (ACLY) converts it into acetyl-CoA, which is a substrate for histone acetylation ^10^. Alternatively, during substrate shortage, acetate is imported from the circulation and converted into acetyl-CoA by acyl-CoA synthetase short chain family member 2 (ACSS2) ^11^. As for the other CoA-linked metabolites and αKG, the mechanism for nuclear delivery is less well-established. Moreover, the question of how substrates modulate the expression of specific target genes remains a challenge.

The current dogma is that metabolic oxidative enzymes and substrate oxidation are largely confined to the mitochondria. However, recent findings, including our own, challenge this belief. In specific, Sutendra et al, reported the presence of all subunits of the pyruvate dehydrogenase (PDH) complex in human sperm, and normal and cancerous lung epithelial cells ^12^, while Nagaraj et al reported it in the human 4/8-cell stage zygote ^13^, and show that it is required for generating acetyl-CoA for histone acetylation. Likewise, oxoglutarate dehydrogenase (OGDH) ^14^ and isocitrate dehydrogenase (IDH2) ^15^ have also been shown to partly localize to the nucleus, as these findings are further supported by a proteomics study that revealed the presence of all tricarboxylic acid (TCA) cycle enzymes in the nucleus of breast cancer cells ^16^. However, except for OGDH in cancer cells ^14^, none of these enzymes have been shown to associate with chromatin. Here we report, that using an unbiased screen for the discovery of proteins that co-localize to H2A.Z-bound chromatin, we uncovered its association with all the TCA cycle and β-oxidation enzymes in the nuclei of mice hearts. Our recent report shows that H2A.Z is mainly localized to the TSS of most transcribed genes, including housekeeping and inducible genes, where it is at its highest levels, but is relatively low at tissue-specific genes ^17^. Therefore, this would position the metabolic enzymes at the TSS of the former groups. In validation, we successfully completed chromatin immunoprecipitation-deep sequencing (ChIP-Seq) using anti-ACAA2 or OGDH, which we found associated with select H2A.Z-bound TSSs, the latter in both mouse heart and human colon cancer cells. Moreover, the knockout of H2A.Z in HAP1 cell provided evidence that this association is indeed dependent on H2A.Z for all of the enzymes tested, with the exception of pyruvate dehydrogenase A1 (PDHA1), which led us to conclude that the recruitment of metabolic enzymes is predominantly, but not exclusively, H2A.Z-dependent. The data also suggest that histone modifications are reliant on the recruitment of mitochondrial enzymes to chromatin.

## RESULTS

### Mitochondrial TCA cycle and β-oxidation enzymes localize to the nucleus, in association with H2A.Z-bound chromatin

We have recently reported that the highly conserved histone variant H2A.Z is assembled at the TSS of all housekeeping and inducible genes in the heart, where it interacts with acidic nuclear protein 32e (ANP32E) to differentially regulate gene expression ^17^. However, cardiac-specific genes (e.g. *Myh6*, *Actc1*‥etc), which are most highly expressed, have relatively low or no bound H2A.Z. This suggested that H2A.Z plays a more intricate role in regulating transcription than currently recognized and is plausibly mediated by protein recruitments to TSSs. In an attempt to discover proteins that associate with H2A.Z-bound nucleosomes, we performed rapid immunoprecipitation-mass spectrometry of endogenous proteins assay ^18^ (RIME, Fig. 1a), which involved chromatin-immunoprecipitation by anti-H2A.Z or a control IgG from the nuclei of mice hearts subjected to transverse aortic constriction (TAC) to induce growth, or a sham operation, followed by mass spectrometry. A total of 73 proteins with a cutoff of 2x enrichment v. the IgG control were identified. Of those, 36 of the most enriched are plotted as total spectra of proteins identified in the anti-H2A.Z immunocomplexes from the normal (sham-operated) and growth-induced (TAC) adult hearts, versus the IgG pull-down. First, as evidence of the efficacy of the RIME, we identified H2A.Z, as well as, core histones in the immunoprecipitated complex (Fig. 1b). Intriguingly, though, the proteins with the highest spectra and enrichment were those of the TCA cycle and the β-oxidation pathway (Fig. 1c-d). Moreover, all the enzymes of both pathways were identified in the chromatin precipitated complex, including enoyl-CoA delta isomerase 1 (ECI1), which is necessary for the oxidation of unsaturated fatty acids (Fig. 1c-d). All proteins were relatively equal in the sham v. TAC hearts, except for the PDH complex subunits, which were ~1.4-fold higher in the TAC hearts. Other genes that were significantly enriched in the RIME complex, included enzymes of branched-chain amino acid metabolism (7 enzymes), and protein translation (17 proteins), cytosolic (5 proteins), other mitochondrial (4 proteins), and intermediate filament proteins (3 proteins; supplementary Fig. 1S). Notably, except for the core histones, no transcription factors or regulators were identified in the precipitate. Thus, a preponderance of mitochondrial enzymes involved in glucose, fatty acid, and branched-chain amino acid oxidation reside in the nucleus, in association with H2A.Z-bound chromatin, in both the normal and hypertrophied hearts. This suggests that metabolites required for histone and DNA modifications are directly delivered to the transcription start sites. Although, this is a powerful approach and highly reproducible with two independent samples (sham and TAC hearts, pool of 20 each), caution, however, has to be exercised in interpreting the data pending its validation by other methods, as described below.

**Figure 1.**
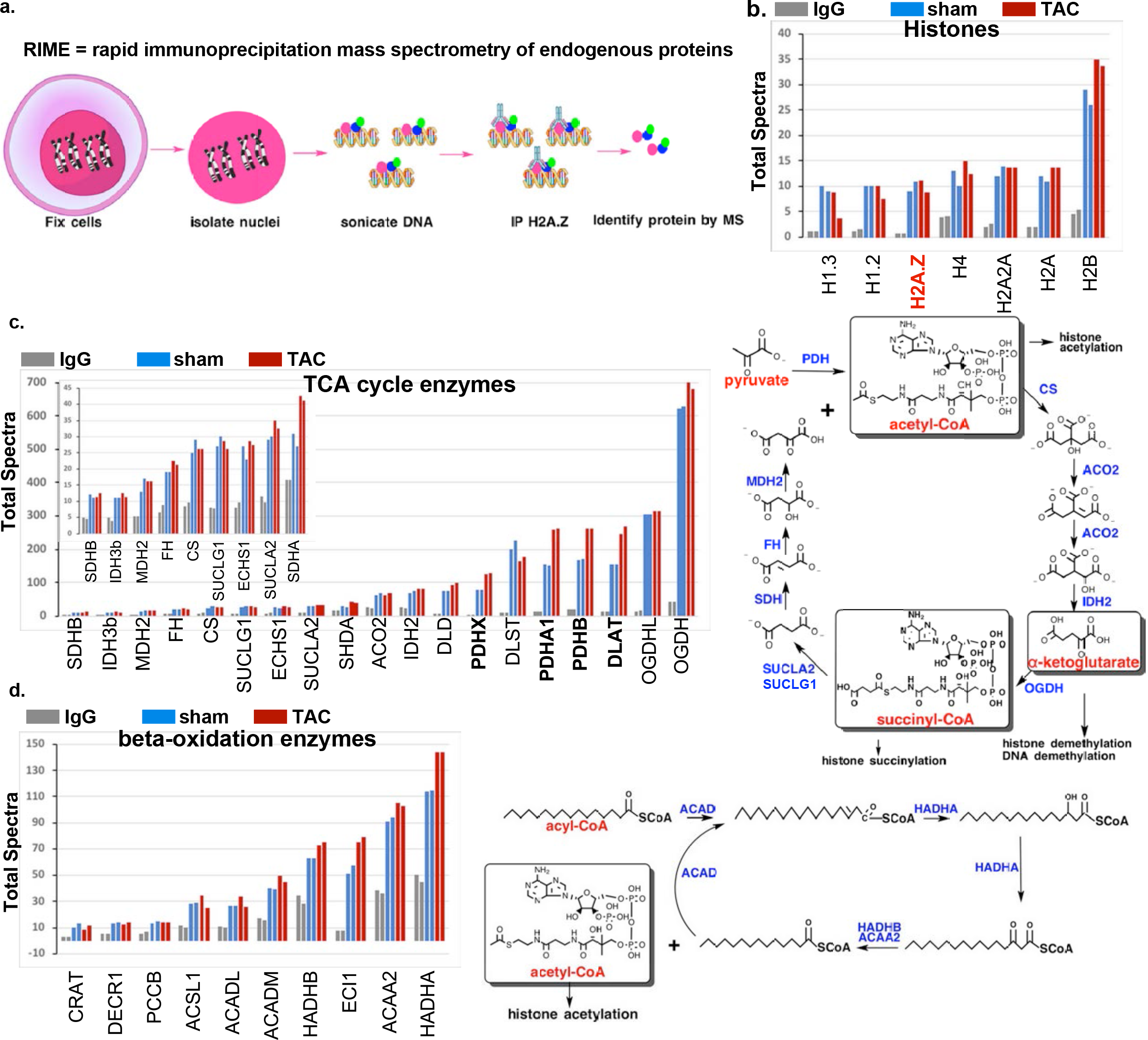
H2A.Z-bound chromatin associates with metabolic enzymes in mouse heart nuclei. ChIP with anti-H2A.Z or a control IgG was performed on nuclei from a pool of 10 hearts each, from mice subjected to a sham or a transverse aortic constriction operation. The sample size was determined based on the quantity of the material required for the assay and for biological averaging of samples based on our prior experience of the variability between mouse hearts in response to growth induction. The ChIP complex was then subjected to MS/MS, each sample analyzed twice. **b.-d.** Total spectra identified in the IgG, sham, and TAC samples were plotted. These include those with a cutoff of more than 2-fold enrichment v. control IgG, after normalization to the IgG C chain region spectra detected in each sample. Data were grouped as, **b.** histones, **c.** TCA cycle enzymes, and **d.** β-oxidation spiral enzymes. The enzymes identified are also indicated in blue in the pathways outlined for the TCA cycle and the β-oxidation spiral on the right in c. and d. Source file for these data are available in Figure 1-source data 1.

### Confirming the nuclear localization of metabolic enzymes in rodent and human cells

The above data suggest that TCA cycle and β-oxidation enzymes, among others, are not confined to mitochondria, but also localize to the nucleus. To confirm, we first used immunocytochemistry (ICC) staining of four mitochondrial enzymes in four cell types, including cultured rat neonatal cardiac myocytes (rNCM), isolated mouse adult cardiac myocytes (mACM), human iPSC-derived cardiac myocytes (hiPSC-CM), and SW620 colon cancer cells. The results revealed the nuclear localization of OGDH (which had the highest levels of spectra in the RIME assay and 15.8 ± 0.5-fold enrichment / IgG), IDH2 (2.8 ± 0.11-fold enrichment / IgG), PDHA1, 14.8 ± 2.25-fold enrichment / IgG), and ACAA2 (2.5 0.1-fold enrichment / IgG, Fig. 2a-d). OGDH, particularly, showed predominant nuclear localization in neonatal myocytes, hiPSC-CM, and SW620 (Fig. 2a), and in the developing embryonic heart (Fig. 2b), but is more evenly distributed in the mitochondria and nuclei of adult mouse myocytes. The antibody used in these images targets an epitope near the C-terminus region of the protein (E1W8H, Cell Signaling Technology). We also confirmed this finding with a second antibody against the N-terminal domain of the protein (Sigma, cat # HPA019514). While it detected nuclear OGDH, it had a stronger affinity to the mitochondrial protein (supplementary Fig. 2S). Moreover, images from the Human Protein Atlas ^19^ show OGDH (also using Sigma, cat # HPA019514) and PDH subunit b (PDHB) in the nuclei of cardiac myocytes in normal adult heart tissue, and OGDH in A-431 squamous carcinoma cells (supplementary Fig. 2S). Thus, it is critical to note that the immune-detection of these enzymes depends on the antibody, which may have differential affinities to the mitochondrially-vs. nuclearly-localized enzyme, plausibly due to differential display of the antigenic epitopes.

**Figure 2.**
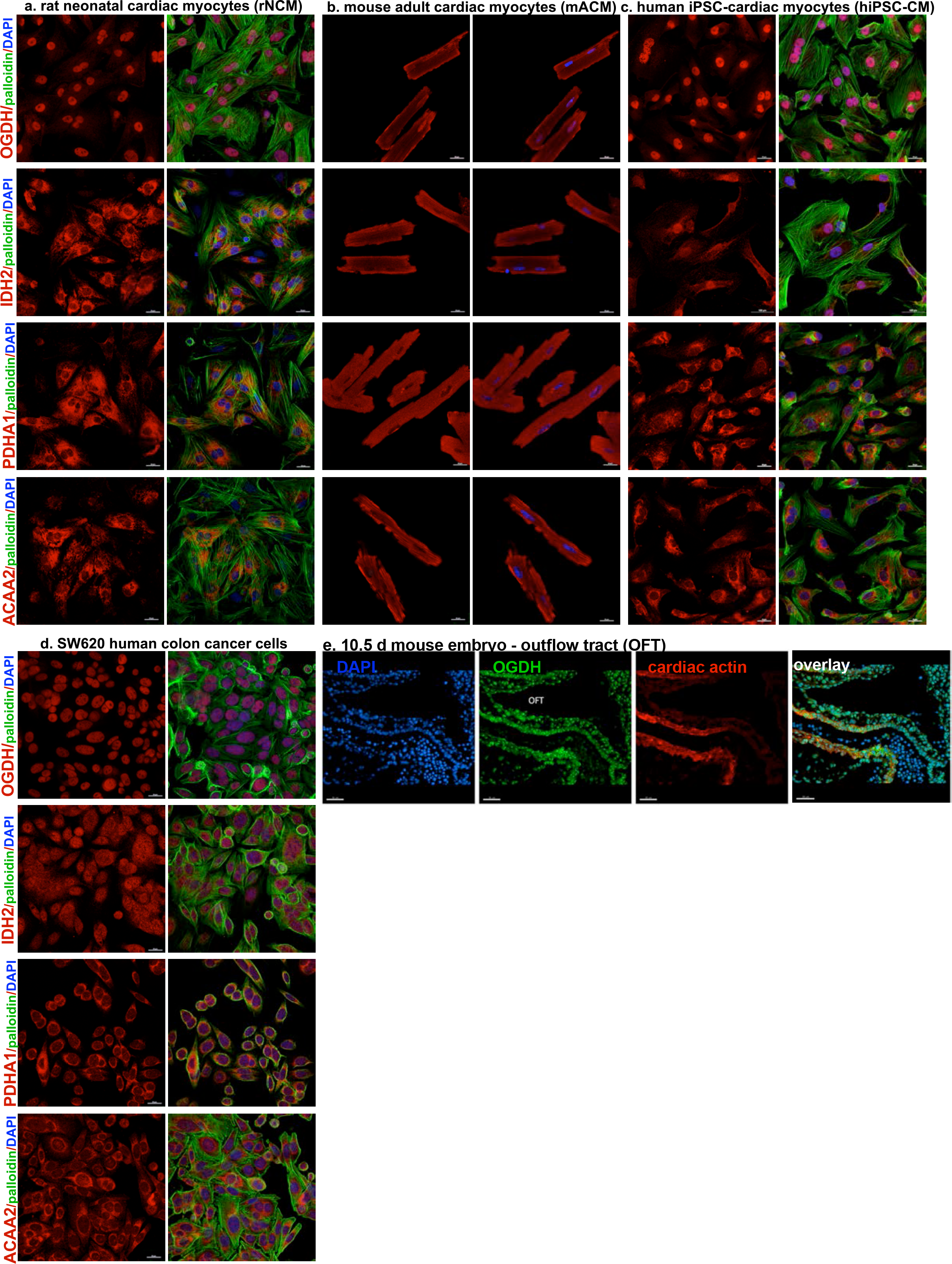
Nuclear localization of metabolic enzymes in both rodent and human cardiac myocytes, and colon cancer cells, observed by confocal imaging. **a.** Isolated rat neonatal cardiac myocytes (rNCM), **b.** mouse adult cardiac myocytes (mACM), **c.** human iPSC-derived cardiac myocytes (hiPSC-CM), and **d.** SW620 colon cancer cells, were cultured, fixed, and immune-stained with the antibodies for the proteins indicated on the left (anti-OGDH, -IDH2, -PDHA1, and -ACAA2, all in red), in addition to phalloidin (green), and DAPI (blue). For each, the left panels show the staining for OGDH, IDH2, PDHA1, or ACAA2 (red) and the right panel, their overlay with phalloidin (except for the mACM) and DAPI. The cells were then imaged using confocal microscopy. **e.** Ten and a half-day-old mouse embryos were immune-stained for OGDH (green), α-actinin (red), and DAPI (blue), each shown separately or in an overlay (rightmost). The scale bars represent: **a-d.** 20 µm and **e.** 50 µm.

While positive immunostaining of metabolic enzymes was detected in the nucleus, it was not always equally compelling for every enzyme or in every cell type tested, thus, prompting further validation. Furthermore, it remained necessary to eliminate the potential non-specific staining. To address these issues, we generated C-termini tGFP-fusion of ACAA2, OGDH, and IDH2. In addition, we identified putative nuclear localization signals (NLS), of which we mutated those within ACAA2 and OGDH (supplementary Fig. 3S). These constructs would allow us to determine: 1) if the enzymes localize to the nucleus, 2) if its fusion to a cytosolic protein (tGFP) confers nuclear localization, 3) if the mutation of a putative NLS can reverse the localization, and 4) confirm localization with antibodies against the enzyme and the tGFP tags in the subcellular protein fractions. The constructs were delivered to cultured cardiac cells or human cancer cells via adenoviral vectors. Figure 3 shows that while tGFP was predominantly located in the cytosol and to a minimal extent in the mitochondrial/membrane (mito/mem) fraction, its fusion with ACAA2, OGDH, or IDH2, resulted in its redistribution, to include the nucleus and chromatin-bound protein fractions (Fig. 3a-c, upper 2 panels). Likewise, the endogenous enzymes were detected in the mito/mem fraction, which contains the mitochondria, as confirmed by VDAC1, in addition to the nuclear and chromatin fractions, as confirmed by TFIIB and histone H3 (Fig. 3a-c, second panels). Ultimately, substitutions of key lysine residues with glutamine in the putative NLS of ACCA2 (mtACAA2) or OGDH (mtOGDH) significantly reduced their nuclear import and chromatin associations, proving that this localization is specific and requires an NLS (Fig. 3a-c, 3d-i). Note, the chromatin bound proteins were not subjected to crosslinking, thus, only the directly- or tightly-associated proteins are retained in this fraction, such as the histones and RNA polymerase II. The ratios of ACAA2-tGFP in the different fractions show that % of total wild vs. NLS-mutant protein is lower in the cytosol and mitochondria, and higher in the nuclear and chromatin fractions as the translocation of the mutant to the nucleus is reduced in the latter (Fig. 3a, 3d-f). On the other hand, the mtOGDH-tGFP protein was predominantly in the mitochondrial, while minimally detected in the nucleus vs. the wild type OGDH-tGFP (Fig. 3b, 3g-h). Similar results were observed in human colon cancer cells (supplementary Fig. 4S). Interestingly, the levels of the ACAA2-tGFP protein were increased when the cells were incubated with palmitate vs. glucose, particularly in the nucleus and chromatin-bound fractions (supplementary Fig. 4S). Additionally, the metabolic enzymes were also detected in the nucleus of the mouse heart tissue and isolated myocytes, although there was no significant difference when growth-induced with pressure overload or endothelin-1 (supplementary Fig. 5S). These data confirm that mitochondrial enzymes reside in the nucleus in significant concentrations and that at least ACAA2 and OGDH harbor NLSs that mediate their nuclear import. The data also suggest that nuclear and chromatin enzymes are limiting and subject to regulation by metabolic substrates.

**Figure 3.**
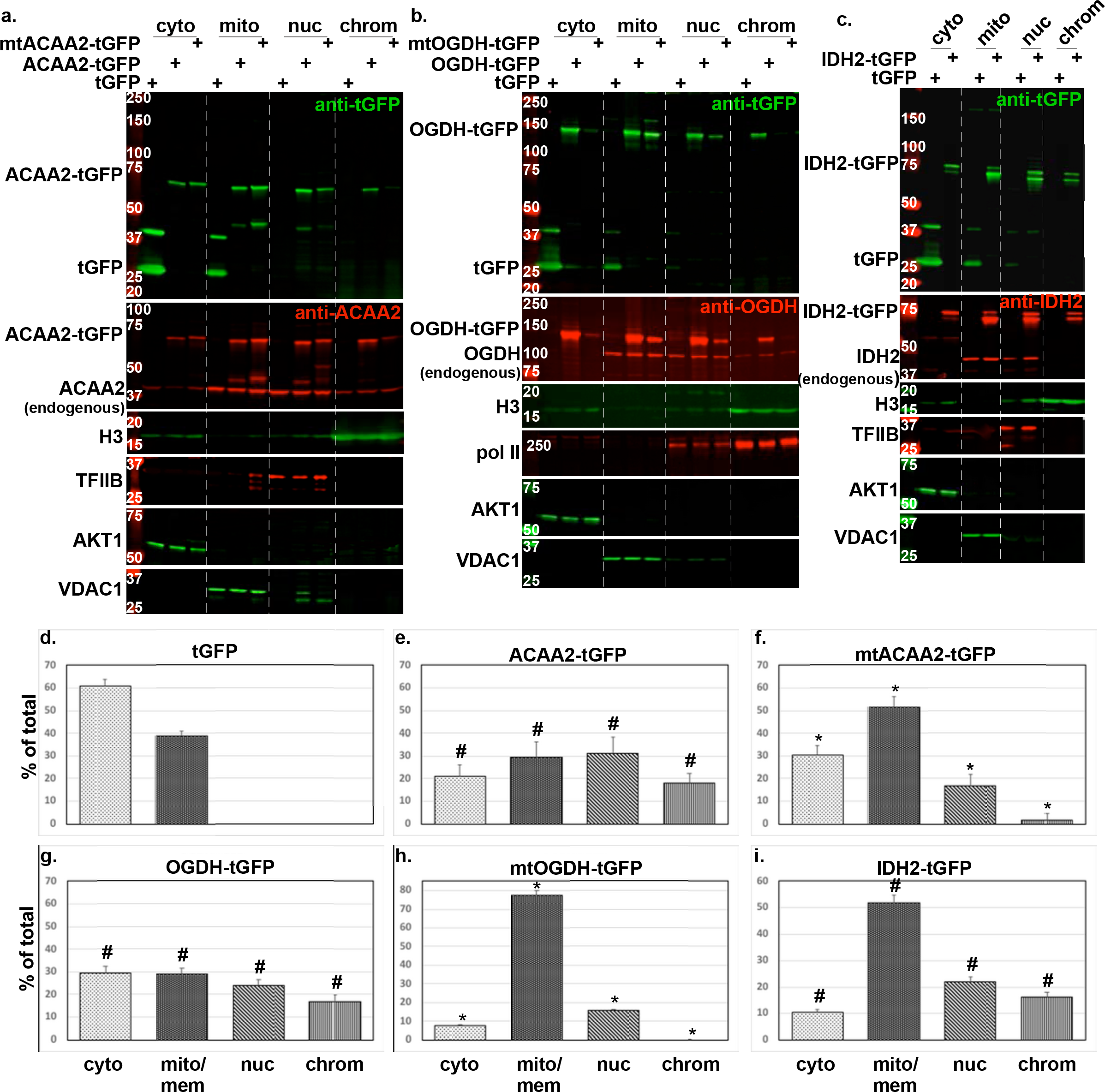
Nuclear localization of metabolic enzymes confirmed by tGFP-fusion proteins and NLS mutations. Cardiac myocytes were infected with a 10-20 moi of recombinant adenoviruses harboring turbo-GFP (tGFP) or **a.** wt ACAA-tGFP or an NLS mutant (mtACAA2-tGFP), **b.** wt OGDH or an NLS mutant (mtOGDH-tGFP), **c.** wt IDH2-tGFP, tGFP-fused cDNAs. After 18 h, the cellular protein/organelles were fractionated into cytosol (cyto), mitochondrial and membrane (mito), nuclear (nuc), and chromatin-bound (chrom) protein fractions that were then analyzed by Western blotting for the proteins listed on the left of each panel. The fusion proteins were detected by anti-GFP (upper panels, a-c) and anti-ACAA2, anti-OGDH, and anti-IDH2 (second panels, a-c), which also detect the endogenous proteins. AKT1, VDAC1, TFIIB or Pol II, and H3, were immunodetected for use as internal controls for the corresponding cell fractions. The signals for the **d.** tGFP, and tGFP-fusion proteins **e.** ACAA2-tGFP, **f.** mtACAA2-tGFP, **g.** OGDH-tGFP, **h.** mtOGDH-tGFP, **i.** IDH2-tGFP (top panels), were quantified using imageJ, normalized to internal controls, and plotted as the mean ± SEM of % total protein in all 4 fractions. Error bars represent SEM, N=3 from 3 repeats. \**p* = 0.0095 vs. wt tGFP-fusion, #*p* ≤ 0.05 vs. tGFP, in corresponding fractions.

**Figure 4.**
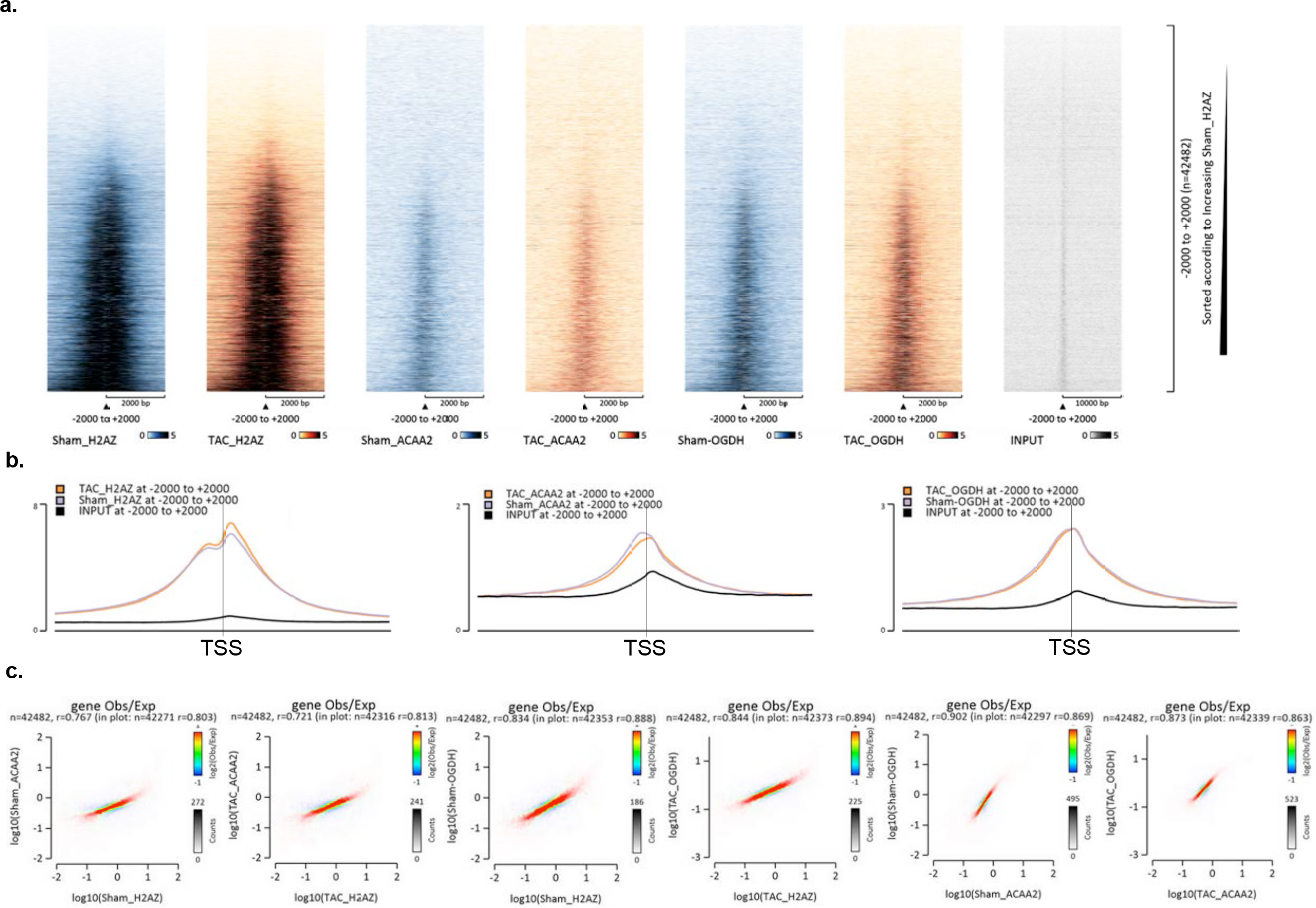
The association of ACAA2 and OGDH with chromatin overlaps with H2A.Z at transcription start sites. Mice were subjected to a sham or TAC operation. One-week post-TAC, the hearts were isolated and analyzed by ChIP-Seq for H2A.Z, ACAA2, or OGDH (pool of 3, each). The sample size was determined based on the quantity of the material required for the assay and for biological averaging of samples based on our prior experience of the variability between mouse hearts in response to growth induction. **a.** Heatmaps of the ChIP-Seq sequence Tags from sham (blue), TAC (brown), and input (grey) aligned at the TSS. The y-axis represents individual positions of bins, and the x-axis represents a region from −2000 to +2000 bp relative to the TSS. **b.** Graphs representing average peak values of H2A.Z, ACAA2, and OGDH ChIP-Seq Tags from sham, TAC, and input from −2000 to +2000 bp relative to the TSS. **c.** Histograms showing the distribution of fragments calculated from their overall frequencies in the ChIP-Seq of H2A.Z (X-axis) v. ACAA2 or OGDH (Y-axis), and of ACAA2 (X-axis) v. OGDH (Y-axis), over the length of the gene and including −2000 bp upstream of the TSS, as labeled. The x- and y-axes were segmented into 75 bins, and the number of fragments within each bin was counted, color coded, and plotted. The bar to the right of the plot illustrates the relationship between count and coloring. The plots represent pseudo-colored 2D matrices showing observed/expected distribution, calculated from the overall frequencies of fragments on each of the axes. This plot shows the relation between H2A.Z and ACAA2 or OGDH, and of ACAA2 and OGDH levels, relative to what is expected if they occurred by chance. The pseudo-color corresponds to the Obs/Exp ratio, and the color intensity is proportional to the log2 of the number of observed fragments within each bin. These plots suggest that there is a positive correlation between the levels of H2A.Z and ACAA2 or OGDH, where the red indicates that this occurs more frequently than expected by chance, as denoted by the correlation coefficient listed above each plot. For the separate observed and expected histograms, please see supplementary Fig. 3S. The plots in this figure were generated by EaSeq software.

**Figure 5.**
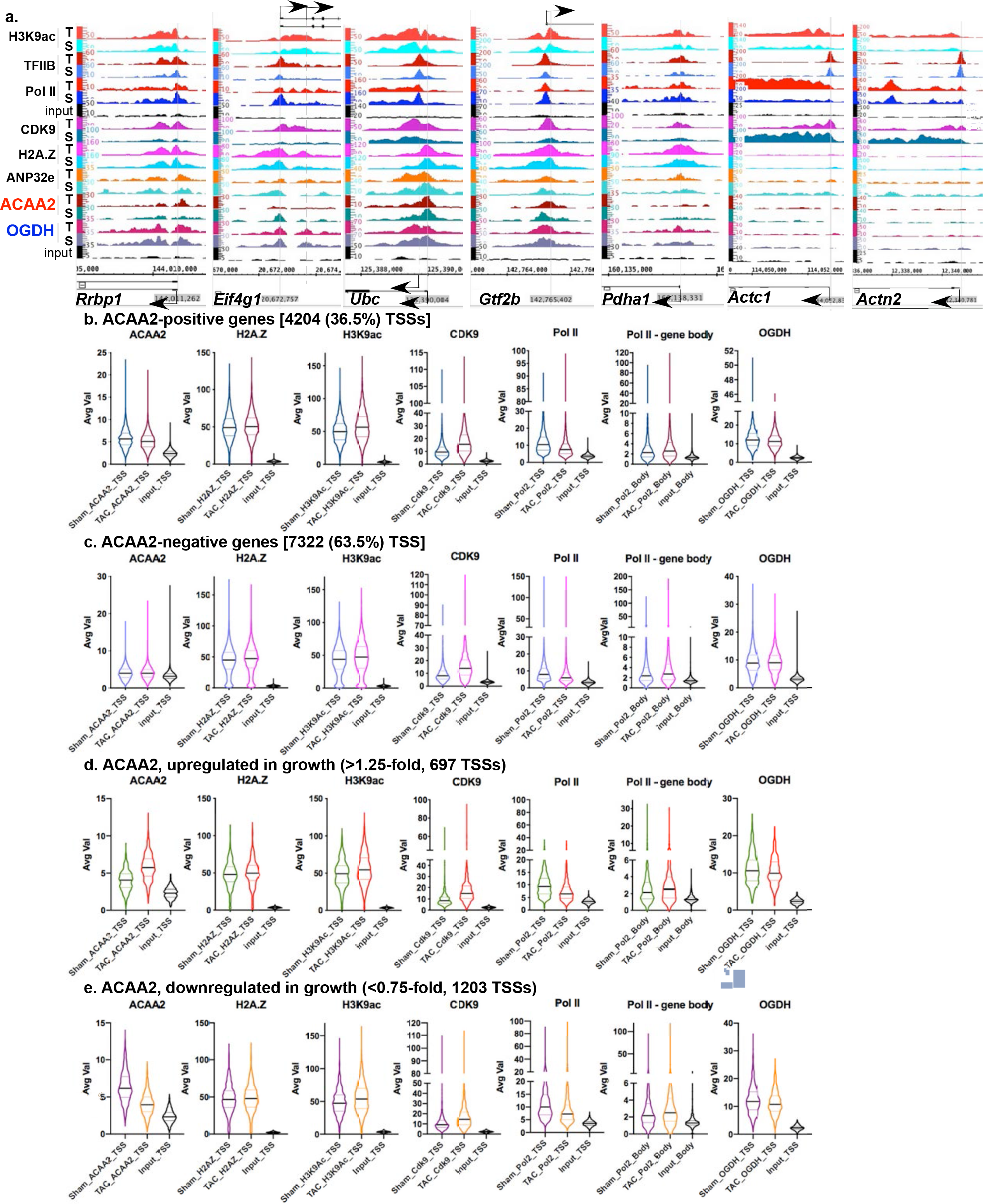
ACAA2 binds selectively to the TSS of genes and is differentially regulated during growth. Mice were subjected to a sham or TAC operation. One-week post-TAC, the hearts were isolated and analyzed by ChIP-Seq for ACAA2. **a.** The alignment of the ChIP-Seq sequence tags for H3K9ac, TFIIB, pol II, Cdk9, H2A.Z, ANP32E, ACAA2, and OGDH (Y-axis) across the genome’s coordinates (X-axis) of the TSS regions of *Rrbp1*, *Eif4g1*, *Ubc*, *Gtf2b*, *Pdha1, Actc1, and Actn2* genes. The arrow shows the start and direction of transcription. Expressed genes (RNA pol II positive) were sorted into 4 groups: **b.** those that bind ACAA2 (ACAA2-positive), **c.** ACAA2-negative, **d.** ACAA2-positive that exhibit upregulation during growth, and **e.** ACAA2-positive that exhibit downregulation during growth. The ChIP-Seq sequence Tags for ACAA2, H2A.Z, H3K9ac, OGDH, and pol II, at the TSS (−1000 to +1000), from these groups, were plotted as violin plots, in which the horizontal solid line represent the median, and the dashed lines the quartiles, whereas, the shape of the violin reflect the tags’ density distribution.

### OGDH and ACAA2 associate with H2A.Z-bound transcription start sites

The co-precipitation of mitochondrial enzymes with chromatin-bound H2A.Z indicates that these enzymes associate with chromatin and co-localize with H2A.Z at TSSs. To confirm this, we performed a ChIP-Seq assay using anti-ACAA2 or anti-OGDH on chromatin extracted from normal vs. hypertrophied hearts (1 wk post-TAC). The ChIP-Seq statistics, including the number of tags, peaks, and empirical false discovery rates (FDR) are reported in supplementary Table 1, whereas, the raw (Fastq files) and aligned data (BigWig and Bam files) are deposited in Gene Expression Omnibus (GEO) datasets (accession pending). The results show that ACAA2 and OGDH, similar to H2A.Z ^17^, predominantly associate with TSSs, as observed with heatmaps of the sequence tags and the curves of the average signals aligned to a region encompassing −2000 to +2000 bp from the TSS, with no substantial differences observed between their total levels in the normal v. growth-induced hearts (Fig. 4a-b). Additionally, the binding of the ACAA2 and OGDH coincided with that of H2A.Z (when analyzed over the length of the gene and including −2000 bp upstream of the TSS) at values higher than expected for a random event (r=0.813, 0.803, 0.888, 0.894, in normal or growth-induced hearts for each gene, respectively (Fig. 4c and supplementary Fig. 6S). Additionally, we found that ACAA2’s and OGDH’s chromatin binding sites extensively overlap (r= 0.86, Fig. 4c). This supports our conclusion that H2A.Z is a major recruiter of metabolic enzymes.

**Table 1.**
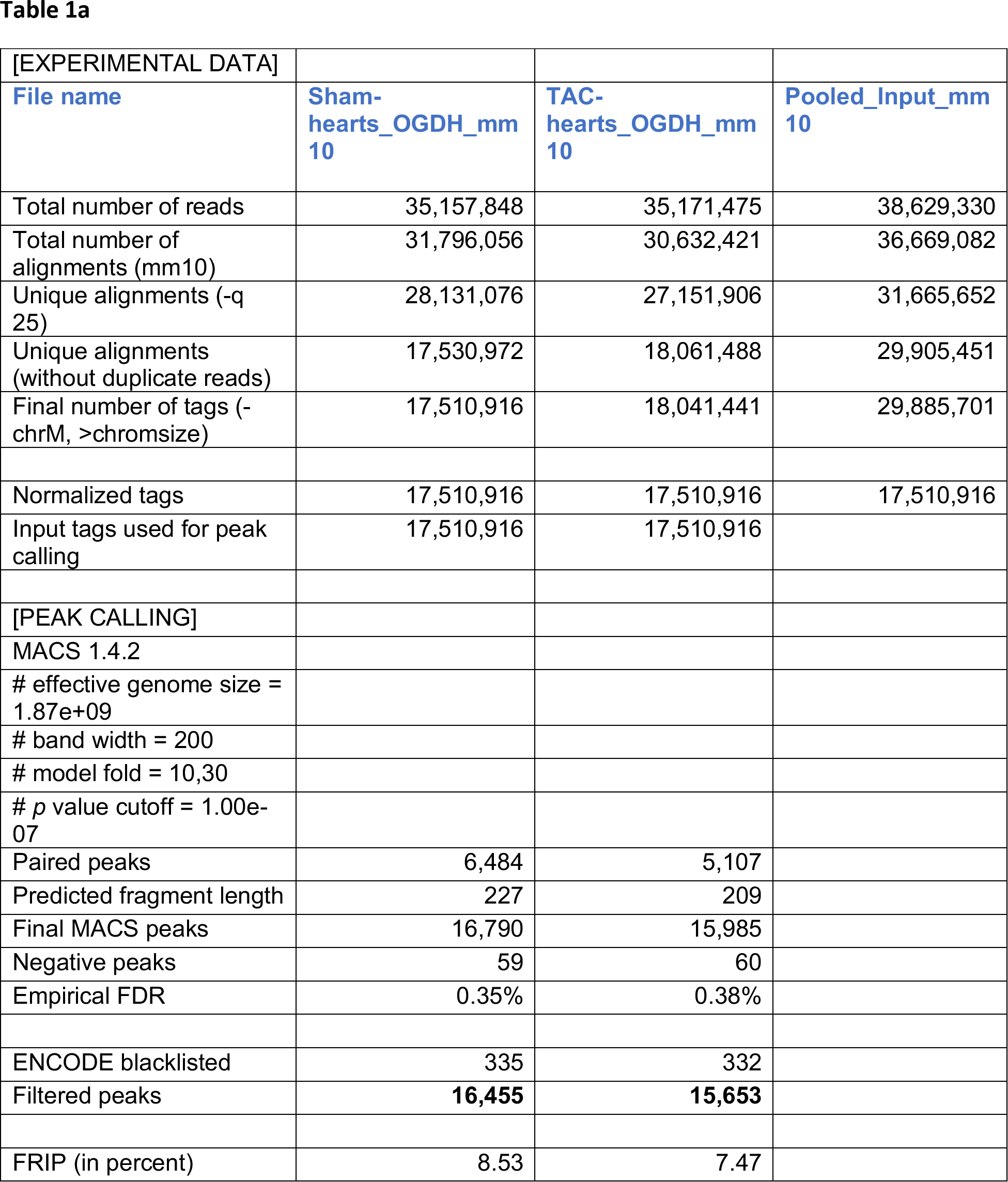
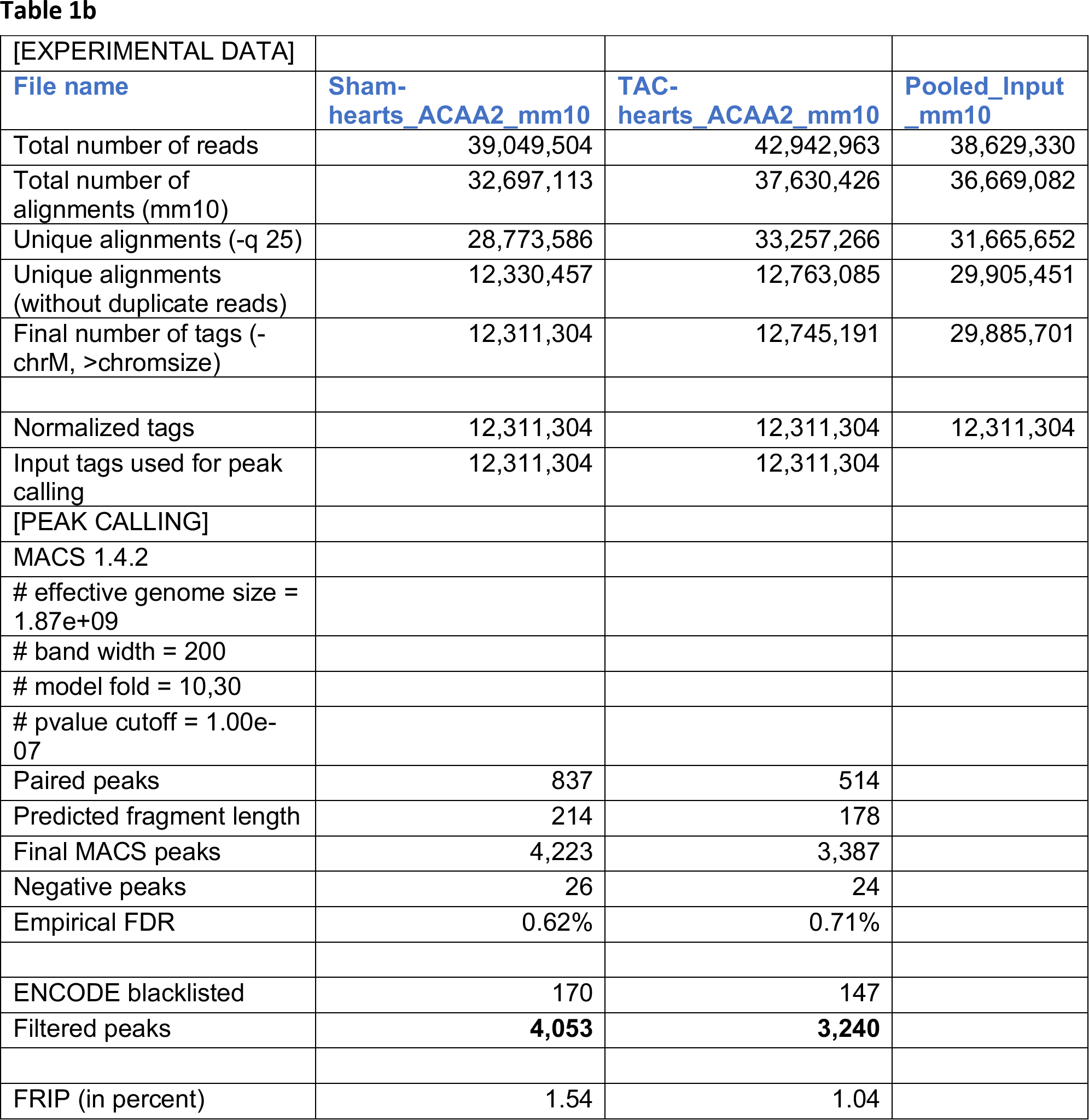
ChIP-Sequencing Statistics (by ActiveMotif) **Antibodies used:** anti-OGDH (Cell Signaling, #26865) and anti-ACAA2 (Origene, #TA506126) Samples: 1. Sham-operated, 12 wk-old, male, mouse heart tissue (n=3). 2. Transverse aortic constriction for 1 wk, 12 wk-old, male, mouse heart tissue (n=3).

| [EXPERIMENTAL DATA] |  |  |  |
| --- | --- | --- | --- |
| File name | Sham-<br>hearts_OGDH_mm<br>10 | TAC-hearts_OGDH_mm<br>10 | Pooled_Input_mm<br>10 |
| Total number of reads | 35,157,848 | 35,171,475 | 38,629,330 |
| Total number of alignments (mm10) | 31,796,056 | 30,632,421 | 36,669,082 |
| Unique alignments (-q 25) | 28,131,076 | 27,151,906 | 31,665,652 |
| Unique alignments (without duplicate reads) | 17,530,972 | 18,061,488 | 29,905,451 |
| Final number of tags (-chrM, >chromsize) | 17,510,916 | 18,041,441 | 29,885,701 |
| Normalized tags | 17,510,916 | 17,510,916 | 17,510,916 |
| Input tags used for peak calling | 17,510,916 | 17,510,916 |  |
| [PEAK CALLING] |  |  |  |
| MACS 1.4.2 |  |  |  |
| # effective genome size = 1.87e+09 |  |  |  |
| # band width = 200 |  |  |  |
| # model fold = 10,30 |  |  |  |
| # p value cutoff = 1.00e-07 |  |  |  |
| Paired peaks | 6,484 | 5,107 |  |
| Predicted fragment length | 227 | 209 |  |
| Final MACS peaks | 16,790 | 15,985 |  |
| Negative peaks | 59 | 60 |  |
| Empirical FDR | 0.35% | 0.38% |  |
| ENCODE blacklisted | 335 | 332 |  |
| Filtered peaks | **16,455** | **15,653** |  |
| FRIP (in percent) | 8.53 | 7.47 |  |
| [EXPERIMENTAL DATA] |  |  |  |
| **File name** | **Sham-**<br>**hearts_ACAA2_mm10** | **TAC-**<br>**hearts_ACAA2_mm10** | **Pooled_Input**<br>**_mm10** |
| Total number of reads | 39,049,504 | 42,942,963 | 38,629,330 |
| Total number of alignments (mm10) | 32,697,113 | 37,630,426 | 36,669,082 |
| Unique alignments (-q 25) | 28,773,586 | 33,257,266 | 31,665,652 |
| Unique alignments (without duplicate reads) | 12,330,457 | 12,763,085 | 29,905,451 |
| Final number of tags (-chrM, >chromsize) | 12,311,304 | 12,745,191 | 29,885,701 |
| Normalized tags | 12,311,304 | 12,311,304 | 12,311,304 |
| Input tags used for peak calling | 12,311,304 | 12,311,304 |  |
| [PEAK CALLING] |  |  |  |
| MACS 1.4.2 |  |  |  |
| # effective genome size = 1.87e+09 |  |  |  |
| # band width = 200 |  |  |  |
| # model fold = 10,30 |  |  |  |
| # pvalue cutoff = 1.00e-07 |  |  |  |
| Paired peaks | 837 | 514 |  |
| Predicted fragment length | 214 | 178 |  |
| Final MACS peaks | 4,223 | 3,387 |  |
| Negative peaks | 26 | 24 |  |
| Empirical FDR | 0.62% | 0.71% |  |
| ENCODE blacklisted | 170 | 147 |  |
| Filtered peaks | **4,053** | **3,240** |  |
| FRIP (in percent) | 1.54 | 1.04 |  |

**Figure 6.**
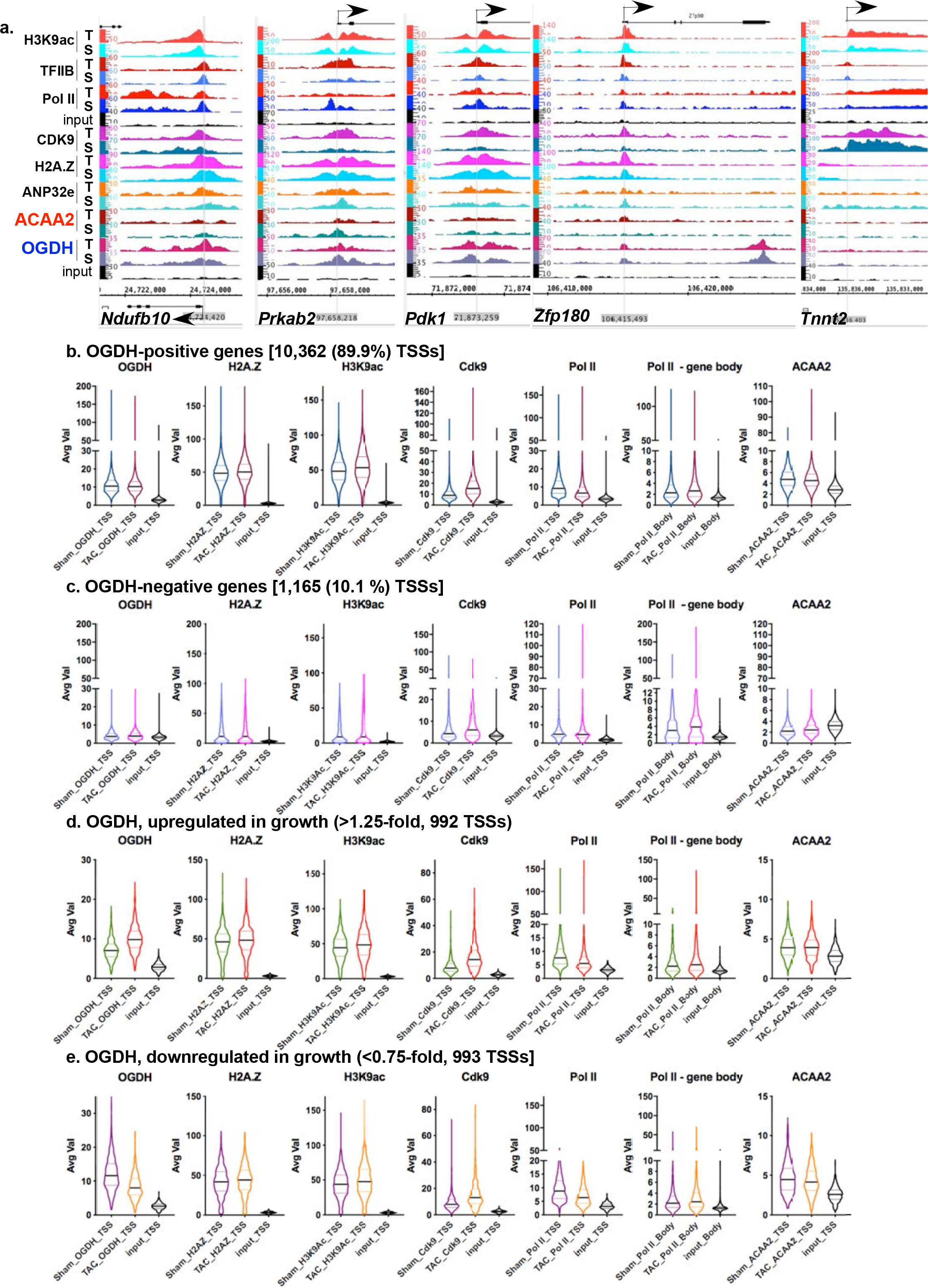
OGDH binds to the TSSs of 89.9% of expressed genes in consistent overlap with H2A.Z. Mice were subjected to a sham or TAC operation. One-week post-TAC, the hearts were isolated and analyzed by ChIP-Seq for OGDH. **a.** The alignment of the sequence tags of H3K9ac, TFIIB, pol II, Cdk9, H2A.Z, ANP32E, ACAA2, and OGDH (Y-axis) across the genome’s coordinates (X-axis) of the TSS regions of *Ndufb10*, *Prkab2*, *Pdk1*, *Zfp180*, and *Tnnt2*. The arrow shows the start and direction of transcription. Expressed genes (RNA pol II positive) were sorted into 4 groups: **b.** those that bind OGDH (OGDH-positive), **c.** OGDH-negative, **d.** OGDH-positive that exhibit upregulation during growth, and **e.** OGDH-positive that exhibit downregulation during growth. The sequence tags for OGDH, H2A.Z, H3K9ac, ACAA2, and pol II, at the TSS (−1000 to +1000), from these groups, were plotted as violin plots, in which the horizontal solid line represent the median, and the dashed lines the quartiles, whereas, the shape of the violin reflect the tags’ density distribution.

### ACAA2 selectively and exclusively associates with H2A.Z-bound TSS

The Avg Val of the sequence tags from the ACAA2 ChIP-Seq analysis were sorted into ACAA2-positve and -negative TSSs (−1000 to +1000 bp from TSS) for transcriptionally active genes (determined by RNA pol II binding), in parallel with those of H2A.Z, H3K9ac, Cdk9, and RNA pol II ChIP-Seq data. The data were graphed as violin plots representing the median, quartiles, and distribution and probability density of the tags. This revealed that ACAA2 associates with the TSS of 4204 genes (36.5% of genes expressed in the heart) that are also all H2A.Z-bound (approx. 90% of expressed genes are H2A.Z-bound, see Fig. 5a-c). Conversely, not all H2A.Z-bound genes were associated with ACAA2, suggesting selectivity and the involvement of other regulators (Fig. 5a-c). Notably, ACAA2-positive TSSs exhibit a higher median for bound H2A.Z (8% and 6% higher for normal and growth-induced hearts, respectively) and the H3K9ac mark (13.6% and 12% higher for normal and growth-induced hearts, respectively) relative to those negative for ACAA2, as reflected in the violin plots (Fig. 5b-c). Interestingly, both H2A.Z and H3K9ac show similar patterns of tag density distribution, as it is altered in the absence vs. presence of ACAA2, demonstrating a positive correlation between the two marks. ACAA2-positive TSSs also exhibit significantly higher levels of pol II and Cdk9 peaks, denoting higher transcriptional activity (supplementary Fig. 7S). On the other hand, there is no correlation between changes in ACAA2 abundance during cardiac growth with the upregulation or downregulation of Cdk9, H3K9ac, or pol II (Fig. 5d-e). Thus, although ACAA2 preferentially associates with transcriptionally active genes (with the exception of cardiac-specific genes), changes in its abundance does not correlate with changes in transcriptional activity. Of the 4204 genes, 697 exhibited ≥ 1.25-fold upregulation of ACAA2, while 1203 genes exhibited ≤ 0.75-fold downregulation, and 1900 genes with minimal or no change of ACAA2, in growth-induced v. normal hearts. Broadly, functional pathway analysis shows that these three categories of ACAA2-bound genes encompass pathways involved in endoplasmic protein processing and proteolysis, metabolism, and RNA transport and protein synthesis, respectively (supplementary Tables 2-4S).

**Table 2.**
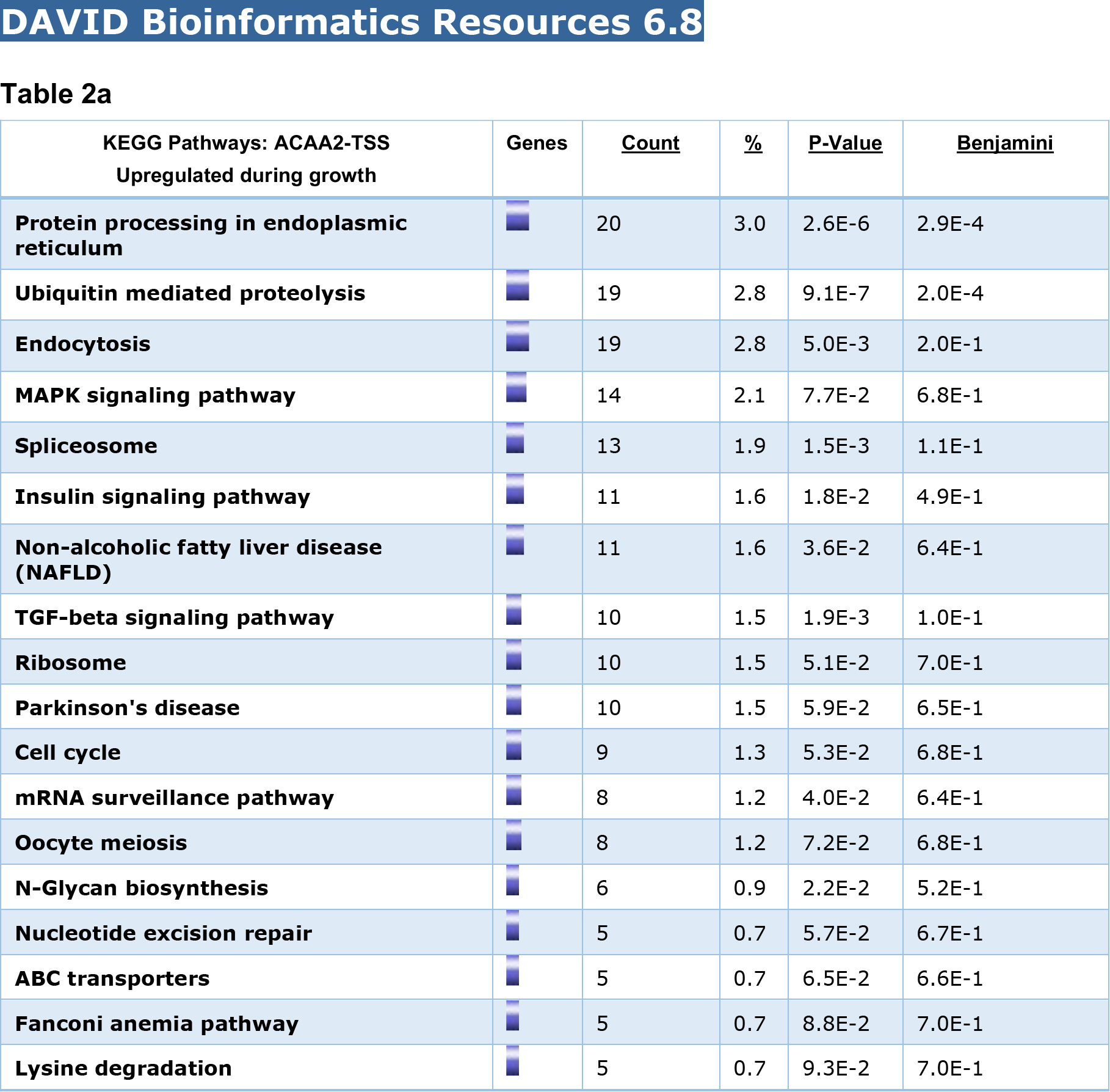
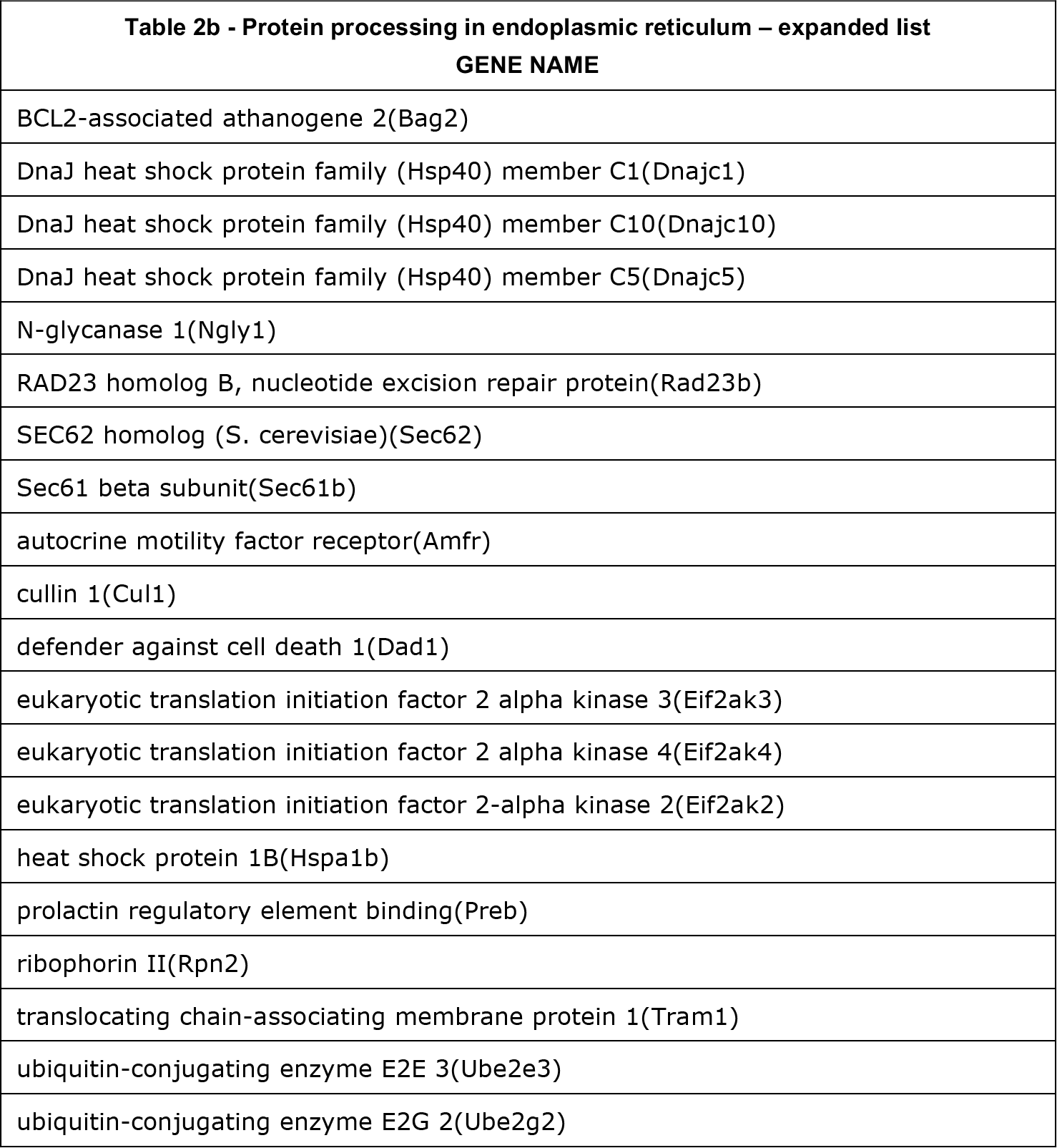
KEGG functional pathway analysis of genes that exhibit upregulation of ACAA2 at the TSS during TAC (697 genes)

| KEGG Pathways: ACAA2-TSS<br>Upregulated during growth | Genes | <u>Count</u> | <u>%</u> | <u>P-Value</u> | <u>Benjamini</u> |
| --- | --- | --- | --- | --- | --- |
| **Protein processing in endoplasmic reticulum** |  | 20 | 3.0 | 2.6E-6 | 2.9E-4 |
| **Ubiquitin mediated proteolysis** |  | 19 | 2.8 | 9.1E-7 | 2.0E-4 |
| **Endocytosis** |  | 19 | 2.8 | 5.0E-3 | 2.0E-1 |
| **MAPK signaling pathway** |  | 14 | 2.1 | 7.7E-2 | 6.8E-1 |
| **Spliceosome** |  | 13 | 1.9 | 1.5E-3 | 1.1E-1 |
| **Insulin signaling pathway** |  | 11 | 1.6 | 1.8E-2 | 4.9E-1 |
| **Non-alcoholic fatty liver disease (NAFLD)** |  | 11 | 1.6 | 3.6E-2 | 6.4E-1 |
| **TGF-beta signaling pathway** |  | 10 | 1.5 | 1.9E-3 | 1.0E-1 |
| **Ribosome** |  | 10 | 1.5 | 5.1E-2 | 7.0E-1 |
| **Parkinson's disease** |  | 10 | 1.5 | 5.9E-2 | 6.5E-1 |
| **Cell cycle** |  | 9 | 1.3 | 5.3E-2 | 6.8E-1 |
| **mRNA surveillance pathway** |  | 8 | 1.2 | 4.0E-2 | 6.4E-1 |
| **Oocyte meiosis** |  | 8 | 1.2 | 7.2E-2 | 6.8E-1 |
| **N-Glycan biosynthesis** |  | 6 | 0.9 | 2.2E-2 | 5.2E-1 |
| **Nucleotide excision repair** |  | 5 | 0.7 | 5.7E-2 | 6.7E-1 |
| **ABC transporters** |  | 5 | 0.7 | 6.5E-2 | 6.6E-1 |
| **Fanconi anemia pathway** |  | 5 | 0.7 | 8.8E-2 | 7.0E-1 |
| **Lysine degradation** |  | 5 | 0.7 | 9.3E-2 | 7.0E-1 |

| **Table 2b - Protein processing in endoplasmic reticulum – expanded list**<br>**GENE NAME** |
| --- |
| BCL2-associated athanogene 2(Bag2) |
| DnaJ heat shock protein family (Hsp40) member C1(Dnajc1) |
| DnaJ heat shock protein family (Hsp40) member C10(Dnajc10) |
| DnaJ heat shock protein family (Hsp40) member C5(Dnajc5) |
| N-glycanase 1(Ngly1) |
| RAD23 homolog B, nucleotide excision repair protein(Rad23b) |
| SEC62 homolog (S. cerevisiae)(Sec62) |
| Sec61 beta subunit(Sec61b) |
| autocrine motility factor receptor(Amfr) |
| cullin 1(Cul1) |
| defender against cell death 1(Dad1) |
| eukaryotic translation initiation factor 2 alpha kinase 3(Eif2ak3) |
| eukaryotic translation initiation factor 2 alpha kinase 4(Eif2ak4) |
| eukaryotic translation initiation factor 2-alpha kinase 2(Eif2ak2) |
| heat shock protein 1B(Hspa1b) |
| prolactin regulatory element binding(Preb) |
| ribophorin II(Rpn2) |
| translocating chain-associating membrane protein 1(Tram1) |
| ubiquitin-conjugating enzyme E2E 3(Ube2e3) |
| ubiquitin-conjugating enzyme E2G 2(Ube2g2) |

**Table 3.**
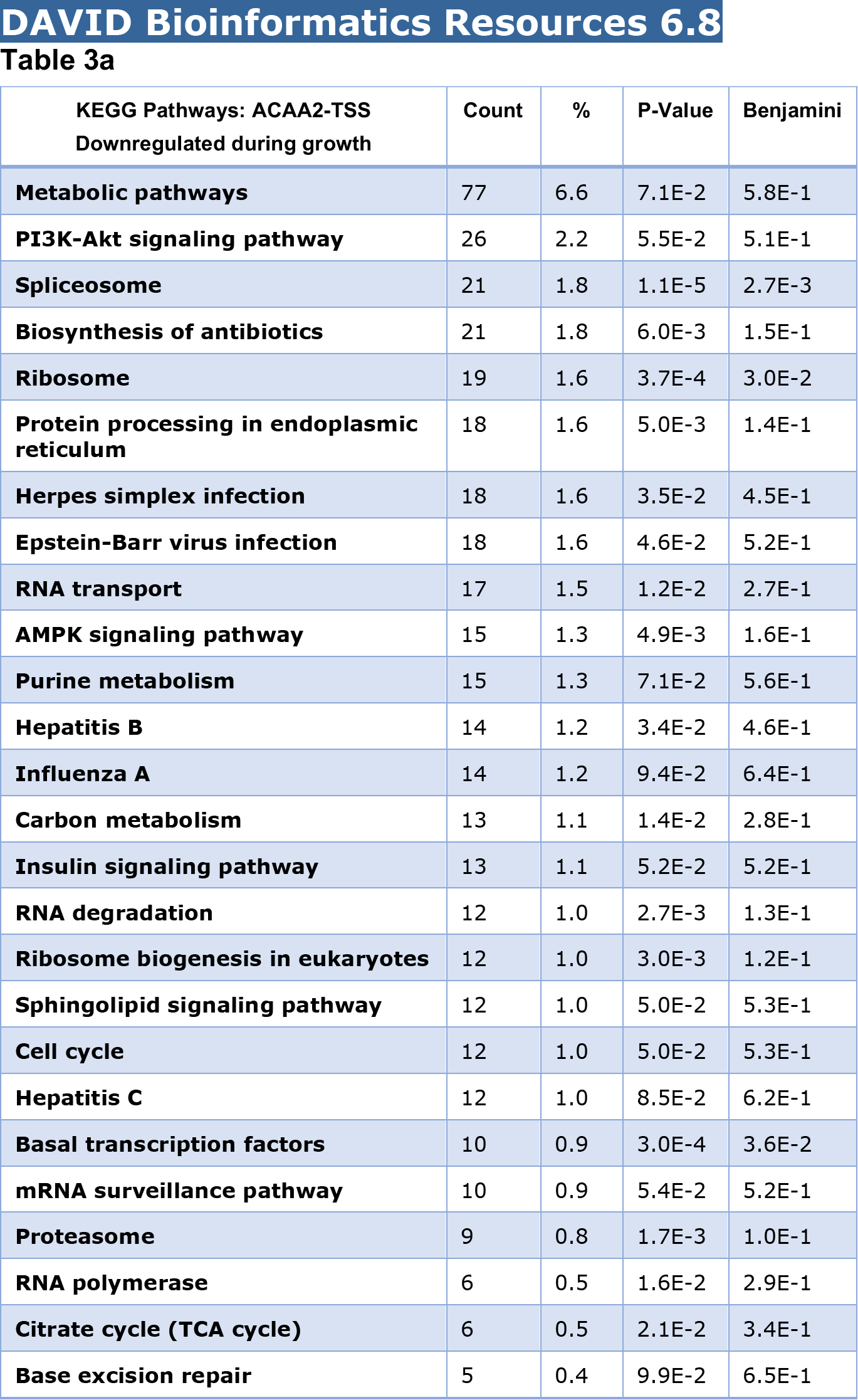
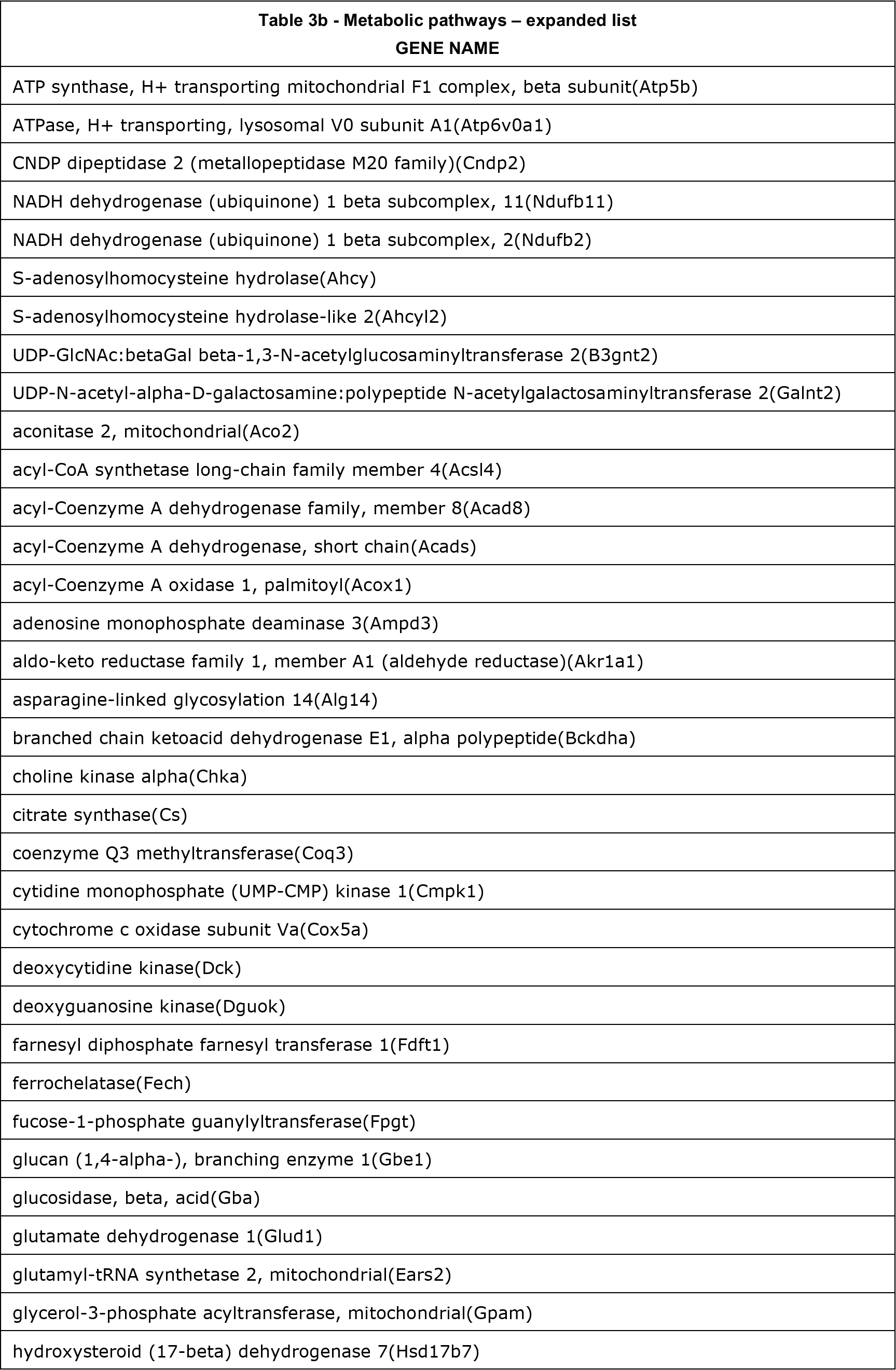
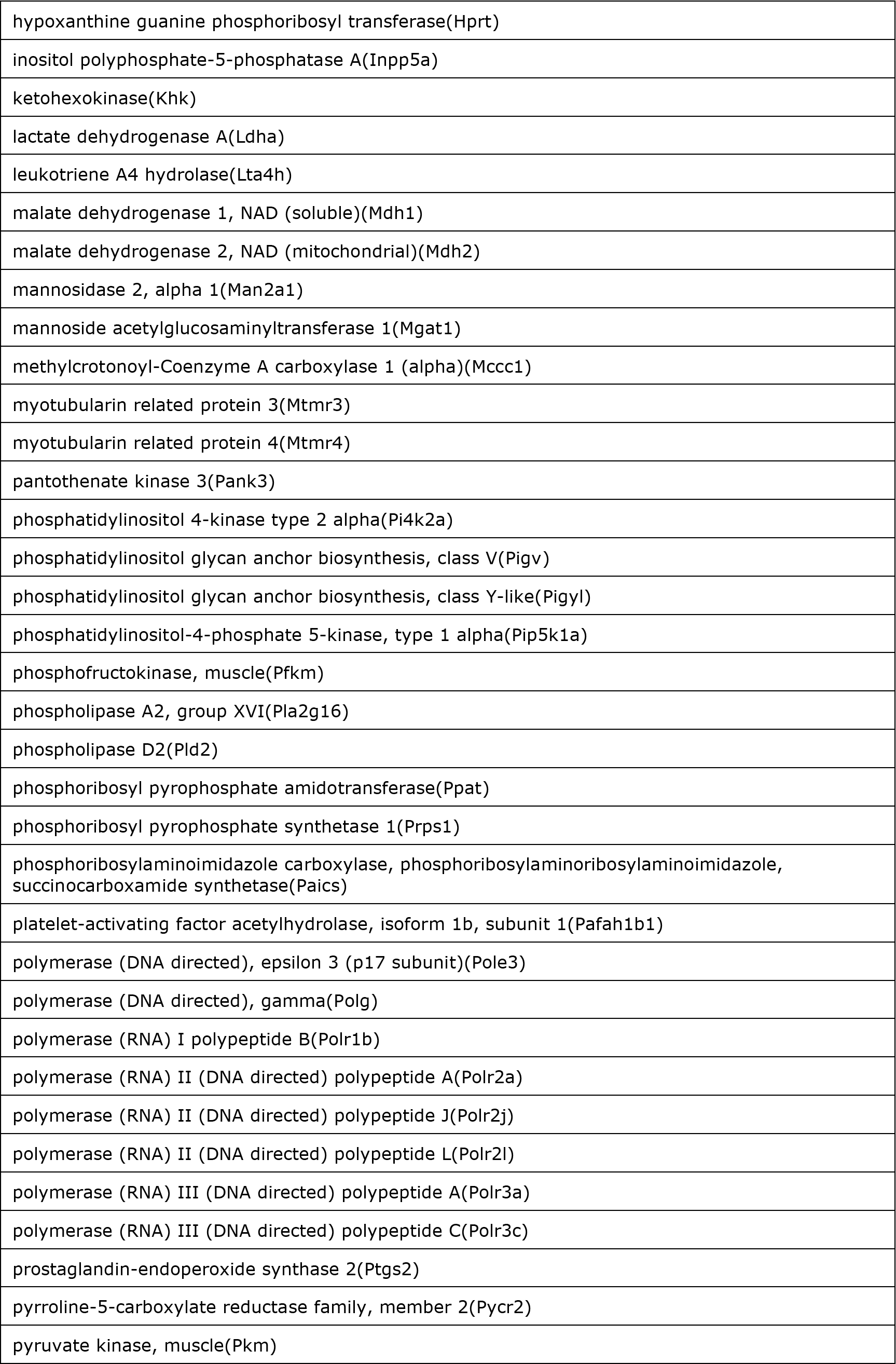
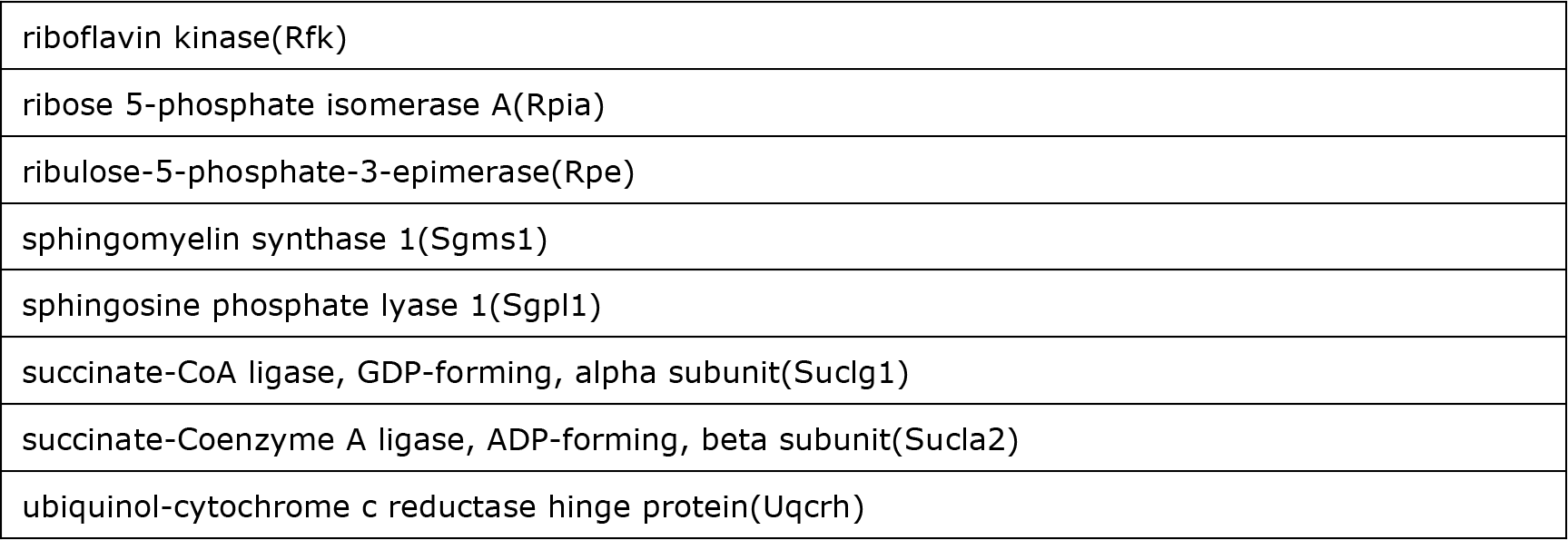
KEGG functional pathway analysis of genes that exhibit downregulation of ACAA2 at the TSS during TAC (1203 genes)

**Figure 7.**
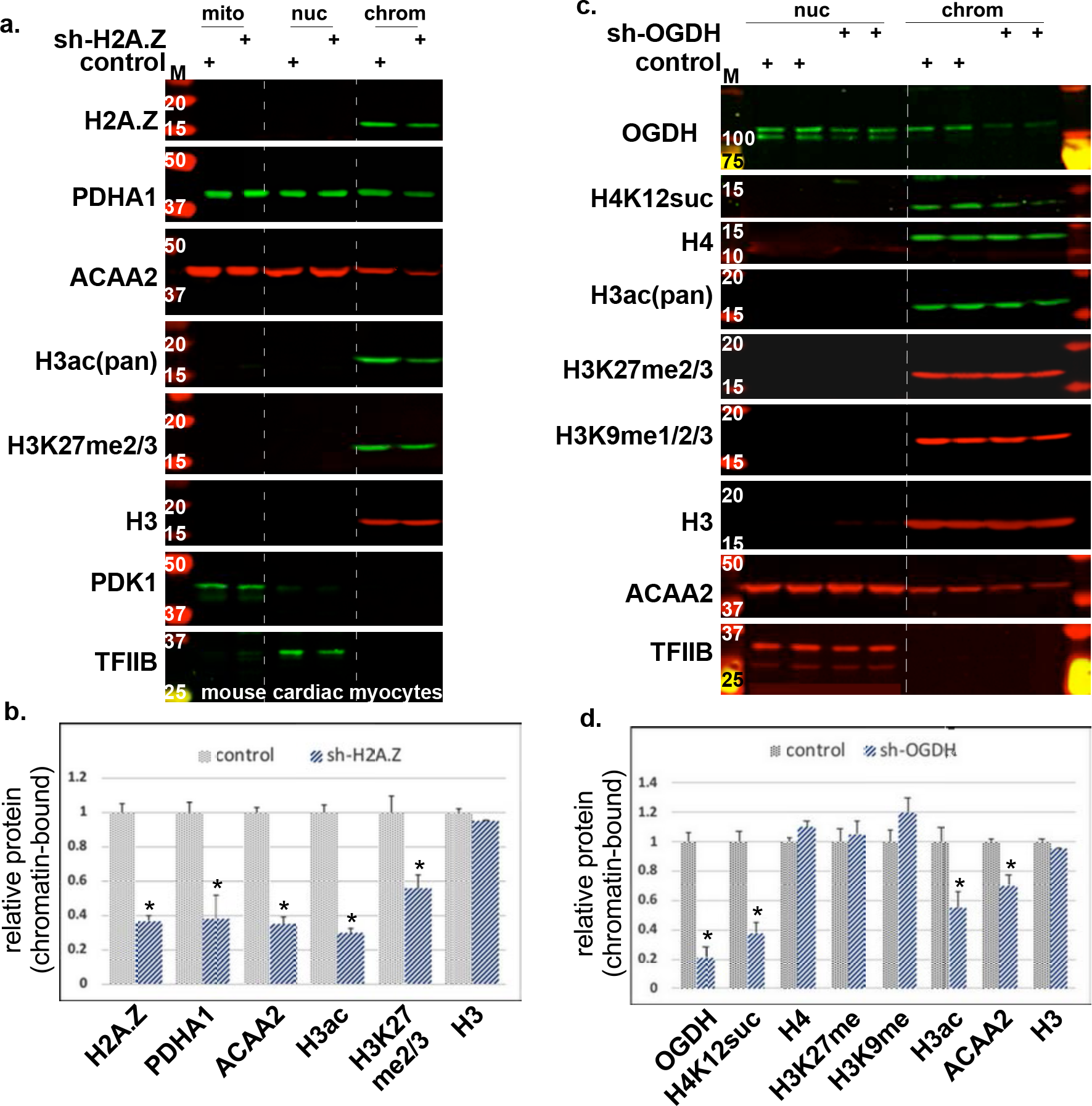
Knockdown of H2A.Z in rodent cells inhibits chromatin binding of metabolic enzymes and reduces histone modifications. Mouse adult cardiac myocytes were isolated from the hearts of 8 wk old male C57/Bl mice. They were the infected with 30 moi of adenoviruses harboring a nonsense shRNA control or one targeting H2A.Z. After 24 h, organelles were isolated and fractionated into membrane/mitochondrial (Mem), nuclear (Nuc), and chromatin-bound (Chrom, no crosslinking applied), using a combination of differential lysis and sequential centrifugation. The proteins extracted from each of these fractions were analyzed by Western blotting for the genes indicated on the right of each panel. **b.** The Western blot signals (n=3, each, from 3 repeats), was quantitated and plotted as the mean ± SEM relative to the control, adjusted to 1. Error bars represent standard error of the mean, and \**p* = 0.03 v. each’s corresponding control. **c.** shRNA targeting OGDH or a control construct, was delivered to isolated rat neonatal cardiac myocytes using adenoviral vectors (moi 30). After 48 h, organelles were isolated, fractionated, and analyzed as described in (a). The Western blot signals (n=4, each, from 4 repeats), was quantitated and plotted as the mean ± SEM relative to the control, adjusted to Error bars represent standard error of the mean, and \**p* = 0.035 v. each’s corresponding control.

The sequence tags of the ACAA2 ChIP-Seq were also aligned with those of H3K9ac, a histone mark that is associated with active promoters; TFIIB, which demarcates the TSS; RNA pol II, which reflects transcriptional activity; CDK9, which reflect transcriptional elongation; and ANP32E, which is a known H2A.Z-interacting protein. Figure 5a shows the changes in peak densities of these molecules in the normal v. growth-induced hearts, across the chromosomal coordinates of the TSS of *Rrbp1*, a ribosome binding protein; *Eif4g1*, which is involved in translation initiation; *Gtf2b*, required for initiation of transcription; and *Ubc*, a substrate for protein ubiquitination. These represent genes that exhibited an increase or a decrease in ACAA2 abundance in growth-induced v. normal hearts (e.g. *Rrbp1 and Eif4g2,* respectively) or remain unchanged (e.g. *Gtf2b* and *Ubc*). Notably, these genes contrasted with all cardiac-specific genes (shown are *Actc1* and *Actn2*), which have no detectable ACAA2, coinciding with the lack of, or undetectable, H2A.Z ^17^ (Fig. 5a). Therefore, these data reveal, for the first time, the nuclear localization and chromatin binding of a beta-oxidation enzyme and validate our RIME analysis. Also, consistent with our H2A.Z-RIME, ACAA2 associated exclusively with H2A.Z-bound TSSs, providing support of specificity for this association.

### OGDH exclusively associates with all H2A.Z-bound TSS

The Avg Val of the sequence tags from the OGDH ChIP-Seq analysis were sorted into OGDH-positive and -negative TSSs (−1000 to +1000 bp from TSS) of transcriptionally active genes (determined by RNA pol II binding), in parallel with those of H2A.Z, H3K9ac, Cdk9, and RNA pol II ChIP-Seq data. The data were graphed as violin plots representing the median, quartiles, and distribution and probability density of the tags. This analysis revealed that OGDH preferentially associates with H2A.Z-bound TSSs with substantially higher H2A.Z densities (4.3- and 4.5-fold higher medians in sham and TAC hearts, respectively, vs. OGDH-negative genes), which includes 89.9% (10,362) of expressed genes (Fig. 6a-c). This also coincides with substantially higher levels of H3K9ac (5.3- and 6.5-fold higher medians for sham and TAC hearts, respectively, vs. OGDH-negative genes) and Cdk9 (2.25- and 2.5-fold higher medians for sham and TAC hearts, respectively, vs. OGDH-negative genes). With regards to pol II, the reduction in paused TSS-pol II that is associated with an increase in TSS-Cdk9 and incremental increase in gene body-pol II, in growth-induced vs. normal hearts, which is characteristic of a pause-release in transcription, was uniquely observed in OGDH-positive TSSs (Fig. 6b and supplementary 7S-c).

Of the 10,362 OGDH-positive genes, 992 exhibited ≥1.25-fold increase of OGDH, while 993 exhibited ≤ 0.75-fold downregulation in OGDH at the TSSs, in growth-induced v. normal hearts (Fig. 6d-e). Similar to ACCA2, there is no correlation of the changes in OGDH abundance with the those observed in Cdk9, H3K9ac, or pol II in normal v. growth-induced hearts, and, thus, transcriptional activity (Fig. 6d-e). The sequence tags of the OGDH ChIP-Seq were also aligned with those of H3K9ac, TFIIB, RNA pol II, Cdk9, ANP32E, and ACAA2 across the genome (Fig. 6a). Figure 6a shows the changes in peak densities in the normal v. growth-induced hearts, across the chromosomal coordinates of TSS regions of *Ndufb10*, which exhibits an increase, *Prkab2*, a decrease, and *Pdk1* no changes in OGDH binding across the TSSs upon growth induction, as examples. Broadly, functional pathway analyses show that genes that exhibit upregulation of OGDH during cardiac growth include a preponderance of metabolic genes, while those that show downregulation include pathways in cancer and endocytosis (supplementary Tables 5-6). In contrast to OGDH-positive genes, a uniform increase in TSS- and gene body-pol II was observed in the OGDH-negative genes, in growth-induced v. normal hearts (Fig. 6c and supplementary Fig. 7S-d). Notably, gene ontology analysis of these genes included the terms sarcomere, Z disc, myofibril,‥etc. that characterize cardiac muscle-specific genes (supplementary Table 7). These have relatively very low or no detectable H2A.Z, as seen in figures 5a (*Actc1 and Actn2*) and 6a (*Tnnt2*), figures 6c and supplementary 7S-d, and as we have previously reported ^17^.

**Table 4.**
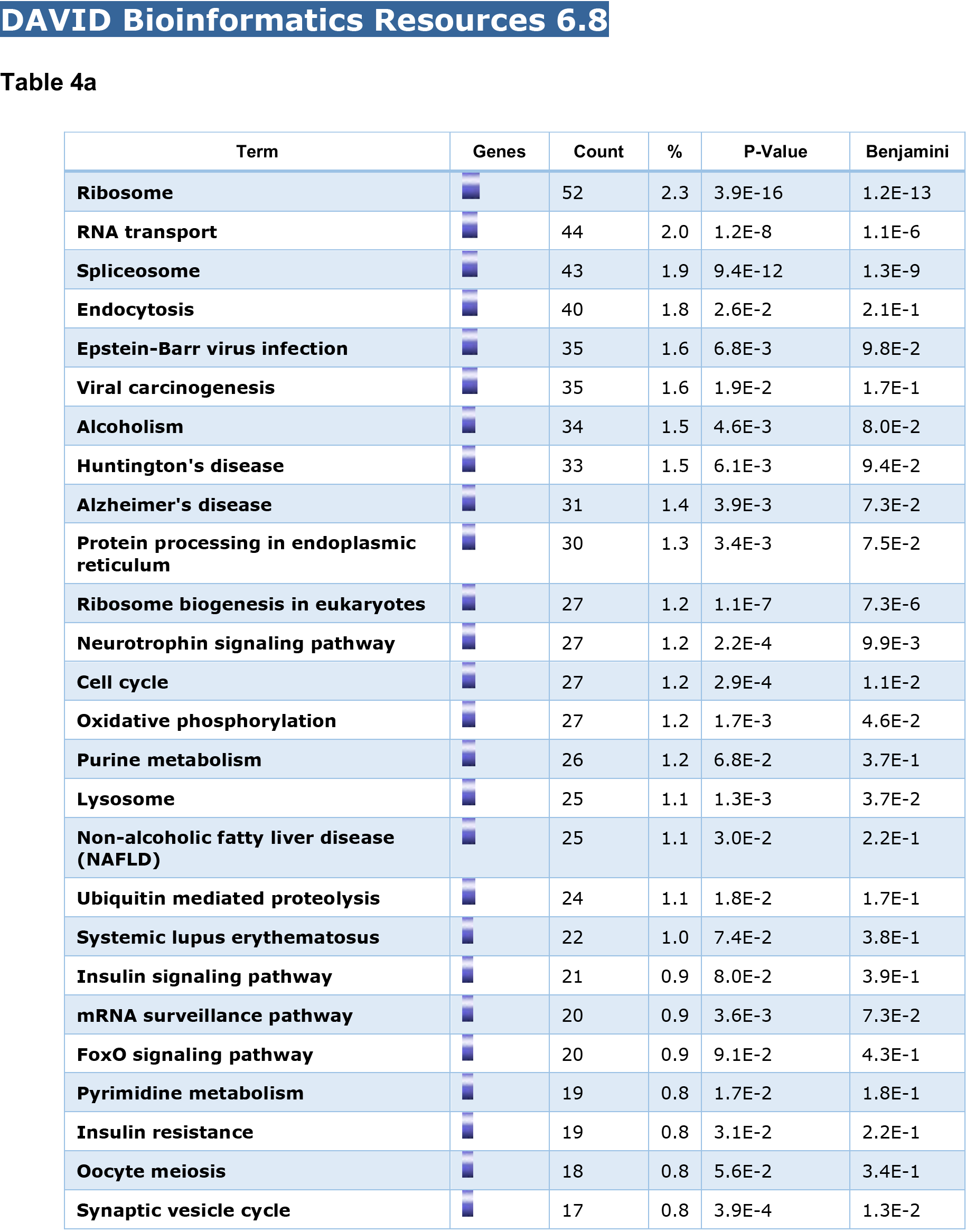
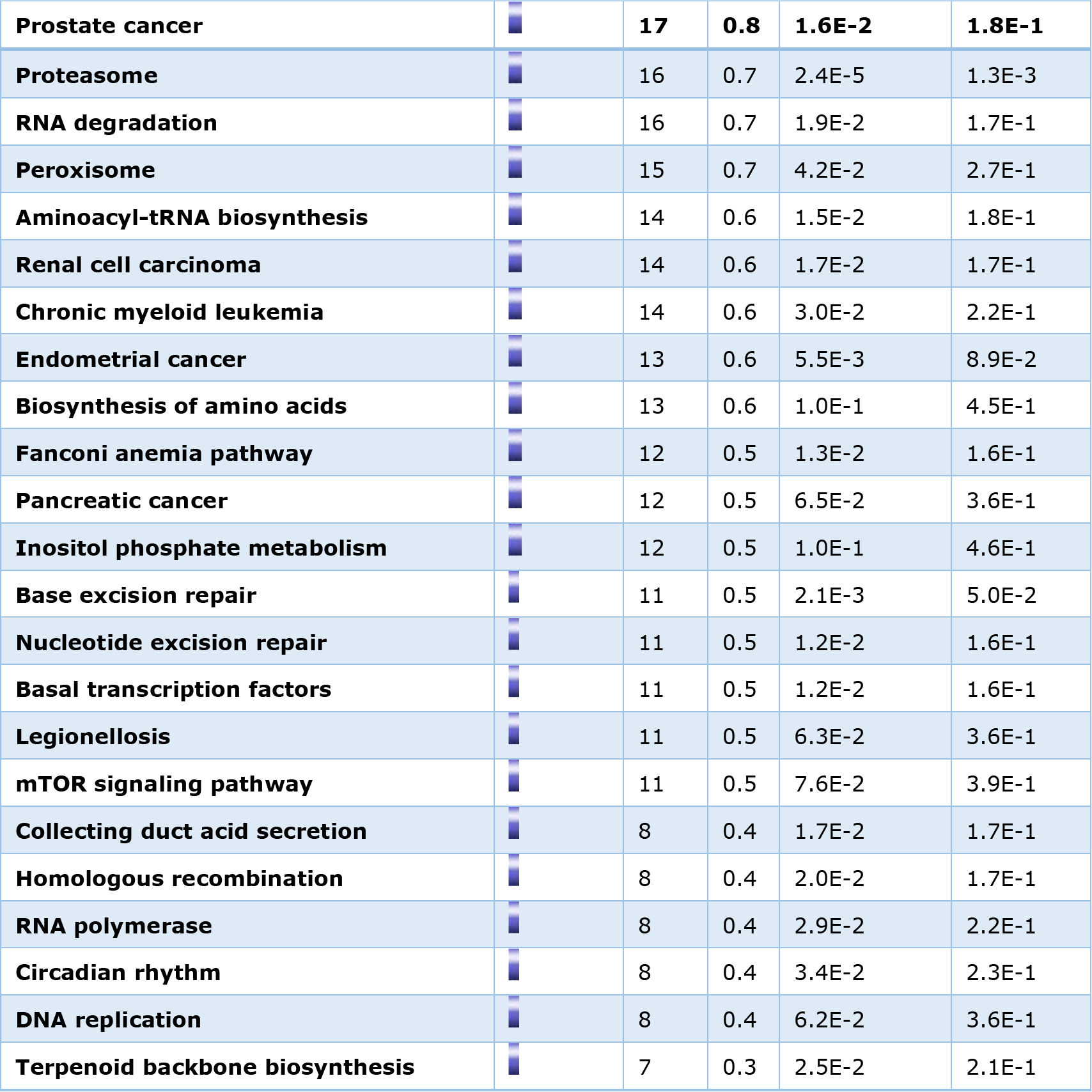
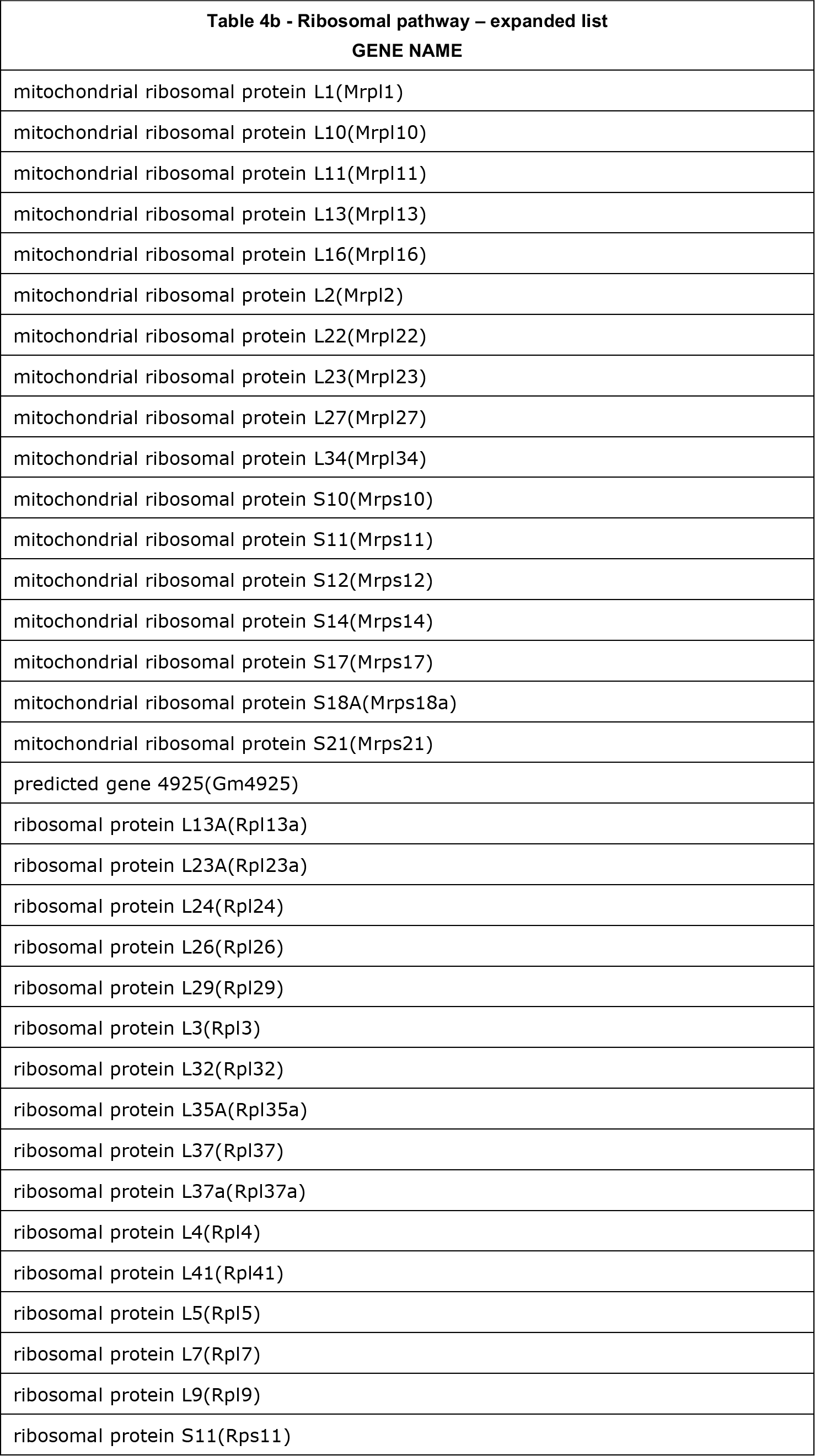
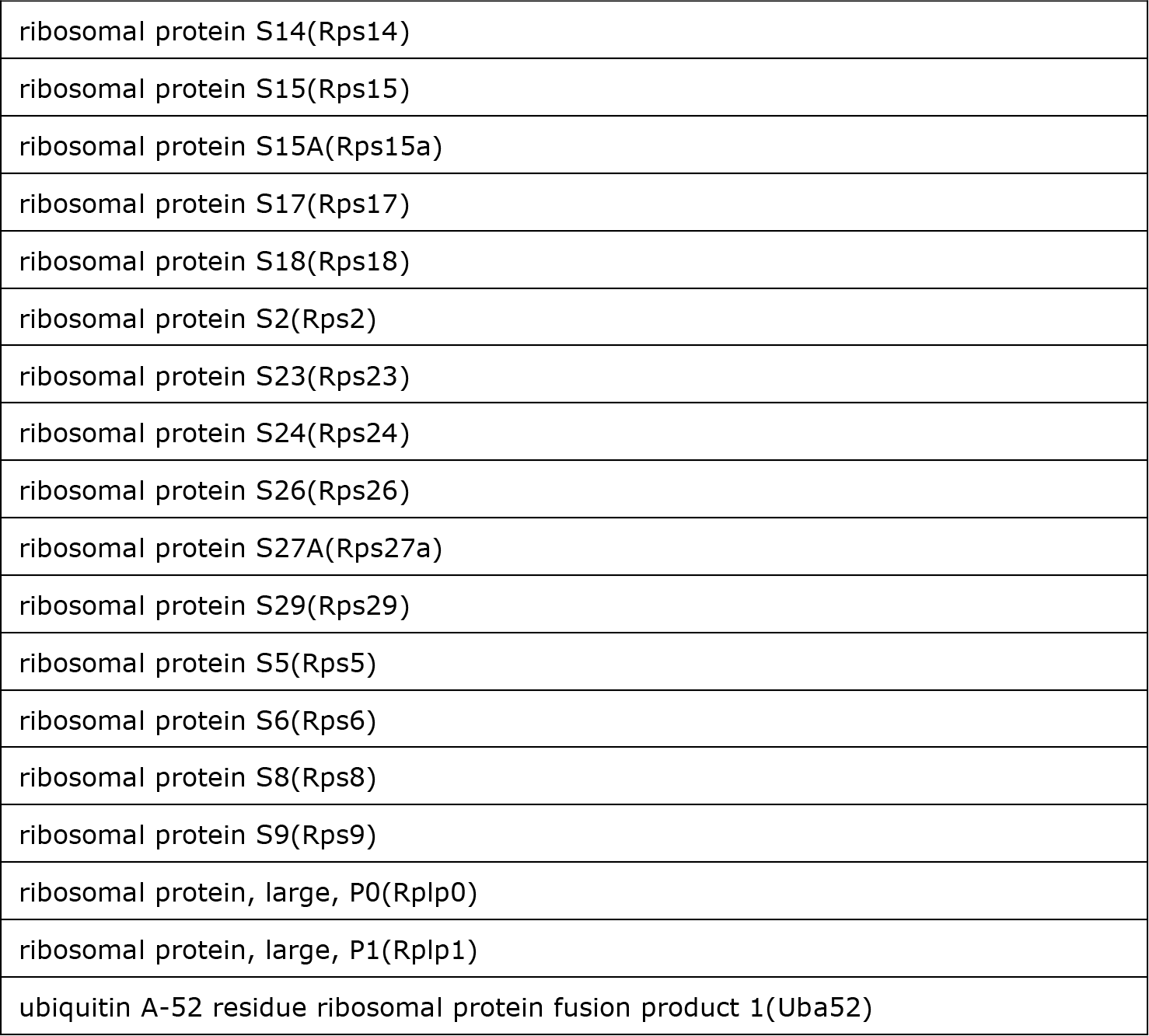
KEGG functional pathway analysis of genes that exhibit no change in ACAA2 abundance at the TSS during TAC (2343 genes)

| Term | Genes | Count | % | P-Value | Benjamini |
| --- | --- | --- | --- | --- | --- |
| **Ribosome** |  | 52 | 2.3 | 3.9E-16 | 1.2E-13 |
| **RNA transport** |  | 44 | 2.0 | 1.2E-8 | 1.1E-6 |
| **Spliceosome** |  | 43 | 1.9 | 9.4E-12 | 1.3E-9 |
| **Endocytosis** |  | 40 | 1.8 | 2.6E-2 | 2.1E-1 |
| **Epstein-Barr virus infection** |  | 35 | 1.6 | 6.8E-3 | 9.8E-2 |
| **Viral carcinogenesis** |  | 35 | 1.6 | 1.9E-2 | 1.7E-1 |
| **Alcoholism** |  | 34 | 1.5 | 4.6E-3 | 8.0E-2 |
| **Huntington's disease** |  | 33 | 1.5 | 6.1E-3 | 9.4E-2 |
| **Alzheimer's disease** |  | 31 | 1.4 | 3.9E-3 | 7.3E-2 |
| **Protein processing in endoplasmic reticulum** |  | 30 | 1.3 | 3.4E-3 | 7.5E-2 |
| **Ribosome biogenesis in eukaryotes** |  | 27 | 1.2 | 1.1E-7 | 7.3E-6 |
| **Neurotrophin signaling pathway** |  | 27 | 1.2 | 2.2E-4 | 9.9E-3 |
| **Cell cycle** |  | 27 | 1.2 | 2.9E-4 | 1.1E-2 |
| **Oxidative phosphorylation** |  | 27 | 1.2 | 1.7E-3 | 4.6E-2 |
| **Purine metabolism** |  | 26 | 1.2 | 6.8E-2 | 3.7E-1 |
| **Lysosome** |  | 25 | 1.1 | 1.3E-3 | 3.7E-2 |
| **Non-alcoholic fatty liver disease (NAFLD)** |  | 25 | 1.1 | 3.0E-2 | 2.2E-1 |
| **Ubiquitin mediated proteolysis** |  | 24 | 1.1 | 1.8E-2 | 1.7E-1 |
| **Systemic lupus erythematosus** |  | 22 | 1.0 | 7.4E-2 | 3.8E-1 |
| **Insulin signaling pathway** |  | 21 | 0.9 | 8.0E-2 | 3.9E-1 |
| **mRNA surveillance pathway** |  | 20 | 0.9 | 3.6E-3 | 7.3E-2 |
| **FoxO signaling pathway** |  | 20 | 0.9 | 9.1E-2 | 4.3E-1 |
| **Pyrimidine metabolism** |  | 19 | 0.8 | 1.7E-2 | 1.8E-1 |
| **Insulin resistance** |  | 19 | 0.8 | 3.1E-2 | 2.2E-1 |
| **Oocyte meiosis** |  | 18 | 0.8 | 5.6E-2 | 3.4E-1 |
| **Synaptic vesicle cycle** |  | 17 | 0.8 | 3.9E-4 | 1.3E-2 |
| **Prostate cancer** |  | **17** | **0.8** | **1.6E-2** | **1.8E-1** |
| **Proteasome** |  | 16 | 0.7 | 2.4E-5 | 1.3E-3 |
| **RNA degradation** |  | 16 | 0.7 | 1.9E-2 | 1.7E-1 |
| **Peroxisome** |  | 15 | 0.7 | 4.2E-2 | 2.7E-1 |
| **Aminoacyl-tRNA biosynthesis** |  | 14 | 0.6 | 1.5E-2 | 1.8E-1 |
| **Renal cell carcinoma** |  | 14 | 0.6 | 1.7E-2 | 1.7E-1 |
| **Chronic myeloid leukemia** |  | 14 | 0.6 | 3.0E-2 | 2.2E-1 |
| **Endometrial cancer** |  | 13 | 0.6 | 5.5E-3 | 8.9E-2 |
| **Biosynthesis of amino acids** |  | 13 | 0.6 | 1.0E-1 | 4.5E-1 |
| **Fanconi anemia pathway** |  | 12 | 0.5 | 1.3E-2 | 1.6E-1 |
| **Pancreatic cancer** |  | 12 | 0.5 | 6.5E-2 | 3.6E-1 |
| **Inositol phosphate metabolism** |  | 12 | 0.5 | 1.0E-1 | 4.6E-1 |
| **Base excision repair** |  | 11 | 0.5 | 2.1E-3 | 5.0E-2 |
| **Nucleotide excision repair** |  | 11 | 0.5 | 1.2E-2 | 1.6E-1 |
| **Basal transcription factors** |  | 11 | 0.5 | 1.2E-2 | 1.6E-1 |
| **Legionellosis** |  | 11 | 0.5 | 6.3E-2 | 3.6E-1 |
| **mTOR signaling pathway** |  | 11 | 0.5 | 7.6E-2 | 3.9E-1 |
| **Collecting duct acid secretion** |  | 8 | 0.4 | 1.7E-2 | 1.7E-1 |
| **Homologous recombination** |  | 8 | 0.4 | 2.0E-2 | 1.7E-1 |
| **RNA polymerase** |  | 8 | 0.4 | 2.9E-2 | 2.2E-1 |
| **Circadian rhythm** |  | 8 | 0.4 | 3.4E-2 | 2.3E-1 |
| **DNA replication** |  | 8 | 0.4 | 6.2E-2 | 3.6E-1 |
| **Terpenoid backbone biosynthesis** |  | 7 | 0.3 | 2.5E-2 | 2.1E-1 |

| **Table 4b - Ribosomal pathway – expanded list** |
| --- |
| **GENE NAME** |
| mitochondrial ribosomal protein L1(Mrpl1) |
| mitochondrial ribosomal protein L10(Mrpl10) |
| mitochondrial ribosomal protein L11(Mrpl11) |
| mitochondrial ribosomal protein L13(Mrpl13) |
| mitochondrial ribosomal protein L16(Mrpl16) |
| mitochondrial ribosomal protein L2(Mrpl2) |
| mitochondrial ribosomal protein L22(Mrpl22) |
| mitochondrial ribosomal protein L23(Mrpl23) |
| mitochondrial ribosomal protein L27(Mrpl27) |
| mitochondrial ribosomal protein L34(Mrpl34) |
| mitochondrial ribosomal protein S10(Mrps10) |
| mitochondrial ribosomal protein S11(Mrps11) |
| mitochondrial ribosomal protein S12(Mrps12) |
| mitochondrial ribosomal protein S14(Mrps14) |
| mitochondrial ribosomal protein S17(Mrps17) |
| mitochondrial ribosomal protein S18A(Mrps18a) |
| mitochondrial ribosomal protein S21(Mrps21) |
| predicted gene 4925(Gm4925) |
| ribosomal protein L13A(Rpl13a) |
| ribosomal protein L23A(Rpl23a) |
| ribosomal protein L24(Rpl24) |
| ribosomal protein L26(Rpl26) |
| ribosomal protein L29(Rpl29) |
| ribosomal protein L3(Rpl3) |
| ribosomal protein L32(Rpl32) |
| ribosomal protein L35A(Rpl35a) |
| ribosomal protein L37(Rpl37) |
| ribosomal protein L37a(Rpl37a) |
| ribosomal protein L4(Rpl4) |
| ribosomal protein L41(Rpl41) |
| ribosomal protein L5(Rpl5) |
| ribosomal protein L7(Rpl7) |
| ribosomal protein L9(Rpl9) |
| ribosomal protein S11(Rps11) |
| ribosomal protein S14(Rps14) |
| ribosomal protein S15(Rps15) |
| ribosomal protein S15A(Rps15a) |
| ribosomal protein S17(Rps17) |
| ribosomal protein S18(Rps18) |
| ribosomal protein S2(Rps2) |
| ribosomal protein S23(Rps23) |
| ribosomal protein S24(Rps24) |
| ribosomal protein S26(Rps26) |
| ribosomal protein S27A(Rps27a) |
| ribosomal protein S29(Rps29) |
| ribosomal protein S5(Rps5) |
| ribosomal protein S6(Rps6) |
| ribosomal protein S8(Rps8) |
| ribosomal protein S9(Rps9) |
| ribosomal protein, large, P0(Rplp0) |
| ribosomal protein, large, P1(Rplp1) |
| ubiquitin A-52 residue ribosomal protein fusion product 1(Uba52) |

**Table 5.**
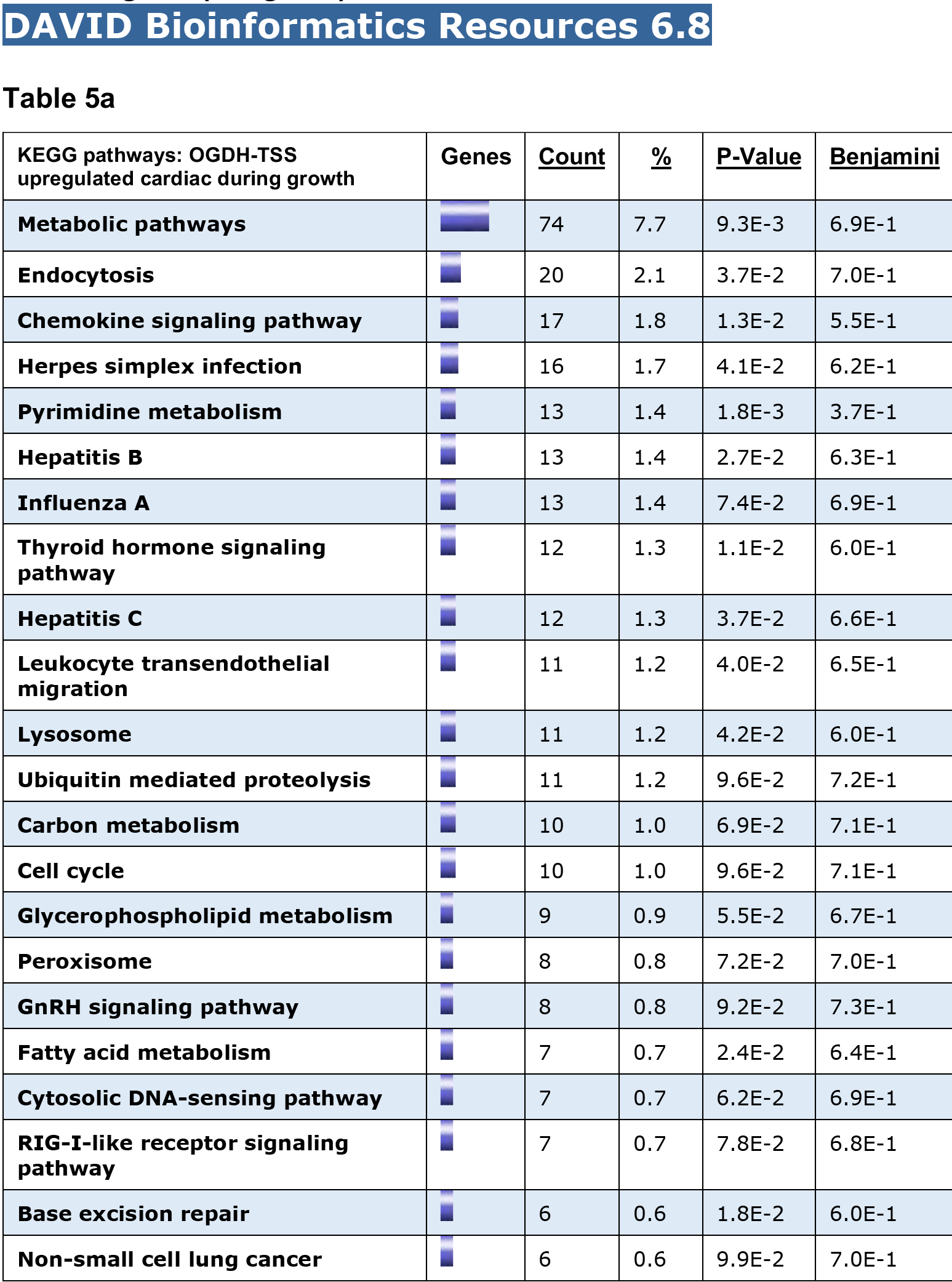
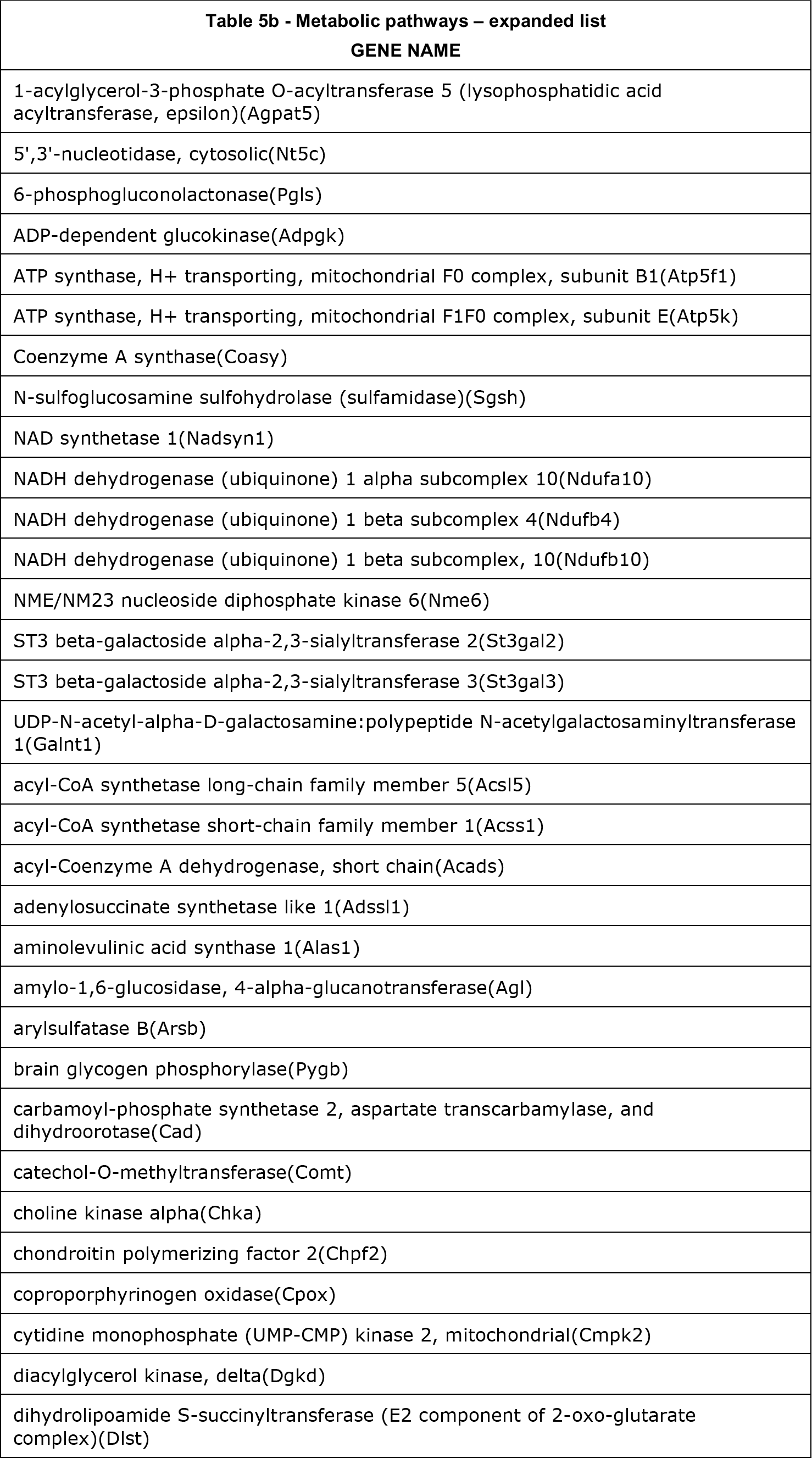
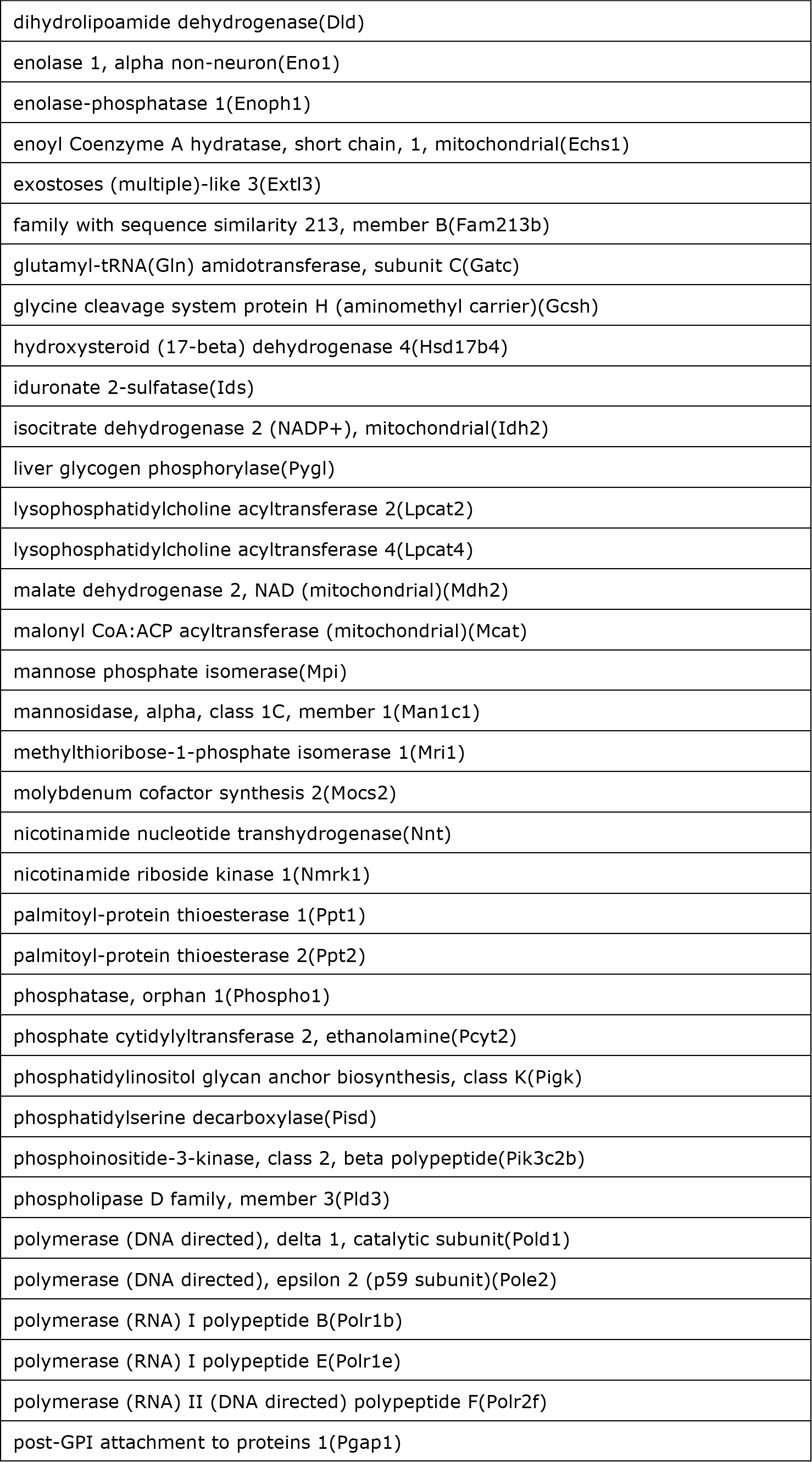
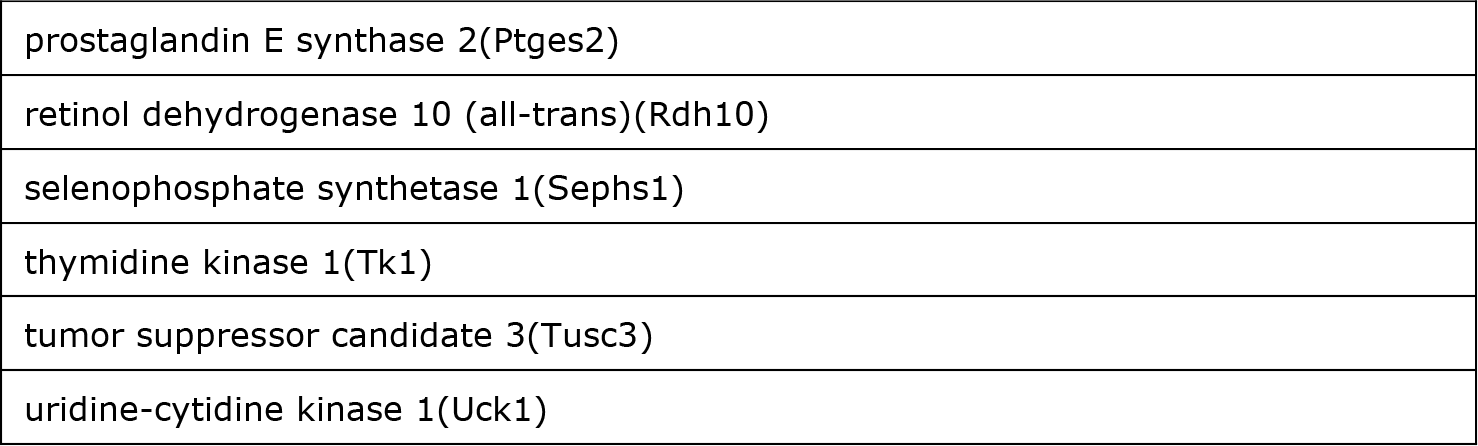
KEGG functional pathway analysis of genes that exhibit upregulation of OGDH at the TSS during TAC (992 genes)

While the vast majority of OGDH peaks overlapped with H2A.Z, 53 peaks appeared to exhibit H2A.Z-independent chromatin binding. Specifically, these peaks were identified in the terminal exon of 53 zinc finger proteins (*Zfp*, Fig. 6a and supplementary Fig. 8S). Ultimately, the binding of OGDH to the TSS of H2A.Z-bound housekeeping and ZFP genes appears to be conserved in humans, as we determined in a human colon cancer cell line (supplementary Fig. 9S-a-c). Thus, the data suggest that OGDH is dependent on H2A.Z for its recruitment to TSSs, however, other factors maybe required for its recruitment to select intragenic sites within *Zfp* genes and that these findings are highly conserved between mouse and human cells.

**Figure 8.**
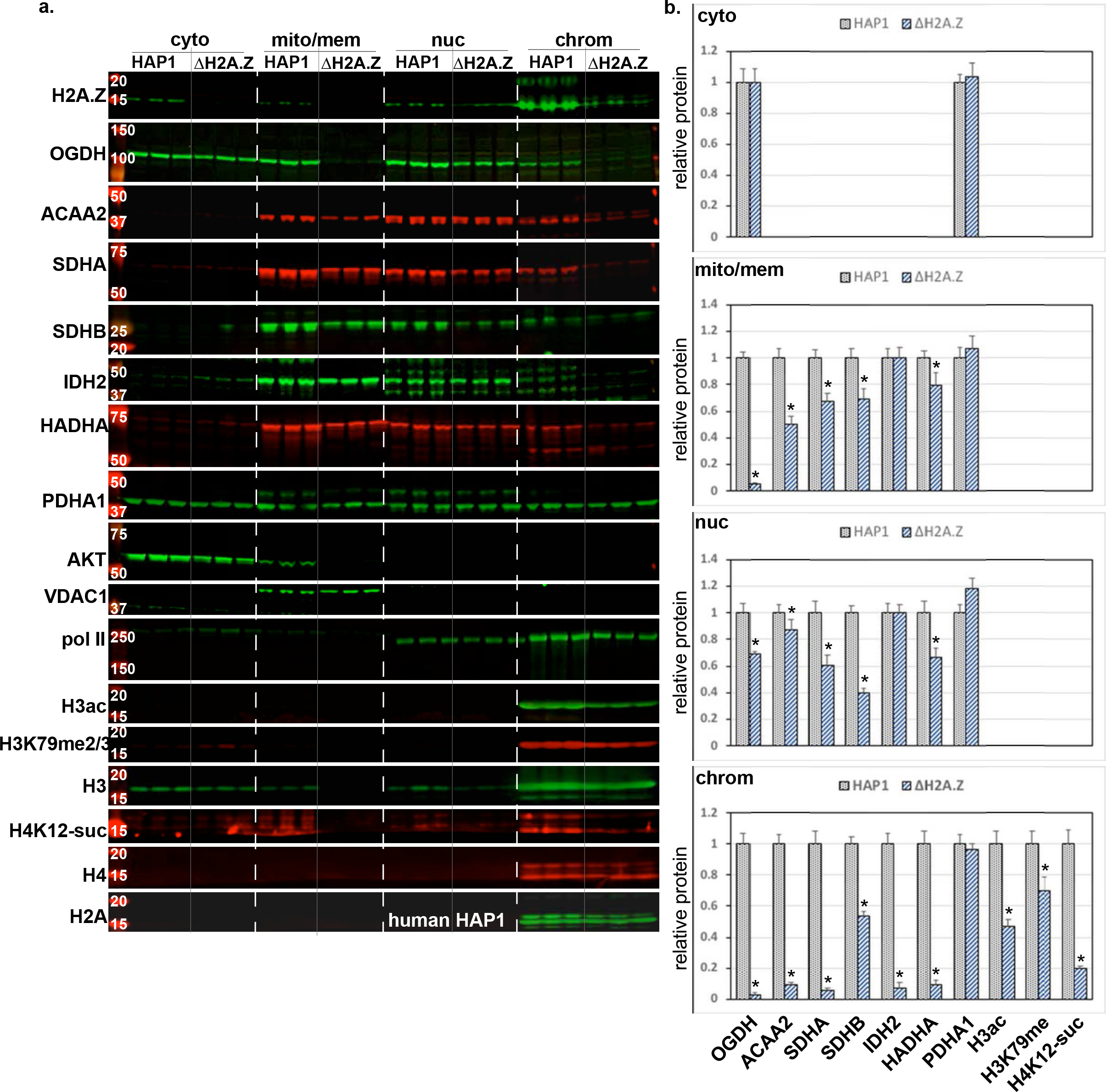
H2A.Z knockout in human HAP1 cells inhibits chromatin binding of metabolic enzymes and reduces histone modifications. **a.** Human HAP1 and H2A.Z-deleted HAP1 (∆H2A.Z) cells were cultured in Iscove’s Modified Dulbecco’s Medium with 10% fetal bovine serum. The cellular protein/organelles were fractionated into cytosol (cyto), mitochondrial and membrane (mito/mem), nuclear (nuc), and chromatin-bound (chrom) protein fractions that were then analyzed by Western blotting for the proteins listed on the left of each panel. **b.** The signals for the enzymes in each of the fractions were quantified by imageJ, normalized to the internal control of the corresponding fraction (AKT1, VDAC1, or Pol II, for the cyto, mito, and nuc and chrom, respectively) and plotted as the mean ± SEM of relative quant of the protein detected in the ∆H2A.Z vs. the parent cells (HAP1) adjusted to 1. H3-ac and K79me2/3 were normalized to H3, and H4K12-suc, was normalized to H4. N=3, each, from 3 repeats, \**p* = 0.001 vs. HAP1 in the corresponding fraction.

### Knockdown of H2A.Z in mouse myocytes reduces chromatin association of metabolic enzymes

To determine the role of H2A.Z in the recruitment of metabolic enzymes to chromatin, we knocked down H2A.Z using short hairpin RNA (sh-H2A.Z) in mACM. This approach induced a significant reduction of chromatin-bound H2A.Z (63 ± 3 %), PDHA1 (62 ± 14 %), ACAA2 (65 ± 4.5), H3ac (70 ± 3%), and H3K27me2/3, (56 ± 8%), vs. control levels, whereas H3 remained unchanged (Fig. 7a-b). Note that the cells morphology/viability remained intact during the 24 h period of this experiment (supplementary Fig. 10S), and that similar results were observed in rNCM (supplementary Fig. 11S). PDK1 and TFIIB were used as mitochondrial and nucleoplasm markers. Although TFIIB directly binds to DNA elements near the TATA-box, this interaction is not preserved in our protein fractionation method, which does not involve protein crosslinking, thus, resulting in its localization to the nucleoplasm. This contrasts with the histones, which are wrapped with chromatin, and, accordingly, strictly localize to the chromatin-bound fraction of proteins. Thus, detection of chromatin-bound PDHA1 and ACAA2, and its disruption by knockdown of H2A.Z, confirms their relatively tight association with chromatin in an H2A.Z-dependent fashion. Moreover, the data show that H2A.Z is required for H3 acetylation and methylation, plausibly as a result of recruitment of metabolic enzymes. This is further supported by a reduction in H3ac after knockdown of ACAA2 (supplementary Fig. 12S).

### Knockdown of OGDH inhibits H4 succinylation

OGDH has been reported to bind to chromatin in U251 glioblastoma cells where it mediates succinylation of H3K79 ^14^. To test the impact of OGDH on histone succinylation in normal cardiac myocytes, we knocked it down using shRNA (sh-OGDH). This treatment induced a significant reduction of chromatin-bound OGDH (79 ± 7 %), and H4K12suc (62 ± 7 %) v. control levels, but not of H3K27me2/3, H3K9me1/2/3, H3, or H4. TFIIB was used as a nuclear marker. In addition, knockdown of OGDH was associated with a reduction in chromatin-bound ACAA2, indicative of the codependence of ACCA2 on OGDH for its recruitment to chromatin. We conclude that OGDH is required for histone succinylation, plausibly through conversion of alpha-ketoglutarate into succinyl-CoA at TSSs. On the other hand, we predict the reduction in H3 acetylation maybe secondary to the reduction in chromatin-bound ACAA2, since knockdown of ACAA2, resulted in 88% reduction in H3 acetylation (supplementary Fig. 11S).

### H2A.Z knockout in human HAP1 cells inhibits chromatin association of metabolic enzymes and posttranslational histone modifications

To validate the above data and investigate its relevance in human cells, we analyzed human near-haploid HAP1 cells with a 2 bp deletion in exon 3 of H2A.Z (∆H2A.Z). These cells are viable, however, they proliferate at ~1/4 of the rate of the parent cells (supplementary Fig. 13S). After fractionating the cellular protein/organelles and analyzing it with Western blots, we confirmed that H2A.Z is deficient in the ∆H2A.Z cells (Fig. 8c). This loss is associated with more than 90% reduction in chromatin bound mitochondrial enzymes, including OGDH, ACAA2, HADHA, IDH2, SDHA and SDHB, and to a lesser extent their nucleoplasmic levels (Fig. 8a-b). Except for IDH2, the mitochondrial content of these enzymes was also reduced, whereas OGDH was undetectable. We predict that this reduction in total enzyme content is a result of direct, or indirect, H2A.Z-dependent transcription of their genes. As for OGDH, it is unclear why it is completely lost from the mitochondria, in particular, in the absence of H2A.Z.

In contrast to the above tested enzymes, while the results show that PDHA1 exhibited strong localization to the nuclear and chromatin fractions, it was the only enzyme for which neither its expression nor chromatin binding were impacted by the knockout of H2A.Z. This proved its H2A.Z-independent chromatin association and, thereby, the selectivity of H2A.Z-dependent recruitment of metabolic enzymes. Other noted differences between the enzymes, include the finding that only OGDH and PDHA1 were detected in the cytosol, the unexpected complete loss of OGDH in the mitochondrial fraction in the ∆H2A.Z cells, and the equivalent reductions of SDHB in all fractions in the ∆H2A.Z cells that suggests its independence of H2A.Z for chromatin binding. Also, notable, is the fact that the nuclear localization of mitochondrial enzymes was selective, since the mitochondrial proteins VDAC1 and PDK1 were not detected in the nucleus, and were, thus, used as mitochondrial markers and internal controls in our blots. Thus,

## DISCUSSION

In this study, we have identified a plethora of metabolic enzymes that bind to the TSSs of transcriptionally active genes in a H2A.Z-dependent and, less frequently, -independent fashion. These were discovered by an unbiased screen using anti-H2A.Z chromatin immunoprecipitation-mass spectrometry. This approach provides the unique advantage of identifying proteins that associate with H2A.Z in its native conformation within the nucleosome. One of the disadvantages, though, as with other immunoprecipitation approaches, is the likelihood of non-specific bindings. To eliminate those from our analysis, we applied the following measures; each sample analyzed consisted of a pool of 20 independent heart, each sample was analyzed twice by mass spec, the H2A.Z pulldown was analyzed in 2 independent samples (the normal heart and the growth-induced), we only considered the co-immunoprecipitated proteins that exhibited ≥ 2-fold enrichment with H2A.Z vs. IgG control, and finally, we validated this finding for 7 of 29 enzymes identified using a combination of various methods that included immunocytochemistry, tGFP fusion proteins, NLS mutagenesis, ChIP-Seq, and H2A.Z knockout. So far, all the metabolic enzymes that we have tested, including ACAA2, OGDH, IDH2, PDHA1, HADHA, SDHA, and SDHB, were confirmed for their nuclear localization and chromatin binding by two or more of the methods listed above, providing confidence in our RIME findings.

We preformed the RIME assay in total heart tissue for the purpose of preserving the 3D milieu of the cells, which is critical for their transcriptional integrity, as we ascertained that the signals obtained by this approach are predominantly derived from cardiac myocytes. This is supported by the fact that smooth muscle actin (*Acta2*), which is expressed in smooth muscle cells and myofibroblasts in the heart, and ATPase plasma membrane Ca^2+^ transporting 4 (*Atp2b4*), which is ubiquitously expressed, including in epithelial cells, have no detectable RNA pol II binding compared to its high abundance in the corresponding cardiac genes, cardiac actin (*Actc1*) and ATPase sarcoplasmic/endoplasmic reticulum Ca^2+^ transporting 2 (*Atp2a2*, supplementary Fig. 14S). We then confirmed the findings by the various methods listed above, in both rodent and human cell lines, including isolated mouse and rat myocytes, human iPSC-derived cardiac myocytes, human colon cancer cells, and human near-haploid cells with or without a H2A.Z deletion. We find that the nuclear localization of the metabolic enzymes is conserved between species and largely H2A.Z-dependent.

H2A.Z is a highly conserved histone variant that plays an essential role in sensing and responding to metabolic and environmental cues, however, the underlying mechanisms remain elusive. It has, though, been shown to interact with 93 proteins, mostly identified by unbiased screens using co-fractionation, affinity capture-MS, and affinity capture-Western [compiled in BioGRID ^20^], some of which may mediate its functions. These proteins include core histones ^21, 22^, proteins that regulate H2A.Z’s chromatin deposition [e.g. INO80 complex subunit C ^23^, vacuolar protein sorting 72 ^24^, ANP32E ^17, 25^], epigenetic regulators [e.g. E1A binding protein p400, lysine acetyltransferase 5, and histone deacetylase 1 and 2 ^22^], in addition to its interaction with the metabolic enzymes ACAA2 and fumarase, observed in an interactome identified by co-fractionation ^26^. However, most of these interactions were not identified or validated in the context of nucleosomal-bound H2A.Z. Using the rapid immunoprecipitation-mass spectrometry of endogenous proteins assay approach ^18^ with anti-H2A.Z we identified its interaction with core histones, however, most of the other associated proteins were those related to metabolism, including all the enzymes of the TCA cycle and β-oxidation spiral, and key enzymes in the branched-chain amino acid metabolism. The fact that these enzymes are associated with chromatin at the TSS explains how metabolites could be directly delivered to target genes where they are used as substrates for histone modifications (e.g. acetyl-CoA, succinyl-CoA, among other short acyl-CoA metabolites) or as co-factors (αKG) for histone modifying enzymes, thereby, allowing promoters to immediately sense and respond to metabolic cues. Consistent with an H2A.Z-dependent recruitment, genes that are devoid of H2A.Z, also lack metabolic enzymes (ACAA2 and OGDH) at their TSS and, conclusively, knockdown or knockout of H2A.Z abrogates chromatin association of multiple enzymes including ACAA2, OGDH, IDH2, HADHA, SDHA, and SDHB. Interestingly, however, PDHA1 was an exception, as it retained its chromatin association in the absence of H2A.Z in HAP1 cells. Therefore, we conclude that while H2A.Z may be required for the recruitment of multiple metabolic enzymes to chromatin, there are some that are H2A.Z-independent, for which the recruiting protein remains to be determined. On the other hand, we speculate that the absence of H2A.Z and metabolic enzymes at the promoters of constitutively expressed, tissue-restricted genes (e.g. sarcomeric proteins), which distinguish an organ’s unique functionality, ensures these are not impacted by metabolic fluctuations, as one would expect. This contrasts with housekeeping and inducible genes that would be immediately modulated in response to changes in oxygen and/or metabolic substrate availabilities, as a mechanism of cellular adaptation. Our results are also consistent with a proteomics study that identified all TCA cycle enzymes in the nucleus of normal and cancer cells ^16^.

H2A.Z is required for transcriptional memory in yeast where it is incorporated in newly deactivated *INO1* at the nuclear periphery ^27, 28^, and in hippocampal memory in mice, where it negatively controls fear memory through suppressing the expression of specific memory-activating genes ^29^, and is necessary for neurogenesis and normal behavioral traits in mice ^30^. As well established, memory is a function of epigenetics, wherein histone acetylation is an essential regulator, demonstrated by Mews et al ^31^. In that study, the authors reported that ACSS2 (a cytosolic and nuclear enzyme that converts acetate into acetyl-CoA) binds near the TSS of hippocampal neuronal genes, where its knockdown diminishes long-term spatial memory and an acetylation-dependent cognitive process. This suggests a potential link between H2A.Z and the intricate and precise regulation of core histone modifications. Concordantly, we show that knockdown or knockout of H2A.Z reduces the association of metabolic enzymes with chromatin, which is paralleled with a significant reduction in histone H3 acetylation and methylation, whereas OGDH knockdown or H2A.Z knockout reduce H4K12 succinylation.

The association of metabolic enzymes with the TSS of genes could potentially explain the mechanism of targeted histone modifications, and, therefore, the direct regulation of transcription via glucose and fatty acid metabolism. Furthermore, the data expands the range of locally-delivered modifying substrates and regulatory co-factor to include not only acetyl-CoA or succinyl-CoA, but also citrate, αKG, succinate, fumarate, and other short chain acyl-CoAs, which considering the complexity of gene regulation and memory, is not surprising.

While acetylation and methylation of histones is a key player in transcriptional regulation and memory, succinylation is another modification that has received less attention. The alpha-ketoglutarate dehydrogenase complex catalyzes the conversion of αKG into succinyl-CoA as a source of cellular succinyl-CoA. However, there is no known route via which this metabolic intermediate is delivered to the nucleus. Interestingly, Wang et al, identified an interaction between OGDH and lysine acetyltransferase 2A in the nucleus, which mediates the succinylation of H3K79 ^14^. Their report shows that ChIP-Seq of OGDH, in U251 glioblastoma cells, identifies 249 peaks, mainly enriched within 2 Kb of the TSS. This agrees with what we observed in our study, where OGDH is enriched at −2000 to +2000 surrounding the TSSs. The differences, though, are that we identified 16,790 peaks, covering 89.9% of TSSs that co-localized and co-immunoprecipitated with H2A.Z, with the exemption of tissue-restricted genes, which lack any substantial amount of H2A.Z. We predict that the difference in the number of peaks is likely due to differences in the cell types or the antibodies used for ChIP-Seq. Other support of nuclear localization of mitochondrial enzymes was reported by Jiang et al, showing that phosphorylated fumarase interacts with H2A.Z in response to ionizing radiation-induced activation of DNA-dependent protein kinase in U2OS cells, a function that regulates DNA repair ^32^. Our study extends these findings to include nuclear occupancy of all the enzymes in the TCA cycle and beta-oxidation spiral, where they are recruited to chromatin via H2A.Z. Thus, our data support the concept that local production of the TCA cycle intermediates is necessary for transcriptional regulation. Other than fumarate and succinyl-CoA, production of citrate, αKG, succinate, and acetyl-CoA, among others, have the capacity of either generating the substrates that are directly required for histone modification, or alternatively, generating metabolites that regulate histone modifying enzymes.

H2A.Z’s function does not always correlate with transcriptional activity, as noted by others and us. Not only do we show that constitutively-expressed, tissue-restricted genes, which have the highest level of cellular expression, have little to no H2A.Z at the TSS or in the gene body, but conversely, we found that developmental suppressed/unexpressed genes (e.g. *Wnt1, Noggin, Tbx1*) have substantial amounts of H2A.Z at the TSS and in gene body (supplementary data, Fig. 15S and Table 8). On the other hand, moderately-expressed housekeeping genes that are amenable to incremental modulation by external stimuli have the highest levels of H2A.Z at their TSS, whereas, inducible genes have high levels of H2A.Z at the TSS that extends into the gene body. The latter pattern enhances the responsiveness of inducible genes to stimuli, as previously reported in *Arabidopsis thaliana* ^33^. While these results support the role for H2A.Z in transcriptional regulation, they suggest that it is not required for constitutive transcription but rather for strictly regulated transcription. Our findings show that ACAA2 and OGDH exist only at H2A.Z-occupied TSSs of expressed and minimally of a few unexpressed genes (supplementary Fig. 15S), where H2A.Z and OGDH fully overlap at 89.9% of expressed TSSs. In comparison, chromatin-bound ACAA2 overlaps with H2A.Z and OGDH at 36.5 % of TSSs. There are, however, a handful of genes that are an exception to this rule. These included 53 *Zfp* genes, which have the highest peaks of OGDH within their terminal exon, where there is no detectable H2A.Z, which are conserved in humans (supplementary Fig. 8S and 9S).

**Table 6.**
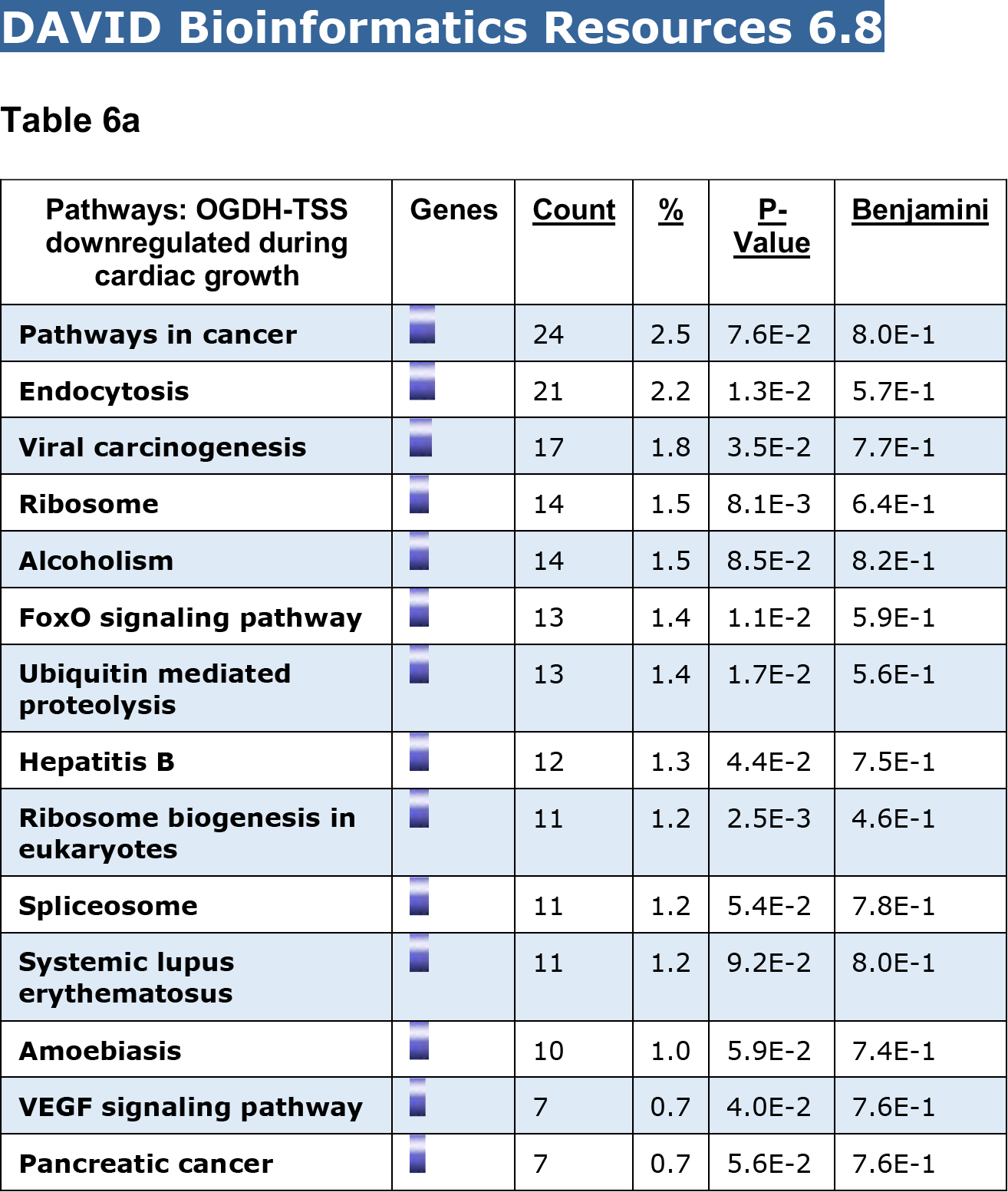
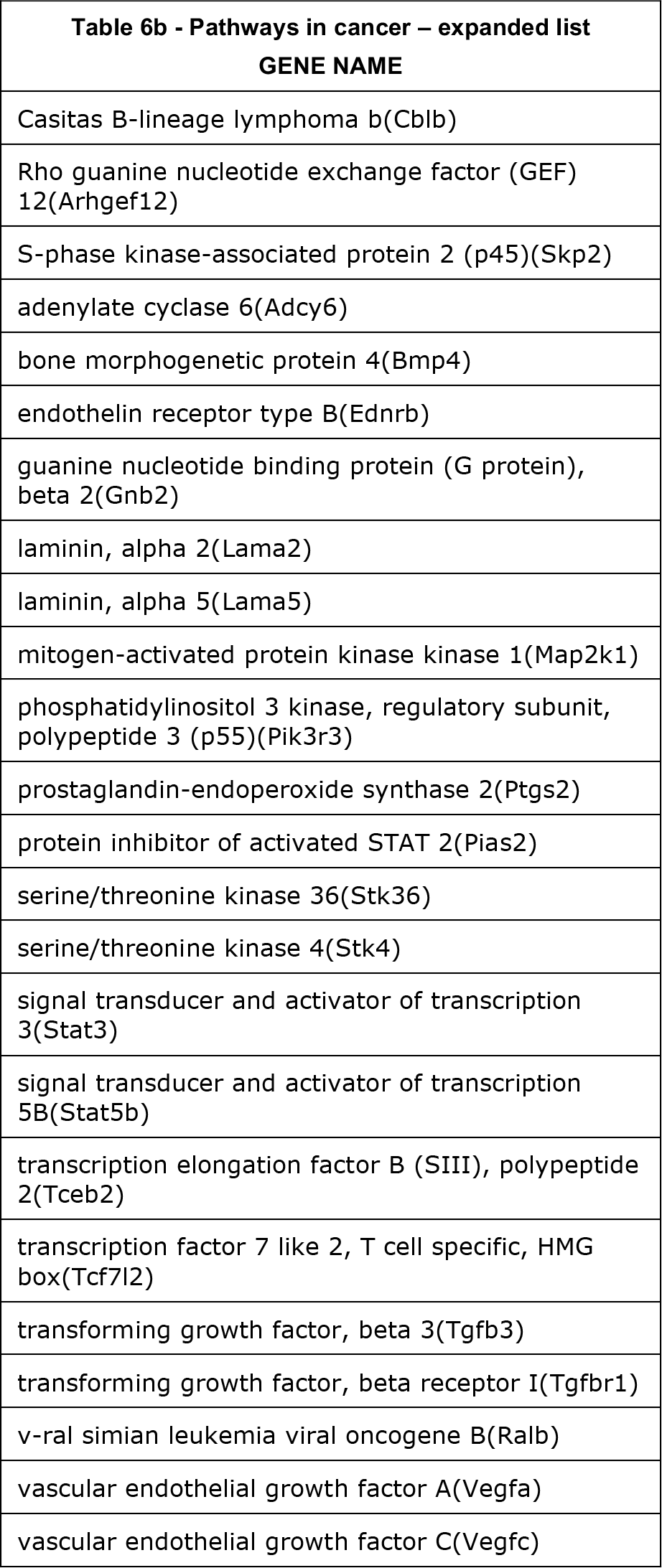
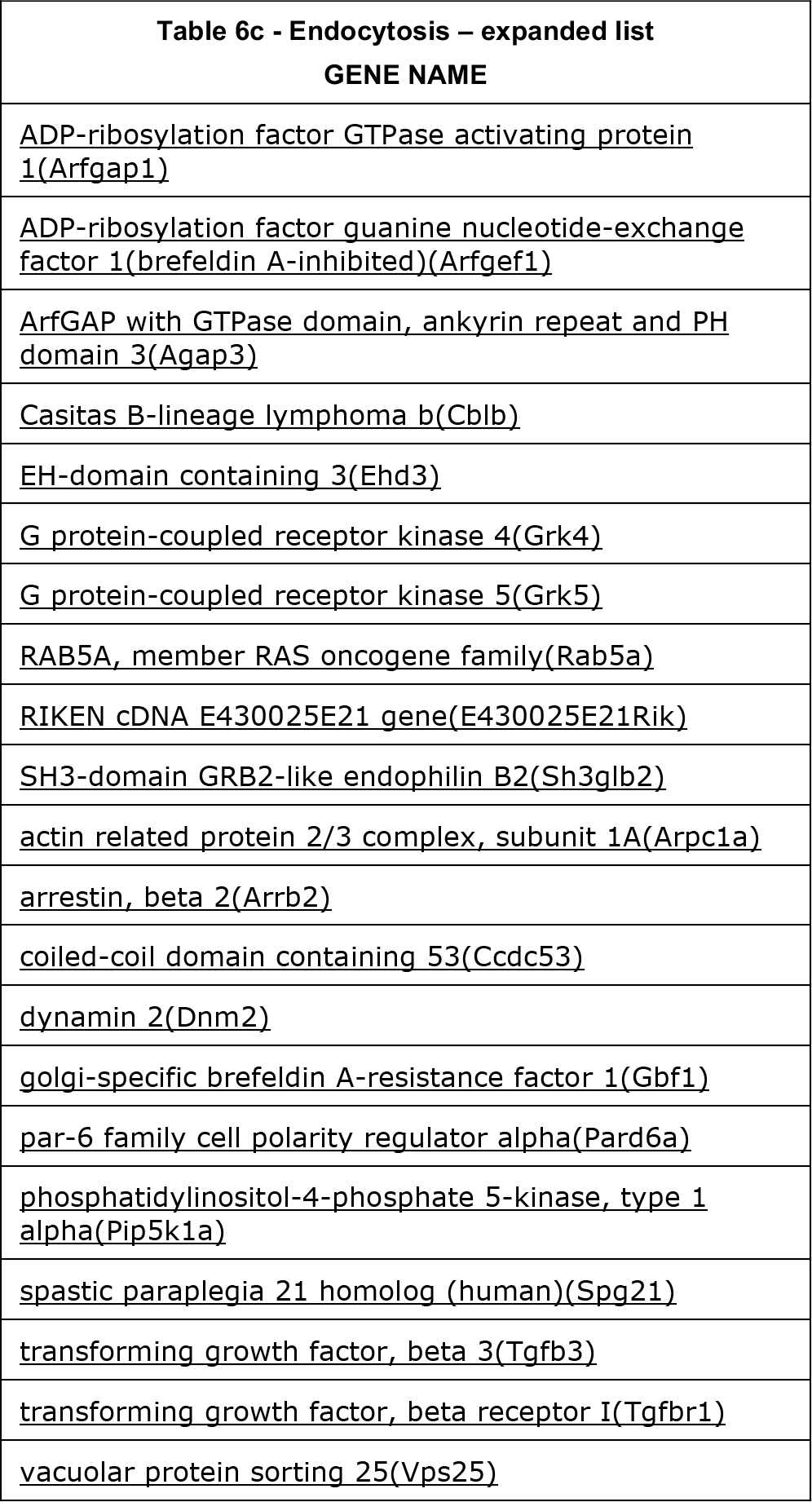
KEGG functional pathway analysis of genes that exhibit downregulation of OGDH at the TSS during TAC (993 genes)

**Table 7a.**
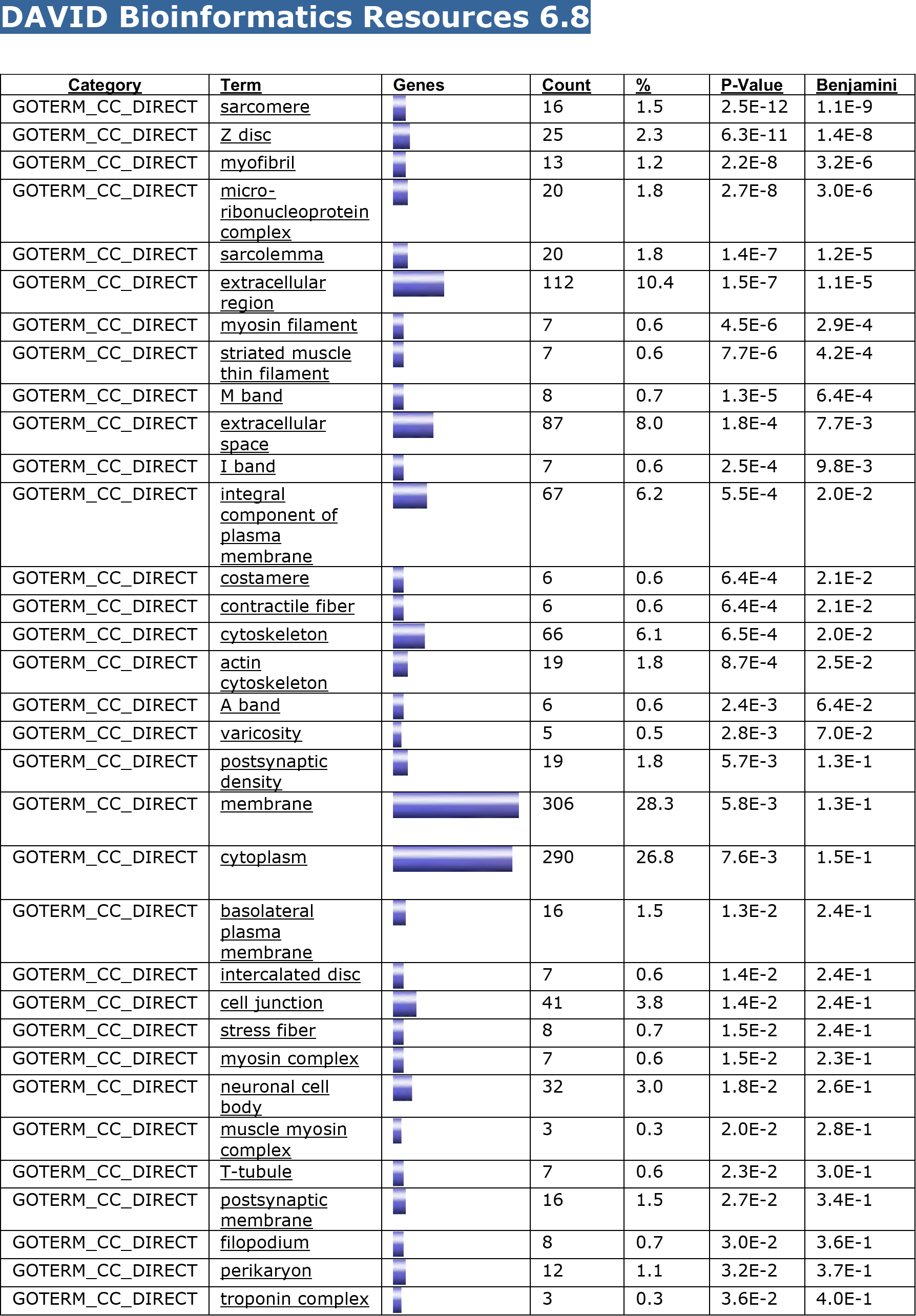
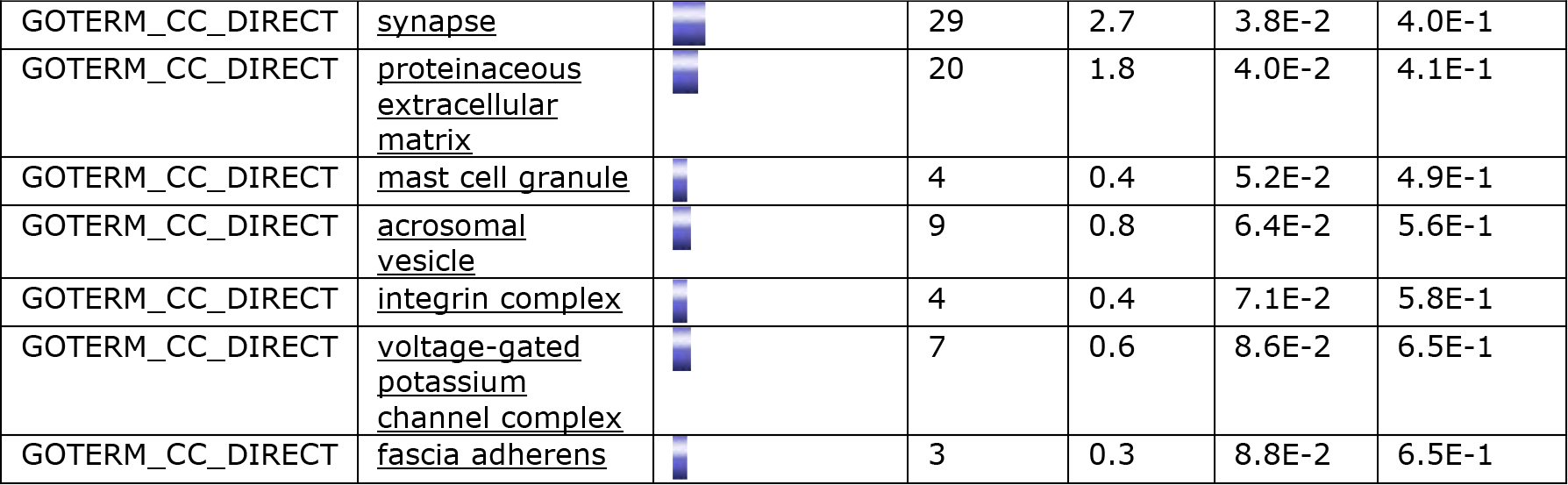
GO Term analysis of genes negative for OGDH at TSS (Total OGDH negative genes = 1165 of 11527 expressed genes in the heart)

| Category | Term | Genes | Count | % | P-Value | Benjamini |
| --- | --- | --- | --- | --- | --- | --- |
| GOTERM_CC_DIRECT | <u>sarcomere</u> |  | 16 | 1.5 | 2.5E-12 | 1.1E-9 |
| GOTERM_CC_DIRECT | <u>Z disc</u> |  | 25 | 2.3 | 6.3E-11 | 1.4E-8 |
| GOTERM_CC_DIRECT | <u>myofibril</u> |  | 13 | 1.2 | 2.2E-8 | 3.2E-6 |
| GOTERM_CC_DIRECT | <u>micro-</u><br><u>ribonucleoprotein</u><br><u>complex</u> |  | 20 | 1.8 | 2.7E-8 | 3.0E-6 |
| GOTERM_CC_DIRECT | <u>sarcolemma</u> |  | 20 | 1.8 | 1.4E-7 | 1.2E-5 |
| GOTERM_CC_DIRECT | <u>extracellular</u><br><u>region</u> |  | 112 | 10.4 | 1.5E-7 | 1.1E-5 |
| GOTERM_CC_DIRECT | <u>myosin filament</u> |  | 7 | 0.6 | 4.5E-6 | 2.9E-4 |
| GOTERM_CC_DIRECT | <u>striated muscle</u><br><u>thin filament</u> |  | 7 | 0.6 | 7.7E-6 | 4.2E-4 |
| GOTERM_CC_DIRECT | <u>M band</u> |  | 8 | 0.7 | 1.3E-5 | 6.4E-4 |
| GOTERM_CC_DIRECT | <u>extracellular</u><br><u>space</u> |  | 87 | 8.0 | 1.8E-4 | 7.7E-3 |
| GOTERM_CC_DIRECT | <u>I band</u> |  | 7 | 0.6 | 2.5E-4 | 9.8E-3 |
| GOTERM_CC_DIRECT | <u>integral</u><br><u>component of</u><br><u>plasma</u><br><u>membrane</u> |  | 67 | 6.2 | 5.5E-4 | 2.0E-2 |
| GOTERM_CC_DIRECT | <u>costamere</u> |  | 6 | 0.6 | 6.4E-4 | 2.1E-2 |
| GOTERM_CC_DIRECT | <u>contractile fiber</u> |  | 6 | 0.6 | 6.4E-4 | 2.1E-2 |
| GOTERM_CC_DIRECT | <u>cytoskeleton</u> |  | 66 | 6.1 | 6.5E-4 | 2.0E-2 |
| GOTERM_CC_DIRECT | <u>actin</u><br><u>cytoskeleton</u> |  | 19 | 1.8 | 8.7E-4 | 2.5E-2 |
| GOTERM_CC_DIRECT | <u>A band</u> |  | 6 | 0.6 | 2.4E-3 | 6.4E-2 |
| GOTERM_CC_DIRECT | <u>varicosity</u> |  | 5 | 0.5 | 2.8E-3 | 7.0E-2 |
| GOTERM_CC_DIRECT | <u>postsynaptic</u><br><u>density</u> |  | 19 | 1.8 | 5.7E-3 | 1.3E-1 |
| GOTERM_CC_DIRECT | <u>membrane</u> |  | 306 | 28.3 | 5.8E-3 | 1.3E-1 |
| GOTERM_CC_DIRECT | <u>cytoplasm</u> |  | 290 | 26.8 | 7.6E-3 | 1.5E-1 |
| GOTERM_CC_DIRECT | <u>basolateral</u><br><u>plasma</u><br><u>membrane</u> |  | 16 | 1.5 | 1.3E-2 | 2.4E-1 |
| GOTERM_CC_DIRECT | <u>intercalated disc</u> |  | 7 | 0.6 | 1.4E-2 | 2.4E-1 |
| GOTERM_CC_DIRECT | <u>cell junction</u> |  | 41 | 3.8 | 1.4E-2 | 2.4E-1 |
| GOTERM_CC_DIRECT | <u>stress fiber</u> |  | 8 | 0.7 | 1.5E-2 | 2.4E-1 |
| GOTERM_CC_DIRECT | <u>myosin complex</u> |  | 7 | 0.6 | 1.5E-2 | 2.3E-1 |
| GOTERM_CC_DIRECT | <u>neuronal cell</u><br><u>body</u> |  | 32 | 3.0 | 1.8E-2 | 2.6E-1 |
| GOTERM_CC_DIRECT | <u>muscle myosin</u><br><u>complex</u> |  | 3 | 0.3 | 2.0E-2 | 2.8E-1 |
| GOTERM_CC_DIRECT | <u>T-tubule</u> |  | 7 | 0.6 | 2.3E-2 | 3.0E-1 |
| GOTERM_CC_DIRECT | <u>postsynaptic</u><br><u>membrane</u> |  | 16 | 1.5 | 2.7E-2 | 3.4E-1 |
| GOTERM_CC_DIRECT | <u>filopodium</u> |  | 8 | 0.7 | 3.0E-2 | 3.6E-1 |
| GOTERM_CC_DIRECT | <u>perikaryon</u> |  | 12 | 1.1 | 3.2E-2 | 3.7E-1 |
| GOTERM_CC_DIRECT | <u>troponin complex</u> |  | 3 | 0.3 | 3.6E-2 | 4.0E-1 |
| GOTERM_CC_DIRECT | <u>synapse</u> |  | 29 | 2.7 | 3.8E-2 | 4.0E-1 |
| GOTERM_CC_DIRECT | <u>proteinaceous</u><br><u>extracellular</u><br><u>matrix</u> |  | 20 | 1.8 | 4.0E-2 | 4.1E-1 |
| GOTERM_CC_DIRECT | <u>mast cell granule</u> |  | 4 | 0.4 | 5.2E-2 | 4.9E-1 |
| GOTERM_CC_DIRECT | <u>acrosomal</u><br><u>vesicle</u> |  | 9 | 0.8 | 6.4E-2 | 5.6E-1 |
| GOTERM_CC_DIRECT | <u>integrin complex</u> |  | 4 | 0.4 | 7.1E-2 | 5.8E-1 |
| GOTERM_CC_DIRECT | <u>voltage-gated</u><br><u>potassium</u><br><u>channel complex</u> |  | 7 | 0.6 | 8.6E-2 | 6.5E-1 |
| GOTERM_CC_DIRECT | <u>fascia adherens</u> |  | 3 | 0.3 | 8.8E-2 | 6.5E-1 |

**Table 7b.**
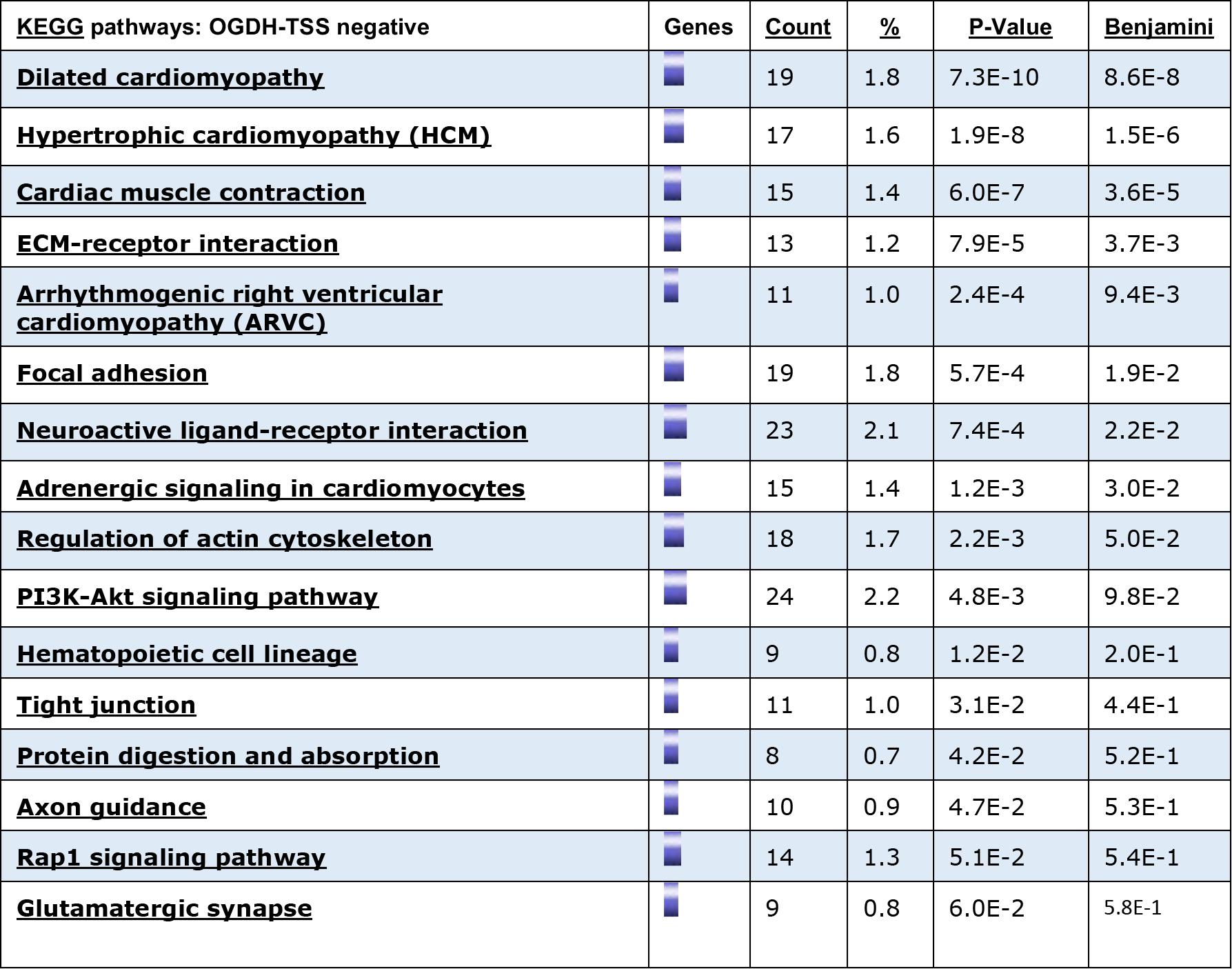
KEGG functional pathway analysis of genes negative for OGDH at TSS (Total OGDH negative genes = 1165 of 11527 expressed genes in the heart)

| <u>KEGG pathways: OGDH-TSS negative</u> | <u>Genes</u> | <u>Count</u> | <u>%</u> | <u>P-Value</u> | <u>Benjamini</u> |
| --- | --- | --- | --- | --- | --- |
| **<u>Dilated cardiomyopathy</u>** |  | 19 | 1.8 | 7.3E-10 | 8.6E-8 |
| **<u>Hypertrophic cardiomyopathy (HCM)</u>** |  | 17 | 1.6 | 1.9E-8 | 1.5E-6 |
| **<u>Cardiac muscle contraction</u>** |  | 15 | 1.4 | 6.0E-7 | 3.6E-5 |
| **<u>ECM-receptor interaction</u>** |  | 13 | 1.2 | 7.9E-5 | 3.7E-3 |
| **<u>Arrhythmogenic right ventricular cardiomyopathy (ARVC)</u>** |  | 11 | 1.0 | 2.4E-4 | 9.4E-3 |
| **<u>Focal adhesion</u>** |  | 19 | 1.8 | 5.7E-4 | 1.9E-2 |
| **<u>Neuroactive ligand-receptor interaction</u>** |  | 23 | 2.1 | 7.4E-4 | 2.2E-2 |
| **<u>Adrenergic signaling in cardiomyocytes</u>** |  | 15 | 1.4 | 1.2E-3 | 3.0E-2 |
| **<u>Regulation of actin cytoskeleton</u>** |  | 18 | 1.7 | 2.2E-3 | 5.0E-2 |
| **<u>PI3K-Akt signaling pathway</u>** |  | 24 | 2.2 | 4.8E-3 | 9.8E-2 |
| **<u>Hematopoietic cell lineage</u>** |  | 9 | 0.8 | 1.2E-2 | 2.0E-1 |
| **<u>Tight junction</u>** |  | 11 | 1.0 | 3.1E-2 | 4.4E-1 |
| **<u>Protein digestion and absorption</u>** |  | 8 | 0.7 | 4.2E-2 | 5.2E-1 |
| **<u>Axon guidance</u>** |  | 10 | 0.9 | 4.7E-2 | 5.3E-1 |
| **<u>Rap1 signaling pathway</u>** |  | 14 | 1.3 | 5.1E-2 | 5.4E-1 |
| **<u>Glutamatergic synapse</u>** |  | 9 | 0.8 | 6.0E-2 | 5.8E-1 |

**Table 8.**
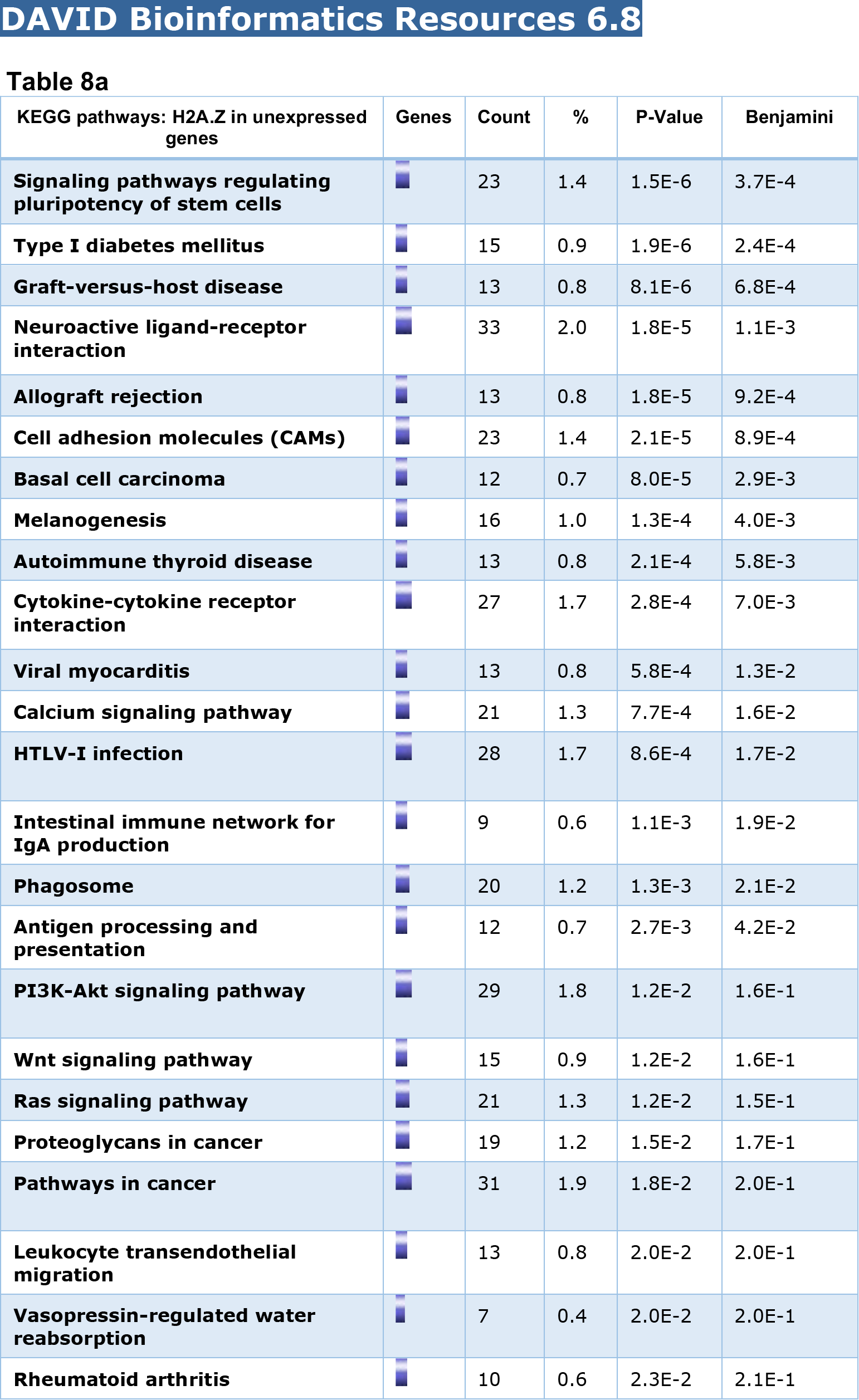
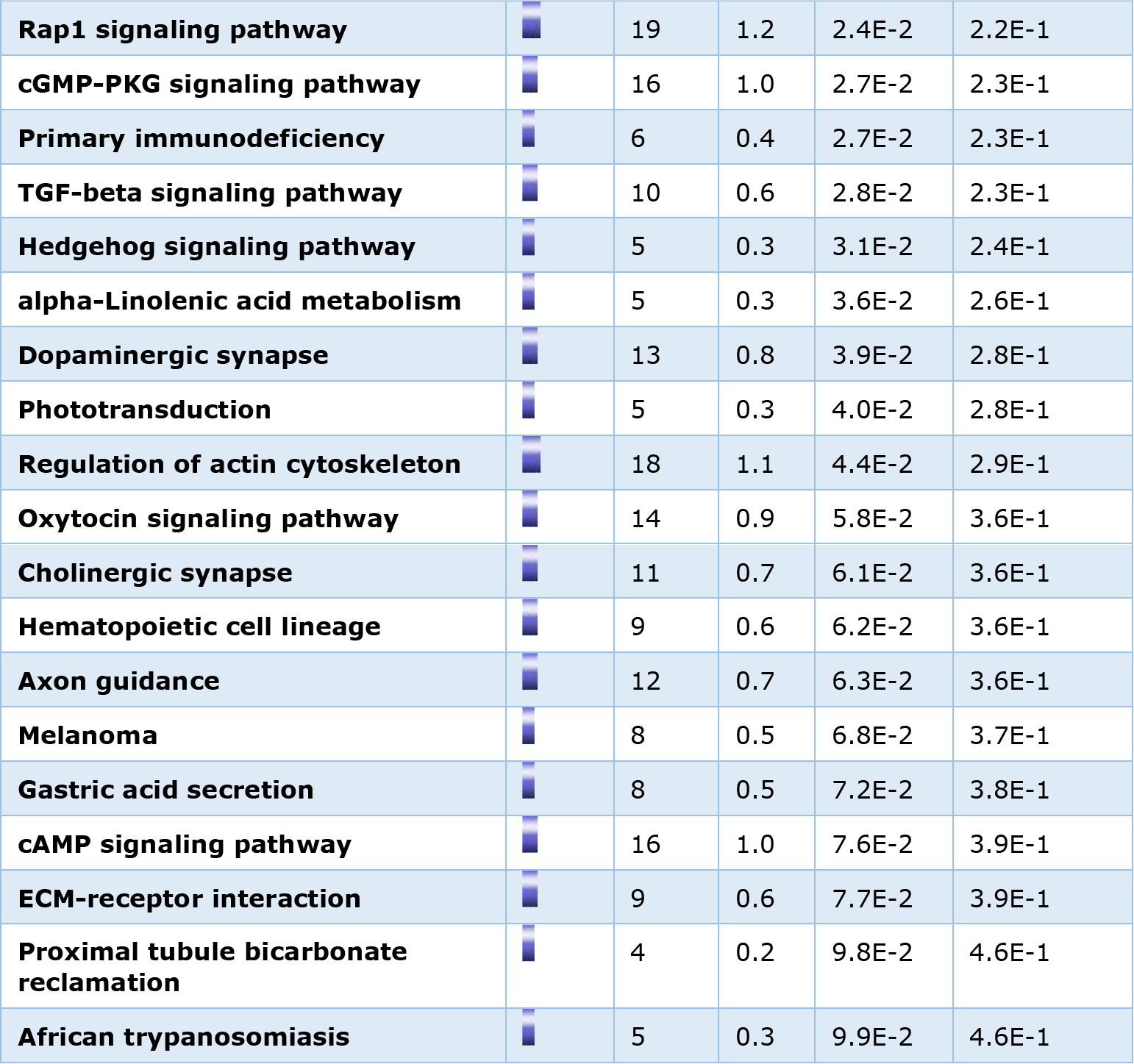
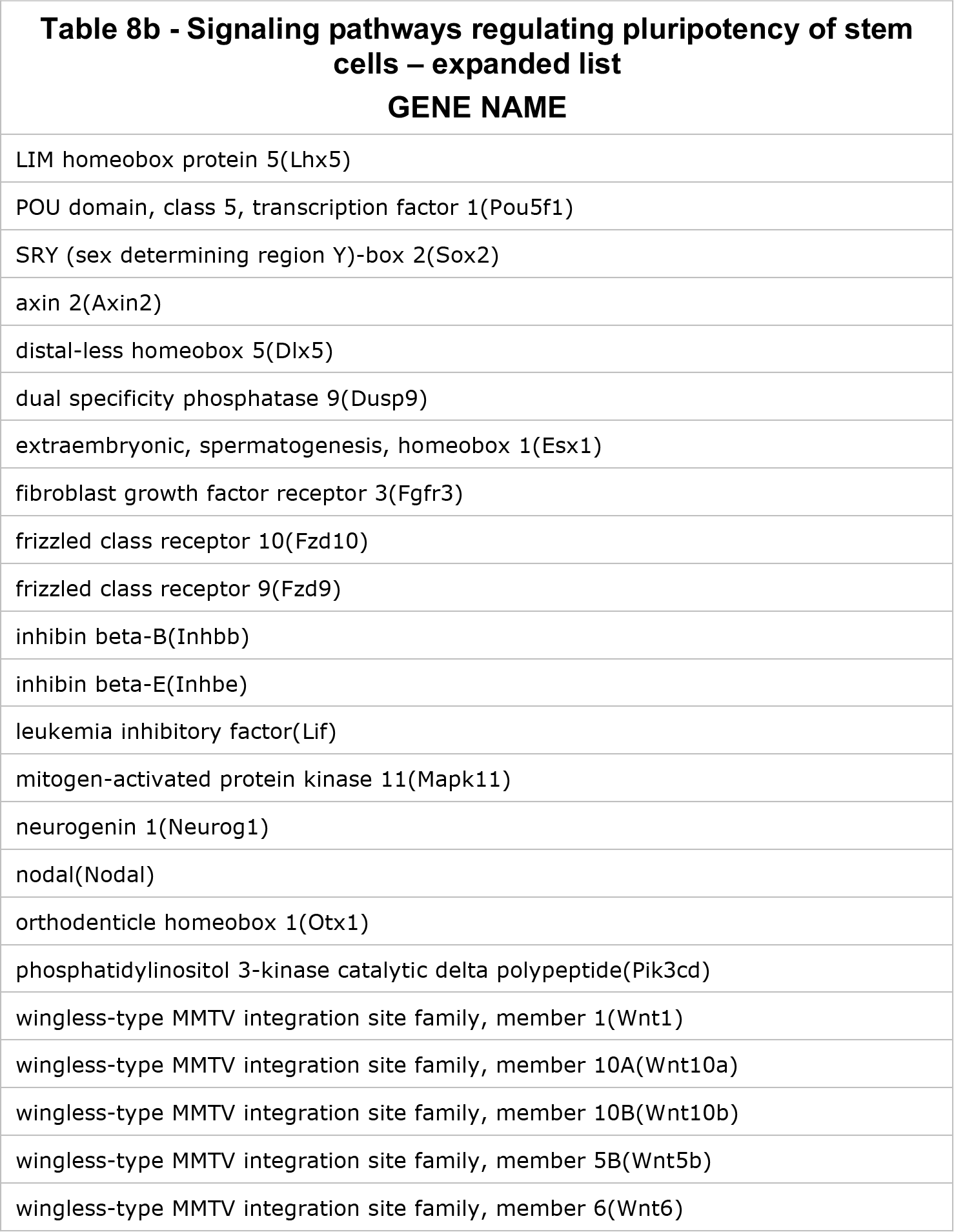
KEGG functional pathway analysis of suppressed genes that are positive for H2A.Z. (Total suppressed gene positive for H2A.Z = 1749)

| KEGG pathways: H2A.Z in unexpressed genes | Genes | Count | % | P-Value | Benjamini |
| --- | --- | --- | --- | --- | --- |
| **Signaling pathways regulating pluripotency of stem cells** |  | 23 | 1.4 | 1.5E-6 | 3.7E-4 |
| **Type I diabetes mellitus** |  | 15 | 0.9 | 1.9E-6 | 2.4E-4 |
| **Graft-versus-host disease** |  | 13 | 0.8 | 8.1E-6 | 6.8E-4 |
| **Neuroactive ligand-receptor interaction** |  | 33 | 2.0 | 1.8E-5 | 1.1E-3 |
| **Allograft rejection** |  | 13 | 0.8 | 1.8E-5 | 9.2E-4 |
| **Cell adhesion molecules (CAMs)** |  | 23 | 1.4 | 2.1E-5 | 8.9E-4 |
| **Basal cell carcinoma** |  | 12 | 0.7 | 8.0E-5 | 2.9E-3 |
| **Melanogenesis** |  | 16 | 1.0 | 1.3E-4 | 4.0E-3 |
| **Autoimmune thyroid disease** |  | 13 | 0.8 | 2.1E-4 | 5.8E-3 |
| **Cytokine-cytokine receptor interaction** |  | 27 | 1.7 | 2.8E-4 | 7.0E-3 |
| **Viral myocarditis** |  | 13 | 0.8 | 5.8E-4 | 1.3E-2 |
| **Calcium signaling pathway** |  | 21 | 1.3 | 7.7E-4 | 1.6E-2 |
| **HTLV-I infection** |  | 28 | 1.7 | 8.6E-4 | 1.7E-2 |
| **Intestinal immune network for IgA production** |  | 9 | 0.6 | 1.1E-3 | 1.9E-2 |
| **Phagosome** |  | 20 | 1.2 | 1.3E-3 | 2.1E-2 |
| **Antigen processing and presentation** |  | 12 | 0.7 | 2.7E-3 | 4.2E-2 |
| **PI3K-Akt signaling pathway** |  | 29 | 1.8 | 1.2E-2 | 1.6E-1 |
| **Wnt signaling pathway** |  | 15 | 0.9 | 1.2E-2 | 1.6E-1 |
| **Ras signaling pathway** |  | 21 | 1.3 | 1.2E-2 | 1.5E-1 |
| **Proteoglycans in cancer** |  | 19 | 1.2 | 1.5E-2 | 1.7E-1 |
| **Pathways in cancer** |  | 31 | 1.9 | 1.8E-2 | 2.0E-1 |
| **Leukocyte transendothelial migration** |  | 13 | 0.8 | 2.0E-2 | 2.0E-1 |
| **Vasopressin-regulated water reabsorption** |  | 7 | 0.4 | 2.0E-2 | 2.0E-1 |
| **Rheumatoid arthritis** |  | 10 | 0.6 | 2.3E-2 | 2.1E-1 |
| **Rap1 signaling pathway** |  | 19 | 1.2 | 2.4E-2 | 2.2E-1 |
| **cGMP-PKG signaling pathway** |  | 16 | 1.0 | 2.7E-2 | 2.3E-1 |
| **Primary immunodeficiency** |  | 6 | 0.4 | 2.7E-2 | 2.3E-1 |
| **TGF-beta signaling pathway** |  | 10 | 0.6 | 2.8E-2 | 2.3E-1 |
| **Hedgehog signaling pathway** |  | 5 | 0.3 | 3.1E-2 | 2.4E-1 |
| **alpha-Linolenic acid metabolism** |  | 5 | 0.3 | 3.6E-2 | 2.6E-1 |
| **Dopaminergic synapse** |  | 13 | 0.8 | 3.9E-2 | 2.8E-1 |
| **Phototransduction** |  | 5 | 0.3 | 4.0E-2 | 2.8E-1 |
| **Regulation of actin cytoskeleton** |  | 18 | 1.1 | 4.4E-2 | 2.9E-1 |
| **Oxytocin signaling pathway** |  | 14 | 0.9 | 5.8E-2 | 3.6E-1 |
| **Cholinergic synapse** |  | 11 | 0.7 | 6.1E-2 | 3.6E-1 |
| **Hematopoietic cell lineage** |  | 9 | 0.6 | 6.2E-2 | 3.6E-1 |
| **Axon guidance** |  | 12 | 0.7 | 6.3E-2 | 3.6E-1 |
| **Melanoma** |  | 8 | 0.5 | 6.8E-2 | 3.7E-1 |
| **Gastric acid secretion** |  | 8 | 0.5 | 7.2E-2 | 3.8E-1 |
| **cAMP signaling pathway** |  | 16 | 1.0 | 7.6E-2 | 3.9E-1 |
| **ECM-receptor interaction** |  | 9 | 0.6 | 7.7E-2 | 3.9E-1 |
| **Proximal tubule bicarbonate reclamation** |  | 4 | 0.2 | 9.8E-2 | 4.6E-1 |
| **African trypanosomiasis** |  | 5 | 0.3 | 9.9E-2 | 4.6E-1 |

| **Table 8b - Signaling pathways regulating pluripotency of stem cells – expanded list** |
| --- |
| **GENE NAME** |
| LIM homeobox protein 5(Lhx5) |
| POU domain, class 5, transcription factor 1(Pou5f1) |
| SRY (sex determining region Y)-box 2(Sox2) |
| axin 2(Axin2) |
| distal-less homeobox 5(Dlx5) |
| dual specificity phosphatase 9(Dusp9) |
| extraembryonic, spermatogenesis, homeobox 1(Esx1) |
| fibroblast growth factor receptor 3(Fgfr3) |
| frizzled class receptor 10(Fzd10) |
| frizzled class receptor 9(Fzd9) |
| inhibin beta-B(Inhbb) |
| inhibin beta-E(Inhbe) |
| leukemia inhibitory factor(Lif) |
| mitogen-activated protein kinase 11(Mapk11) |
| neurogenin 1(Neurog1) |
| nodal(Nodal) |
| orthodenticle homeobox 1(Otx1) |
| phosphatidylinositol 3-kinase catalytic delta polypeptide(Pik3cd) |
| wingless-type MMTV integration site family, member 1(Wnt1) |
| wingless-type MMTV integration site family, member 10A(Wnt10a) |
| wingless-type MMTV integration site family, member 10B(Wnt10b) |
| wingless-type MMTV integration site family, member 5B(Wnt5b) |
| wingless-type MMTV integration site family, member 6(Wnt6) |

One of the unresolved issues that needs to be thoroughly investigated, is the exact stoichiometry of the enzymes and the interdependence of their bindings and functions, which is apt to determine the concentrations of the metabolites that are produced/consumed at the TSSs. Accordingly, we expect that the composition of these enzymatic complexes to define the underlying histone modifications and the responsiveness of the genes’ expression to oxidative or metabolic cues, while any perturbation may result in pathogenesis. This is underscored by the fact that while ACAA2 fully overlaps with ODGH at TSSs, the precise pattern of binding and their responsiveness to growth stimuli in the heart are distinct. For instance, in the *Ubc* gene, the two start sites exhibit differential binding to ACAA2 and OGDH, particularly, during growth, when OGDH shows a decrease in abundance at the first TSS accompanied by an increase at the second TSS, whereas, the ACAA2 assembled at the first TSS remains unchanged (Fig. 5). Meanwhile, H2A.Z abundance does not vary significantly. Another incompletely resolved matter, is how these enzymes are imported into the nucleus. We were able to confirm that, at least, ACAA2 and OGDH harbor NLSs that are required for their nuclear import (Fig. 3 and supplementary Fig 3S), however, it remains necessary to investigate the other enzymes.

In summation, the findings add a new level of understanding to the intricacies of the transcriptional machinery and its regulation by metabolism.

## MATERIALS AND METHODS

### Animal care

All animal procedures used in this study are in accordance with US National Institute of Health *Guidelines for the Care and Use of Laboratory Animals (No. 85-23)*. All protocols were approved by the Institutional Animal Care and Use Committee at the Rutgers-New Jersey Medical School.

### H2A.Z rapid immunoprecipitation mass spectrometry of endogenous proteins (RIME)

Male C57/Bl, 12 wk-old mice, 10 each, were subjected to a sham or transverse aortic constriction (TAC) procedure. After 1 wk, the hearts were isolated, pooled for each condition, and sent to Active Motif for RIME analysis by anti-H2A.Z (Active Motif, cat # 39943).

#### DATABASE SEARCHING (Active Motif)

All MS/MS samples were analyzed using X! Tandem (The GPM, thegpm.org; version X! Tandem Alanine (2017.2.1.4)). X! Tandem was set up to search the UP_mouse_CrapE2F1_rev database (unknown version, 106444 entries) assuming the digestion enzyme trypsin. X! Tandem was searched with a fragment ion mass tolerance of 20 PPM and a parent ion tolerance of 20 PPM. Glu->pyro-Glu of the N-terminus, ammonia-loss of the N-terminus, gln-pyro-Glu of the n-terminus deamidated of asparagine and glutamine, oxidation of methionine and tryptophan and dioxidation of methionine and tryptophan were specified in X! Tandem as variable modifications. Each sample was analyzed twice by MS/MS.

#### CRITERIA FOR PROTEIN IDENTIFICATION (Active Motif)

Scaffold (version Scaffold_4.8.4, Proteome Software Inc., Portland, OR) was used to validate MS/MS based peptide and protein identifications. Peptide identifications were accepted if they could be established at greater than 50.0 % probability by the Scaffold Local false discovery rates (FDR) algorithm. Peptide identifications were also required to exceed specific database search engine thresholds. X! Tandem identifications required at least. Protein identifications were accepted if they could be established at greater than 5.0% probability to achieve an FDR less than 5.0% and contained at least 1 identified peptide. Protein probabilities were assigned by the Protein Prophet algorithm ^34^. Proteins that contained similar peptides and could not be differentiated based on MS/MS analysis alone were grouped to satisfy the principles of parsimony. Proteins sharing significant peptide evidence were grouped into clusters.

#### ANALYSIS

The total spectra counts for all genes in the three samples, were normalized to the corresponding spectra count of the rabbit IgG (control or anti-H2A.Z) used for the immunoprecipitation, which was detected in each sample. Fold enrichment of total spectra for sham : IgG and TAC : IgG, for each gene, was calculated. Seventy-three of these had a ≥ 2-fold enrichment, those are shown in figure 1 and supplementary figure 1S.

### Culturing rat neonatal cardiac myocytes

Cardiac myocytes were cultured as described in our previous reports ^35^. Briefly, hearts were isolated from 1 day old of Sprague-Dawley rats. After dissociation with collagenase, cells were subjected to Percoll gradient centrifugation followed by differential pre-plating for 30 min to enrich for cardiac myocytes and deplete non-myocyte cells. Myocytes were cultured in Dulbecco’s Modified Eagle’s medium supplemented with 10% fetal bovine serum (FBS). All experiments were initiated after a 24 h culturing period.

### Culturing mouse adult cardiac myocytes

Adult cardiac myocytes were isolated and cultured from C57/Bl mice (8-9 wks old), according to the protocol described by Ackers-Johnson et al. ^36^. Briefly, mice were anesthesia with Ketamine/Xylazine/Acepromazine (65/13/2 mg/kg) by intraperitoneal injection. The mouse chest cage was then opened, the ascending aorta clamped, and both descending aorta and inferior vena cava cut. First, ethylenediaminetetraacetic acid (EDTA) is injected into the base of the right ventricle, the heart is then transferred into a petri dish, and a second EDTA injection is administered into the left ventricular wall above the apex. Following this, the cells are dissociated using collagenase, and the rod-shaped myocytes are differentially separated by gravity, where calcium is re-introduced. The cells are plated on laminin-coated dishes or glass slides in M199 medium with 5 % FBS for 1 hour, after which the FBS is replaced with 0.1% bovine serum albumin for longer culturing periods.

### Human iPSC-derived cardiac myocyte cultures

Cardiac myocytes derived from human iPSCs were purchased from Cellular Dynamics International and cultured as recommended by the manufacturer.

### Colon cancer cell culture

SW620 were purchased from the American Type Culture Collection (ATCC). Cells were cultured in Leibovitz’s (Gibco) medium with 10% FBS and maintained in a CO2 free incubator.

### Human haploid HAP1 and ∆H2A.Z-HAP1 cell cultures

HAP1 and H2A.Z knockout HAP1 (∆H2A.Z-HAP1) cell lines were purchased from Horizon Discovery and cultured in Iscove’s Modified Dulbecco’s Medium with 10% FBS, according to the company’s protocol. These cells are fibroblast-like derived from human male chronic myelogenous leukemia (CML) cell line KBM-7. The ∆H2A.Z-HAP1 was generated by CRISPER/Cas, creating a 2bp deletion in exon 3 of H2A.Z. The cells are viable but have a much slower proliferation rate than the parent cell-line.

### Construction of GFP fusion proteins

The plasmids harboring cDNAs of turbo-GFP (tGFP), OGDH (NM_002541), ACAA2 (NM_177470), and IDH2 (NM_002168) were purchased from Origene. The cDNA of the latter three genes were then sub-cloned, in-frame, upstream of tGFP, and the fusion cDNA was subsequently sub-cloned into the pDC316 shuttle plasmid vector (Microbix), which was used to generate recombinant adenoviral vectors via homologous recombination.

### Harvesting and immunostaining mouse embryos

Embryos were dissected at E10.5 in ice-cold phosphate-buffered saline (PBS), fixed with 4% paraformaldehyde at 4°C overnight, and then washed with ice-cold PBS. Embryos were then embedded into optimal cutting temperature matrix and sectioned in sagittal orientation. For immunofluorescence, sections were incubated with blocking buffer: 5% donkey serum (Sigma, cat # D9663) diluted in PBS containing 0.05% Tween-20 (PBST), for 30 minutes at room temperature.

Sections were then incubated with primary antibodies at 4°C overnight: OGDH (Cell Signaling Technology, cat # 26865, 1:100 dilution) and alpha-cardiac actin (Sigma, cat # A9357, 1:300 dilution) diluted in the blocking buffer. Slides were then washed in PBS and incubated with 4′,6-diamidino-2-phenylindole (DAPI) to mark nuclei and with secondary antibodies diluted 1:300 in the blocking buffer for 2 hours at room temperature. 2° antibodies were donkey anti-rabbit-Alexa 488 (Invitrogen, cat # A21206) and donkey anti-mouse-Alexa 555 (Invitrogen, cat # A31570).

### Transverse aortic constriction (TAC) in mice

This was performed as described in our previous reports ^37, 38^. Briefly, a 7-0 braided polyester suture was tied around the transverse thoracic aorta, against a 27-gauge needle, between the innominate artery and the left common carotid artery. Control mice were subjected to a sham operation involving the same procedure, minus the aortic constriction.

### Echocardiography and doppler

This was performed as described in our previous reports ^37, 38^. Briefly, transthoracic echocardiography was performed using the Vevo 770 imaging system (Visual Sonics, Inc.) with a 707B-30MHz scanhead, encapsulated, transducer. Electrocardiographic electrodes were taped to the four paws, then one dimensional (1D) M-mode and 2D B-mode tracings were recorded from the parasternal short-axis view at the mid papillary muscle level. In addition, pulse-wave Doppler was used to measure blood flow velocity and peak gradient pressure in the aorta. For analysis, we used the Vevo 770 Software (Vevo 770, Version 23), which includes: analytic software package for B-Mode (2D) image capture and analysis; cine loop image capture, display, and review; software analytics for advanced measurements and annotations; and physiological data on-screen trace.

### Construction and delivery of recombinant adenovirus (Ad) vector

Recombinant adenoviral vectors were constructed, propagated, purified, and tittered as described in our previous reports ^39–41^. Briefly, short hairpin RNAs (shRNAs) were cloned into pDC311 shuttle (Microbix Biosystems Inc.), downstream of a U6 promoter. These were transfected with the replication-defective Ad5 viral DNA backbone into 293HEK cells, in which a recombination reaction introduces the DNA insert into the viral DNA. Single virus plaques were amplified in 293HEK cells, purified on a CsCl_2_ gradient, dialyzed, and tittered on 293HEK cells with agarose overlay. Ad vectors were constructed with the following inserts, shRNA targeting OGDH - ccagccactggcaacaagaaTTCAAGAGAAttcttgttgccagtggctggTTTTTT, shRNA targeting H2A.Z - gtcacttgcagcttgctataTTCAAGAGAAtatagcaagctgcaagtgacTTTTTT, and shRNA targeting ACAA2 - cagttcttgtctgttcagaaTTCAAGAGAAttctgaacagacaagaactgTTTTTT, or a nonsense control shRNA gaaccgagcccaccagcgagcTTCAAGAGAAgctcgctggtgggctcggttcTTTTTT, the shaded areas are the loop sequence and the terminal 6xT’s is the stop signal for the U6 promoter used for the expression of these shRNAs. Cardiac myocytes were infected with 10-30 multiplicity-of-infection (moi) of the viruses for 24 h or 48 h, as indicated in the figure legends.

### Subcellular fractionation and Western blotting

Proteins were fractionated using the subcellular protein fractionation kit (Thermo Fisher, cat # 78840), according to the manufacturer’s protocols. The cellular fractions were separated on a 4% to 12% gradient SDS-PAGE (Criterion gels, Bio-Rad) and transferred to nitrocellulose membrane. The antibodies used include: anti-turboGFP (Origene, cat # TA150041), -PDHA1 (Cell Signaling Technology, cat # 3205), -IDH2 (Cell Signaling Technology, cat # 56437), -OGDH (E1W8H, Cell Signaling Technology, cat # 26865), -ACAA2 (Origene Technologies, cat # TA506126), -H2A.Z (Active Motif, cat # 39943), -H3 (Active Motif, cat # 61476), -H3 pan-acetyl (Active Motif, cat # 39140), -H3K27 di-and tri-methyl (H3K27me2/3, Active Motif, cat # 39538), -nuclear pore glycoprotein p62 (NUP62, US Biological, cat # USB326547), -hydroxyacyl-CoA dehydrogenase trifunctional multienzyme complex subunit alpha (HADHA, Abcam, cat # ab203114), -pyruvate dehydrogenase kinase 1 (PDK1, Novus Biologicals, cat # 100-2383), - transcription factor II B (TFIIB, Cell Signaling Technology, cat # 4169), -H4K12-succinyl (H4K12suc, Epigentek, cat # A70383), -H2A (Active Motif, cat # 35951), -H4 (Upstate, cat # 07-108), -H3K9 mono-, di-, tri-methyl (H3K9me1/2/3, Active Motive, cat # 38241), -AKT1 (Cell Signaling Technology, cat # 9272), -voltage-dependent anion-selective channel 1 (VDAC1, Genscript, cat # A01419), -succinate dehydrogenase complex, subunit A (SDHA, Thermo Fisher, cat # 459200), -succinate dehydrogenase complex, subunit b (SDHB, Santa Cruz Biotechnology, cat # sc-25851), and -RNA pol II (Abcam, cat # ab5095). The Western blot signals were detected by the Odyssey imaging system (LI-COR).

### Immunocytochemistry

Cells were seeded on glass chamber slides coated with fibronectin for neonatal myocytes and hiPSC-CM, or with laminin for adult myocytes, fixed with 3% formaldehyde / 0.3% triton x-100, then incubated with antibodies (1:100) or phalloidin (in Tris-buffered saline with 1% bovine serum albumin), washed and mounted using Prolong Gold anti-fade with DAPI (Molecular Probes). The antibodies included: anti-PDHA1, -IDH2, -OGDH (Cell Signaling Technology, same as those listed for the Western blotting, above), as well as, anti-OGDH (Sigma, cat # HPA019514), and anti-ACAA2 (Origene Technologies, cat # TA506126). The slides were imaged using Nikon A1R laser scanning confocal microscope with Plan Apo 60x objective.

### ChIP-Seq (Active Motif) and data analysis

Mice were subjected to transverse aortic constriction or a sham operation. After 7 days, cardiac function and structure were assessed by echocardiography, before isolation of the hearts. The hearts were then analyzed by ChIP using the following antibodies: anti-RNA pol II (Abcam, cat # ab5095), -H2A.Z (Active Motif, cat # 39113), -H3K9-acetyl (H3K9ac) (Active Motif, cat # 39918), -TFIIB (Santa Cruz Biotechnology, cat # sc-225), -cyclin-dependent kinase 9 (CDK9, Santa Cruz Biotechnology, cat # sc-8338), -ANP32e (Abcam, cat # ab5993), -ACAA2 (Origene Technologies, cat # TA506126), and -OGDH (E1W8H, Cell Signaling Technology, cat # 26865), followed by next generation sequencing (Active Motif). We have previously reported the results of our ChIP-Seq for RNA pol II ^42^, H3K9ac ^42^, TFIIB ^35^, H2A.Z ^17^, and ANP32E ^17^, and, thus, are not further described here. Briefly, ChIP libraries were sequenced using NextSeq 500, generating 75-nt sequence reads that are mapped to the genome using BWA algorithms. The reads/tags were extended *in silico* by 150-250 bp at their 3’end (fragments), the density of which is determined along the genome, divided in 32 nt bins, and the results saved in bigWig and BAM (Binary Alignment/Map) files. Fragment peaks were identified using MACS, which identifies local areas of enrichment of Tags, defined as ‘intervals’, while overlapping intervals are grouped into ‘Merged Regions’. The locations and proximities to gene annotations of intervals and active regions are defined and compiled in Excel spreadsheets, which include average and peak fragment densities. Regarding tag normalization and input control, the sample with the lowest number of tags is used for normalization of all samples, while the input is used to identify false positive peaks. The statistics for ACAA2 and OGDH ChIP-Seq results, including total number of reads, peaks, empirical FDR, and peak calling parameters are listed in supplementary Table 1.

In addition, we separately analyzed the fragment densities by gene region, where the average value (Avg Val) of fragment densities at the TSS (−1000 to +1000) and in-gene/gene body (+1000 to 3’ end) regions for all genes were calculated separately. Subsequently, we used these values to sort genes according to TSS-pol II, -ACCA2, or -OGDH, occupancy.

### ChIP-Seq analysis software

The heatmaps, curves, and histograms shown in figure 4a-c, were generated using EaSeq ^43^. Images of sequence alignments of fragments across chromosomal coordinates were generated using the Integrated Genome Browser (Fig. 6-7) ^44^.

### Statistical analysis

The significance of differences between 2 experimental groups was calculated using T-test (equal variance, 2-tailed), where *p* < 0.05 was considered significant.

## Computational Resources

The RNA polymerase II AND H3K9ac ChIP-Seq data (accession: GSE50637), the TFIIB ChIP-Seq data (accession: GSE56813) and H2A.Z and ANP32E ChIP-Seq data (accession: GSE104702) are available in the Gene Expression Omnibus Datasets. The ChIP-Seq data for ACAA2, OGDH, and Cdk9 will be deposited in GEO with a private link for the reviewers, which will be made public upon acceptance.

Integrated genome browser ^44^ can be downloaded free at: http://bioviz.org/igb/, and Easeq ^43^ can be downloaded free at: http://easeq.net/

## ACKNOWLEDGEMENT

We thank Dr. Junichi Sadoshima, Chairman of the Department of Cell Biology and Molecular Medicine, Rutgers University, for his support.

---

**Figure 1S.**
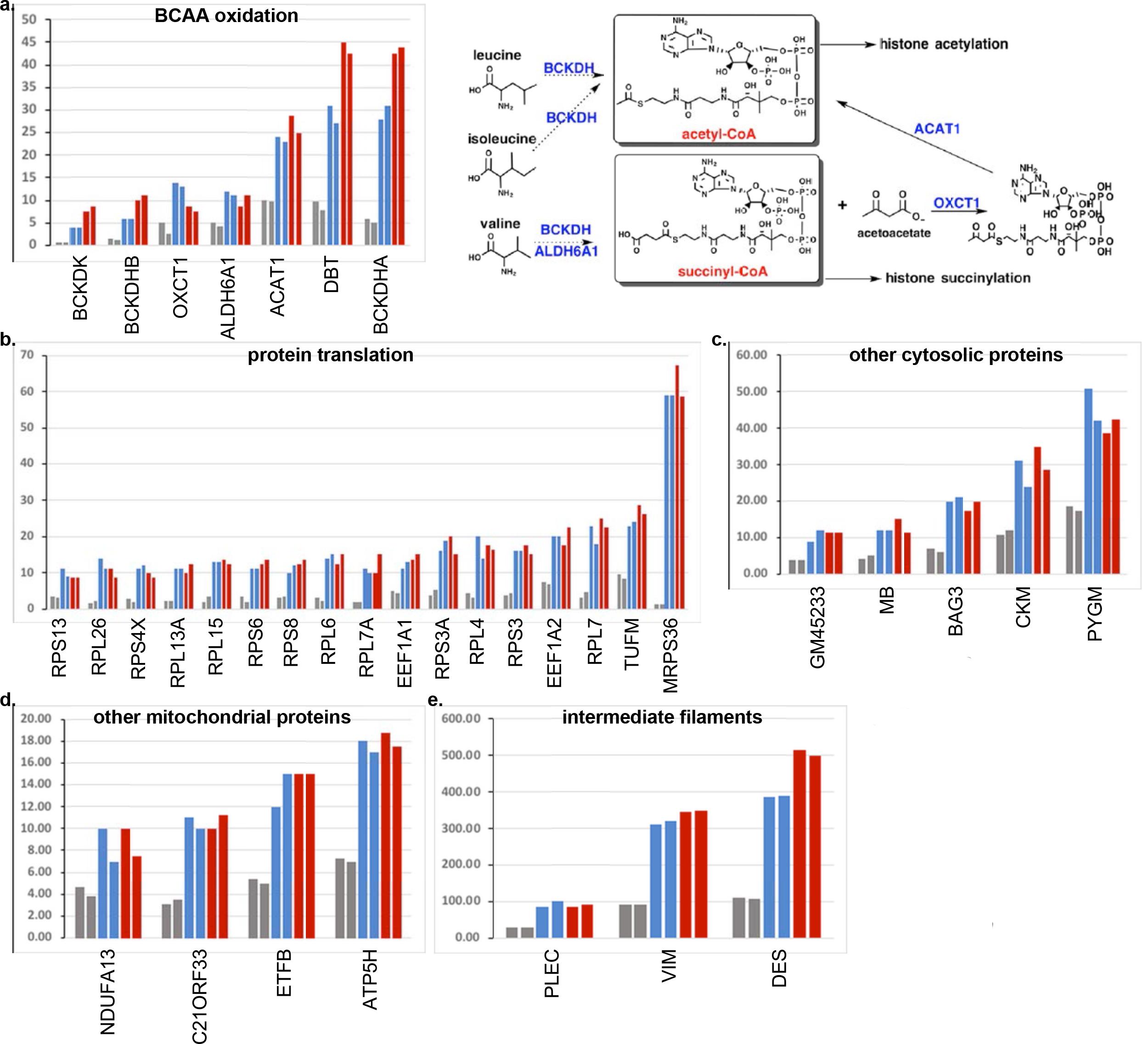
Additional proteins that co-precipitate with H2A.Z-bound chromatin in the RIME assay. ChIP with anti-H2A.Z or a control IgG was performed on nuclei from a pool of 10 hearts each, from mice subjected to a sham or a transverse aortic constriction operation. The ChIP complex was then subjected to MS/MS, each sample analyzed twice. a.-e. Total spectra identified in the IgG, sham, and TAC samples were plotted. These include those with a cutoff of more than 2-fold enrichment v. control IgG, after normalization to the IgG C chain region spectra detected in each sample. Data were grouped as **a**. branched-chain amino acid (BCAA) oxidation enzymes, **b.** protein translation proteins, **c.** other cytosolic proteins, **d.** other mitochondrial proteins, and **e.** intermediate filament proteins. The enzymes 1 in a. are represent in blue in the BCAA oxidation pathway illustrated on its right.

**Figure 2S.**
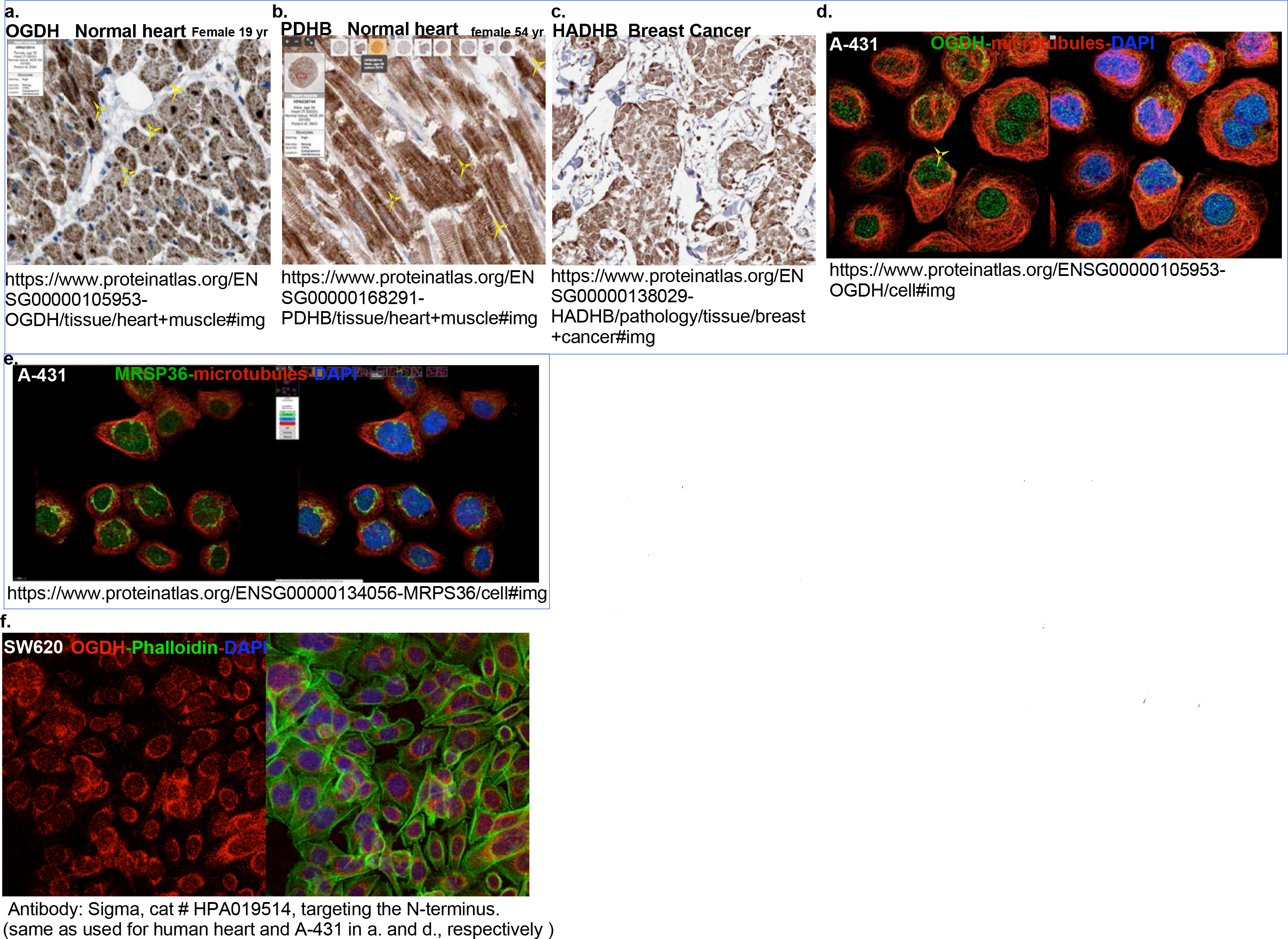
Images from the human protein atlas project (www.proteinatlas.org). The Human Protein Atlas is a Swedish-based project that includes antibody-based imaging of human tissue and cell lines, and is open access for scientists allowing free use of the data, given that it is properly cited [Ref. 18]. Shown are images from that project that include normal human heart sections immuno-stained with **a.** anti-OGDH and **b.** anti-PDHB, **c.** human breast cancer sections immuno-stained with anti-HADBH, **d.** A431 cells immuno-stained with anti-OGDH, and **e.** A-431 cells immuno-stained with anti-MRSP36. Direct links to the web pages are listed beneath each image. Note, different antibodies had differential affinities to the nuclear v. mitochondrial form of a given protein, as demonstrated in our data **f.** We also stained the human colon cancer cell line SW620 with the a second OGDH antibody that targets the N-terminus v. the C-terminus, used in the main figure 2, for validation of the data.

**Figure 3S.**
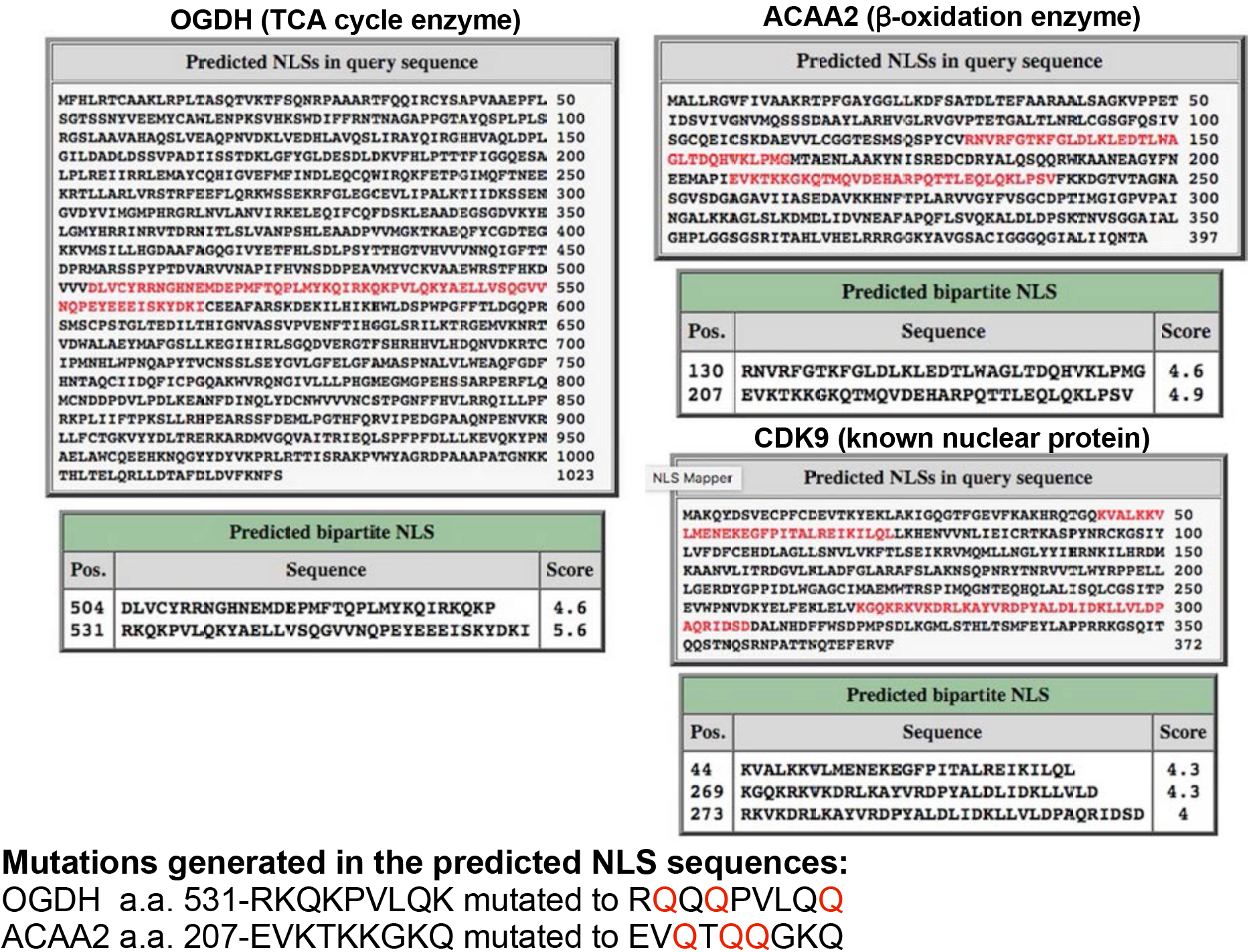
Prediction of importin α-dependent nuclear localization signal (NLS) using cNLS Mapper. The images show the output of the NLS prediction results for OGDH, ACAA2, and CDK9 (used as a positive control), by cNLS Mapper, a free web-based predictions of software: http://nls-mapper.iab.keio.ac.jp/cgi-bin/NLS_Mapper_form.cgi. The predicted NLS is indicated by red lettering. The scoring system is such that a protein with a score of 8, 9, or 10 is exclusively nuclear; 7 or 8 is partially nuclear; a score of 3, 4, or 5 is both nuclear and cytoplasmic; and a score of 1 or 2 is cytoplasmic. The mutations generated in the OGDH and ACAA2 predicted NLS are shown at the bottom, where the substituted amino acids are indicated in red.

**Figure 4S.**
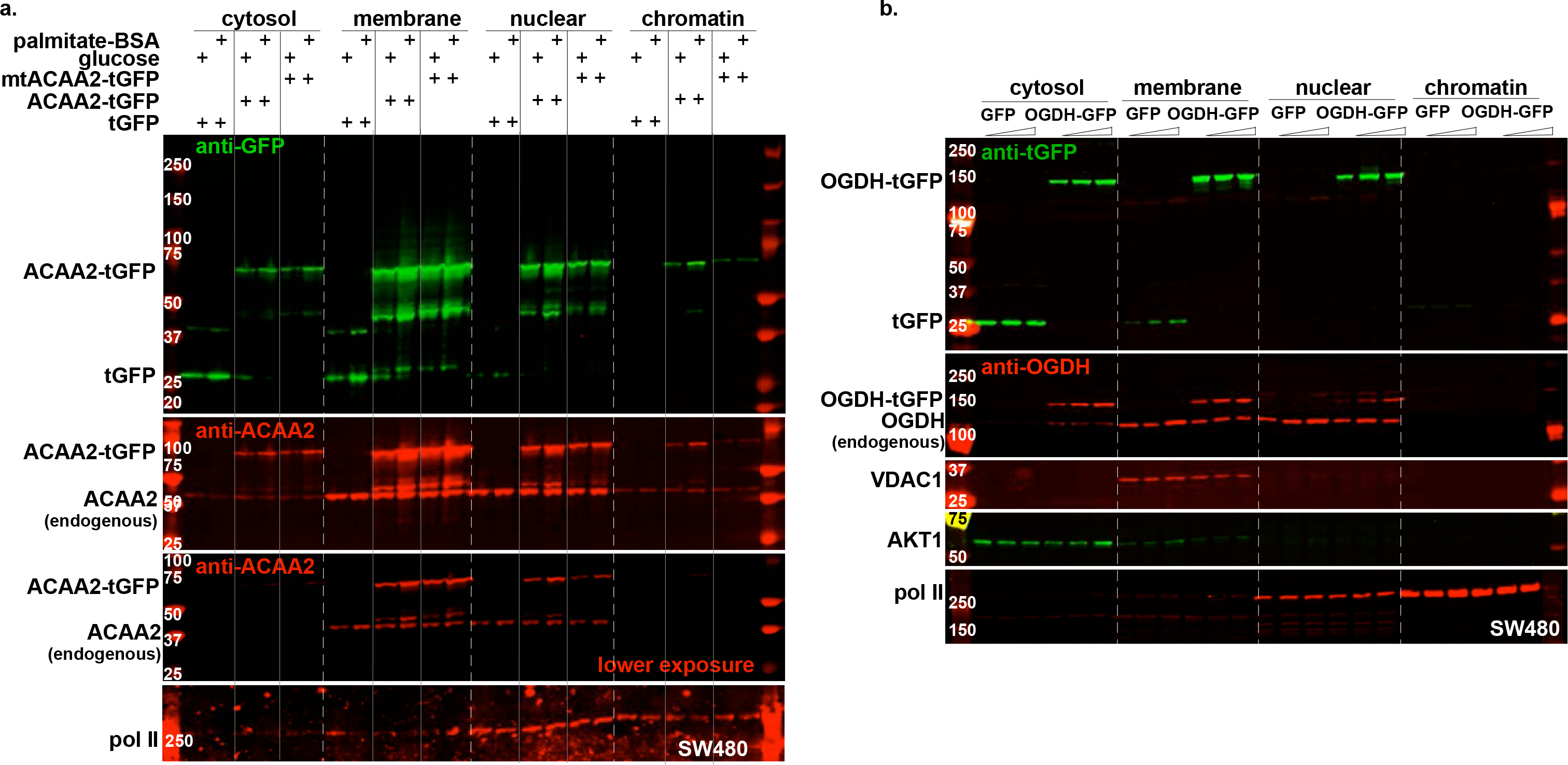
Nuclear localization of metabolic enzymes confirmed by tGFP-fusion proteins and NLS mutation. SW480 human colon cancer cell were infected with a 10-30 moi of recombinant adenoviruses harboring turbo-GFP (tGFP) or **a.** wt ACAA-tGFP or an NLS mutant (mtACAA2-tGFP), **b.** wt OGDH. In **a.,** the cells were incubated in either glucose-containing (fatty acid and serum-free) or in palmitate-BSA (glucose-free and serum-free) medium, as indicated at the top of the lanes. After 18 h, the cellular protein/organelles were fractionated into cytosol (cyto), mitochondrial and membrane (mito), nuclear (nuc), and chromatin-bound (chrom) protein fractions that were then analyzed by Western blotting for the proteins listed on the left of each panel. The fusion proteins were detected by anti-GFP (upper panels, a-b) and anti-ACAA2 or anti-OGDH (second panels, a-b), which also detect the endogenous proteins. AKT1, VDAC1, Pol II, were immunodetected for their use as internal controls for the corresponding cell fractions: cytosol, mitochondria and, nuclear and chromatin, respectively.

**Figure 5S.**
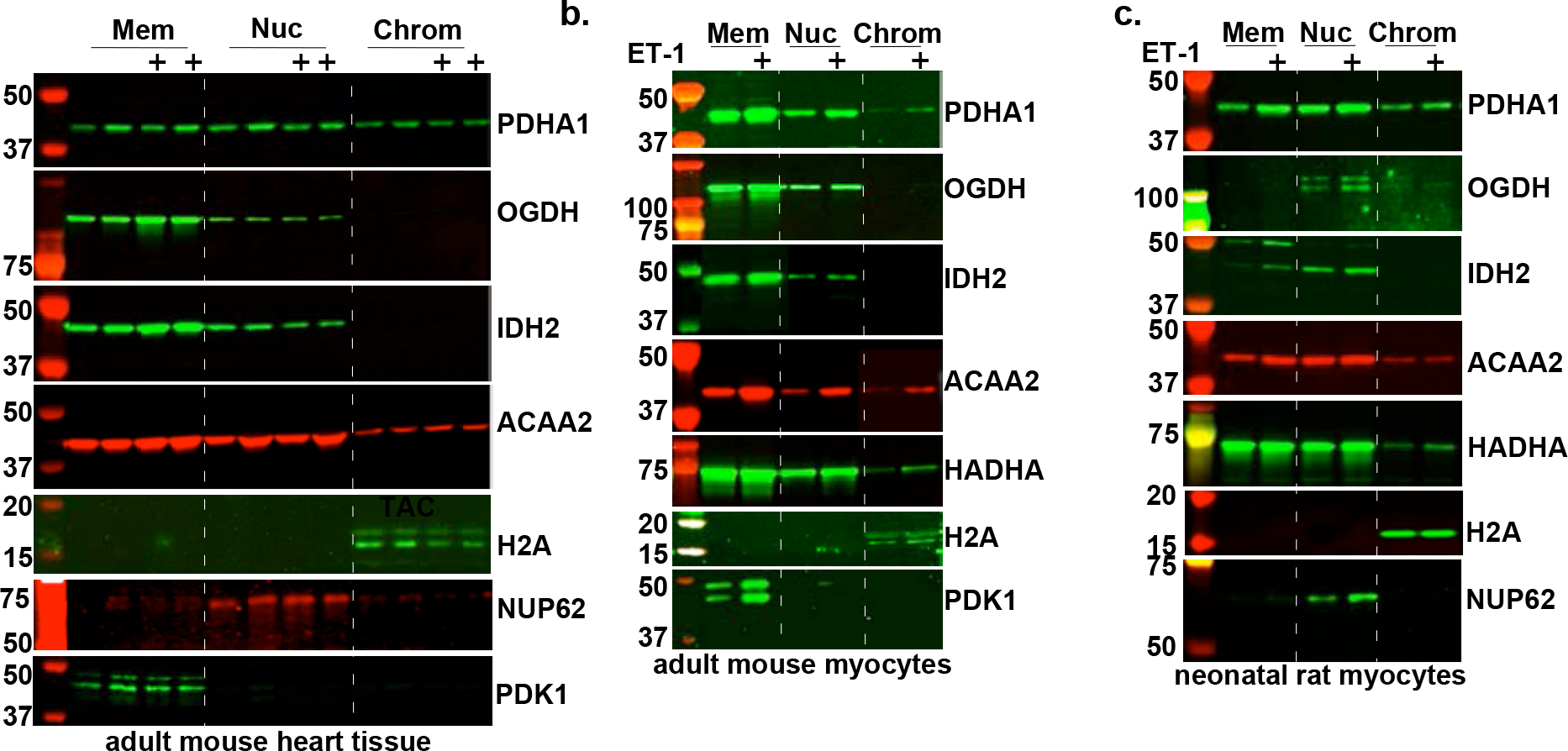
Nuclear localization of metabolic enzymes in mouse heart and isolated cardiac myocytes, confirmed by Western blotting. Cellular organelles were extracted from a. mouse sham or TAC hearts, b. isolated mouse adult cardiac myocytes (mACM), and c. rat neonatal cardiac myocytes. These were fractionated into membrane, including mitochondria (Mem), nuclear (Nuc), and chromatin-bound (Chrom, no crosslinking applied), using a combination of differential lysis and sequential centrifugation. The protein extracted from each of these fractions was analyzed by Western blotting for the genes indicated on the right of each panel (n=3, each).

**Figure 6S.**
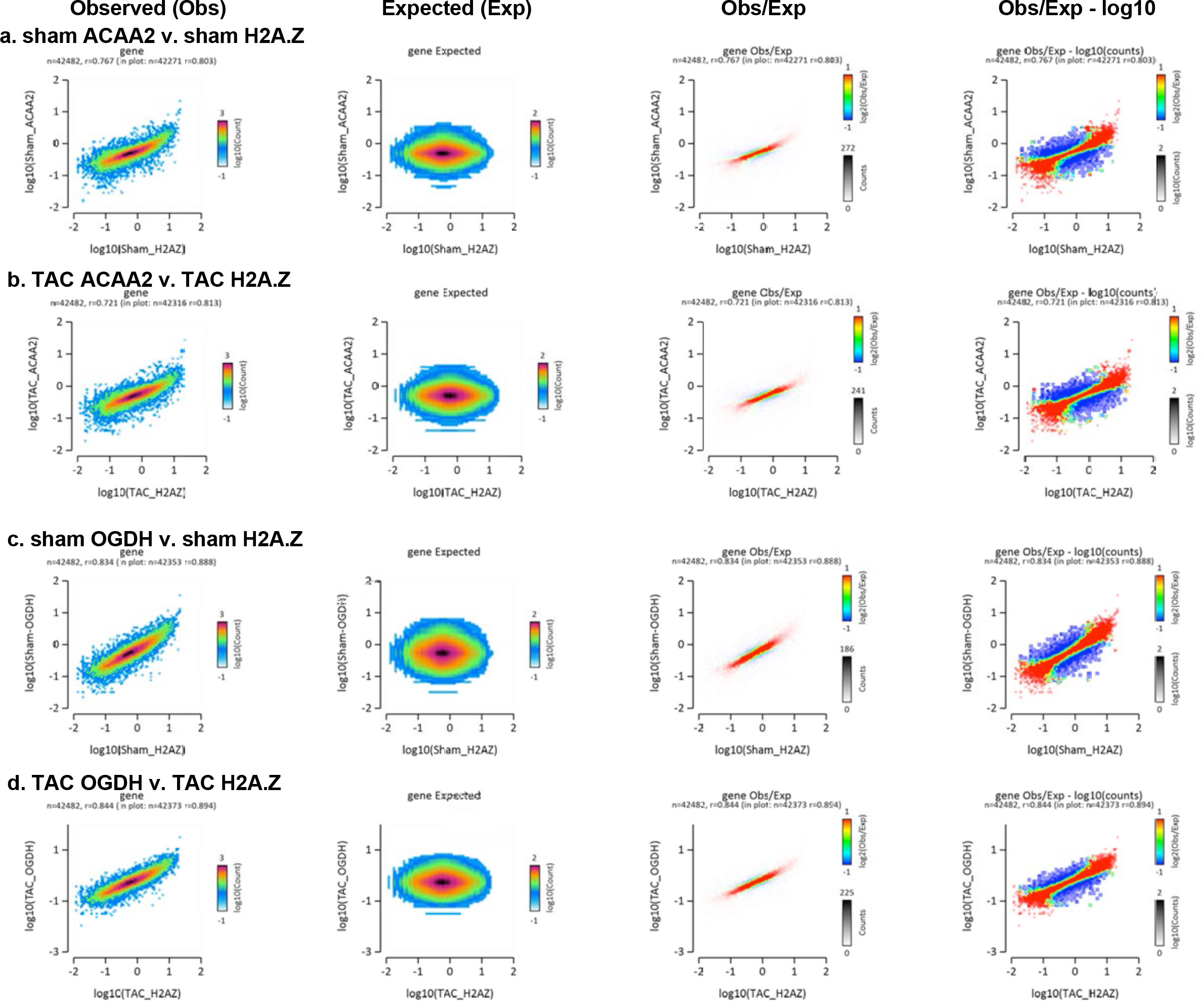
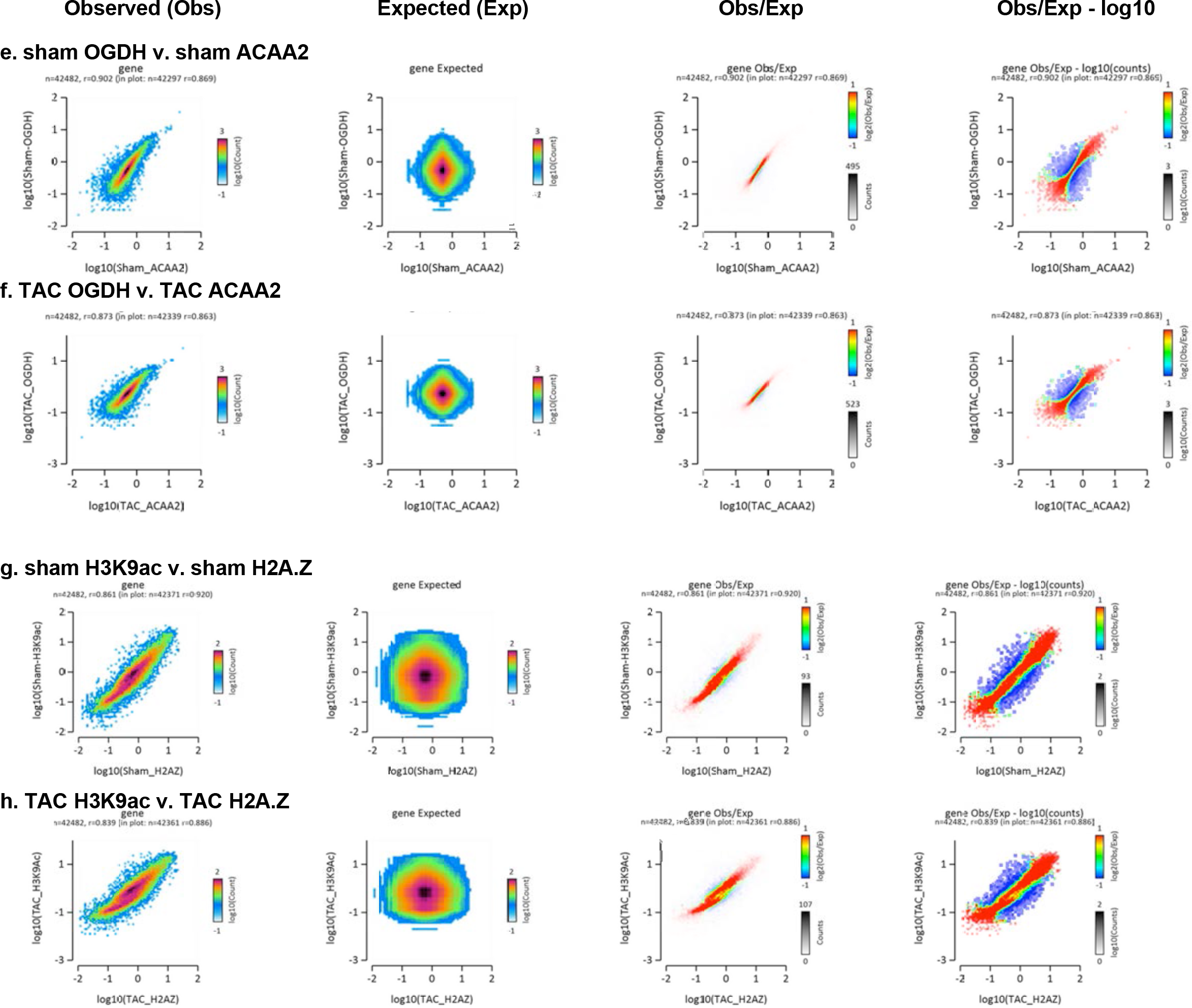
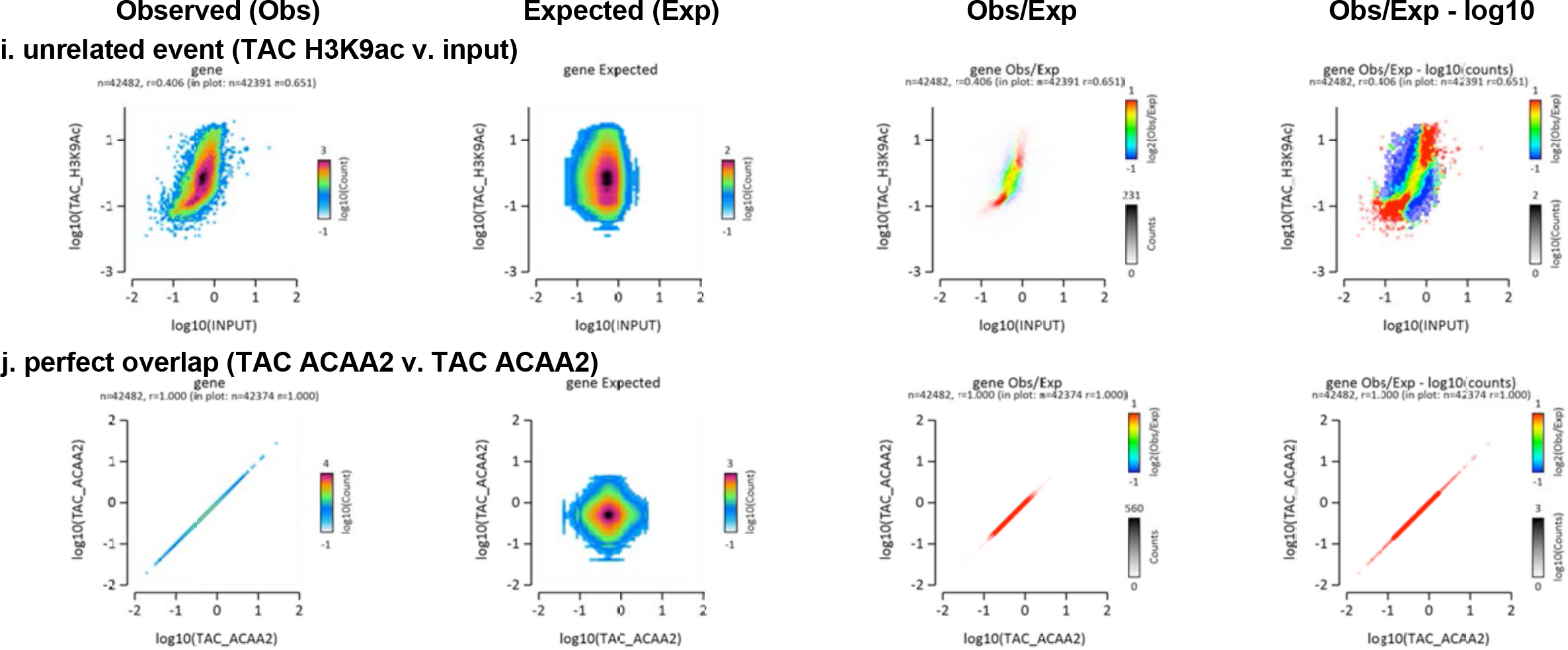
The association of ACAA2 and OGDH with chromatin overlaps with H2A.Z at transcription start sites. Histograms showing the distribution of fragments calculated from their overall frequencies in the ChIP-Seq of H2A.Z (X-axis) v. ACAA2 or OGDH (Y-axis), and of ACAA2 (X-axis) v. OGDH (Y-axis), over the length of the gene and including −2000 bp upstream of the TSS, as labeled. The X and Y-axes were segmented into 75 bins, and the number of fragments within each bin was counted, color coded, and plotted. The bar to the right of the plot illustrates the relationship between count and coloring. The plots represent pseudo-colored 2D matrices showing observed, expected, and observed/expected distribution, calculated from the overall frequencies of fragments on each of the axes. These show the relation between **a.-b**. H2A.Z and ACAA2, **c.-d.** H2A.Z and OGDH, **e.-f.** ACAA2 and OGDH, **g.-h.** H2A.Z and H3K9ac, all in the sham and TAC hearts. The pseudo-color corresponds to the Obs/Exp ratio, and the color intensity is proportional to the log2 of the number of observed fragments within each bin. These plots suggest that there is a positive correlation between the levels of H2A.Z and ACAA2 or OGDH, where the red indicates that this occurs more frequently than expected by chance, as denoted by the correlation coefficient listed above each plot. **i.** A histogram showing the relation between H3K9ac and the input, as an example of unrelated binding events, and **j.** as an example of a perfect correlation. This figure was generated by EaSeq software.

**Figure 7S.**
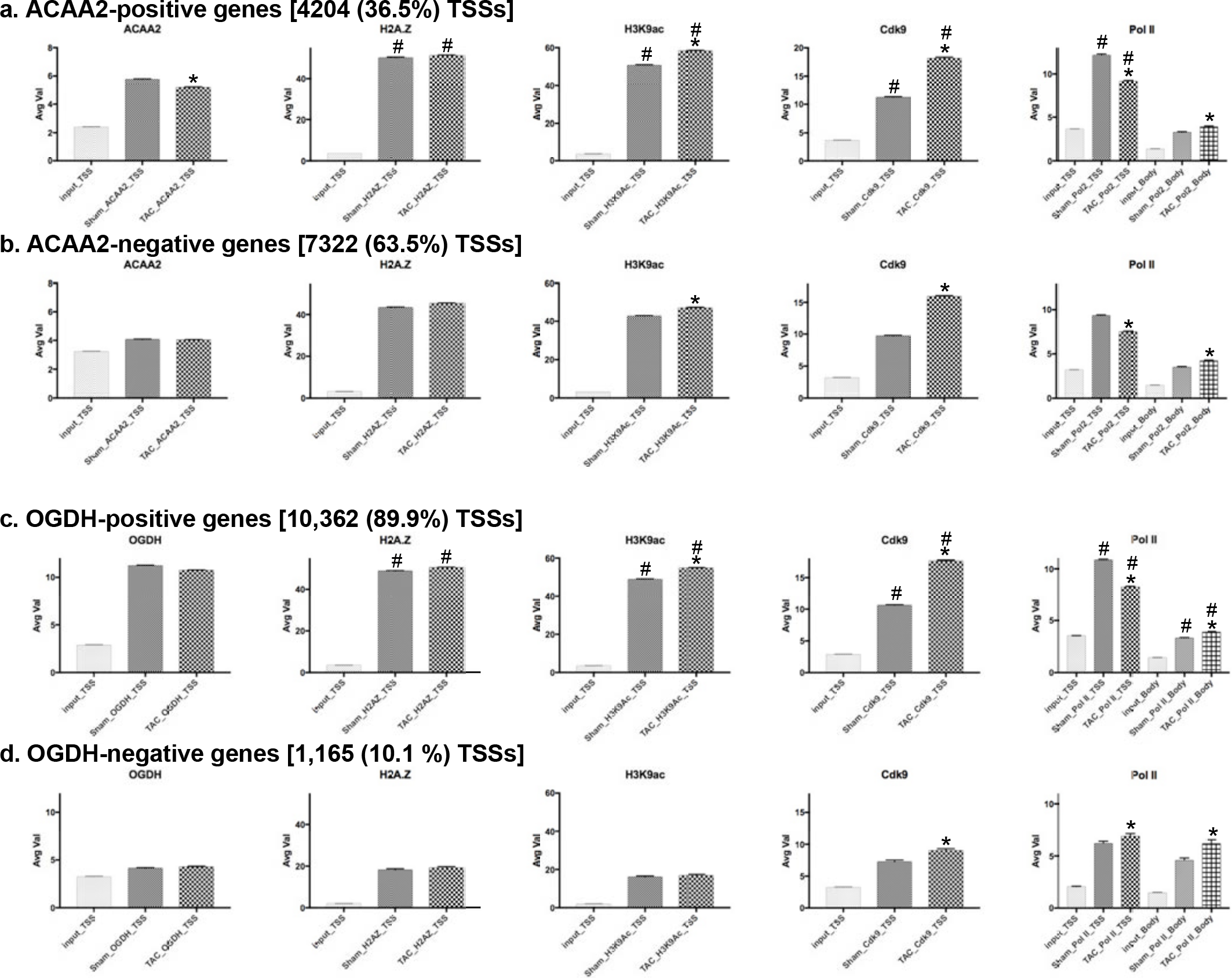
The mean for the average values of sequence Tags at the TSS of ACAA2- and OGDH-, positive and negative genes. Expressed genes (RNA pol II positive) were sorted into 4 groups: a. ACAA2-positive, b. ACAA2-negative, c. OGDH-positive, and d. OGDH-negative. The mean of the average values (AvgVal) of sequence Tags for ACAA2, OGDH, H2A.Z, H3K9ac, Cdk9, and pol II, at the TSS and gene body (pol II only), were calculated and plotted. Error bars represent standard error of the mean, and *p ≤ 0.05 v. sham in same plot, # p ≤ 0.05 v. corresponding data point in the ACCA2-negative or OGDH-negative gene subsets.

**Figure 8S.**
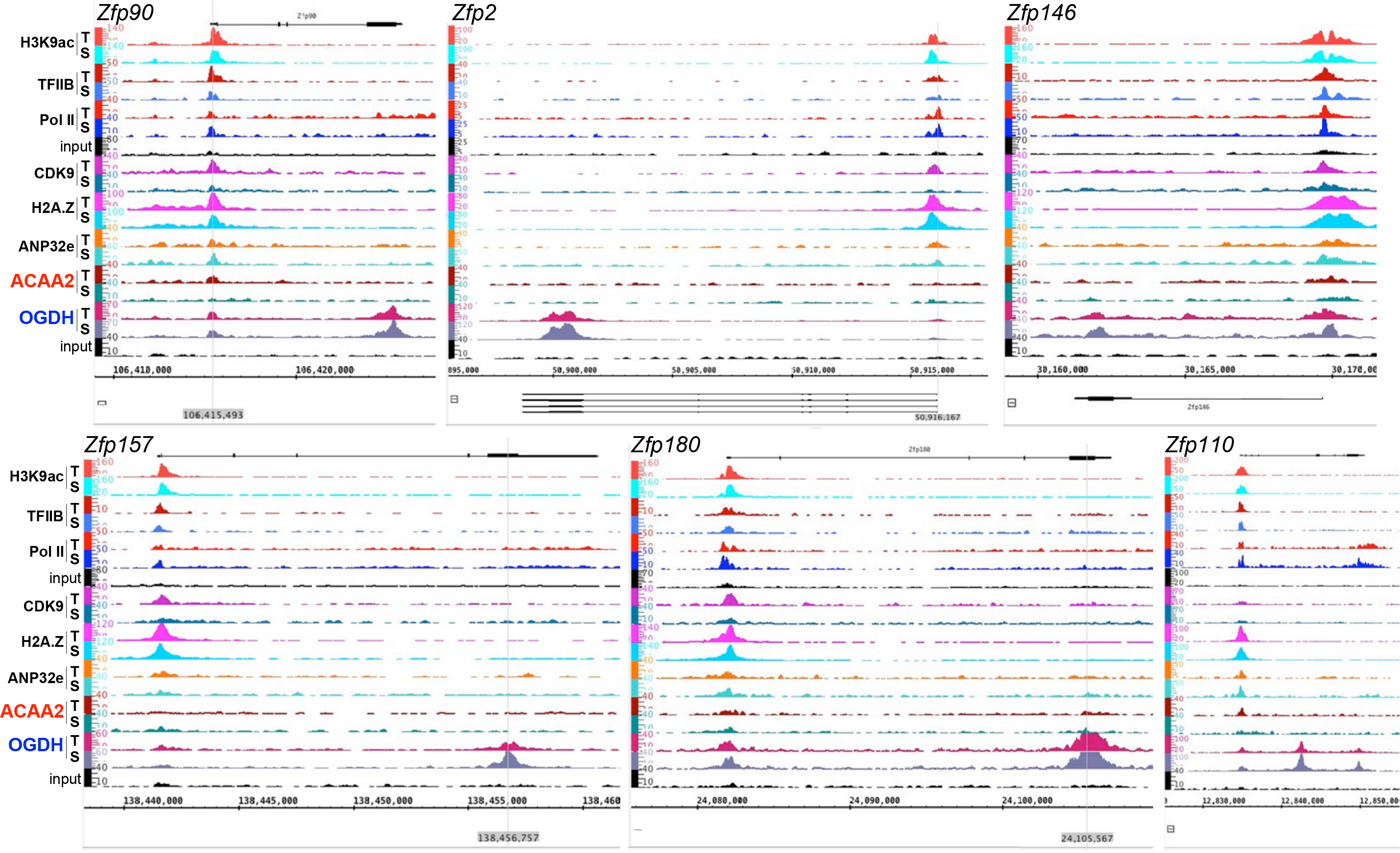
OGDH binds to the terminal exons of zinc finger proteins, in a H2A.Z-independent manner. The alignment of the ChIP-Seq sequence tags for H3K9ac, TFIIB, pol II, Cdk9, H2A.Z, ANP32E, ACAA2, and OGDH (Y-axis) across the genomic coordinates (X-axis) of *Zfp90, Zfp2, Zfp146, Zfp157, Zfp180, and Zfp110 genes*. The arrow shows the start and direction of transcription. The results show a substantial peak of OGDH in the terminal exons of these genes that is subject to differential regulation during cardiac hypertrophy. In addition, all genes show OGDH at their TSSs, albeit at a lower density.

**Fig. 9S-a-b.**
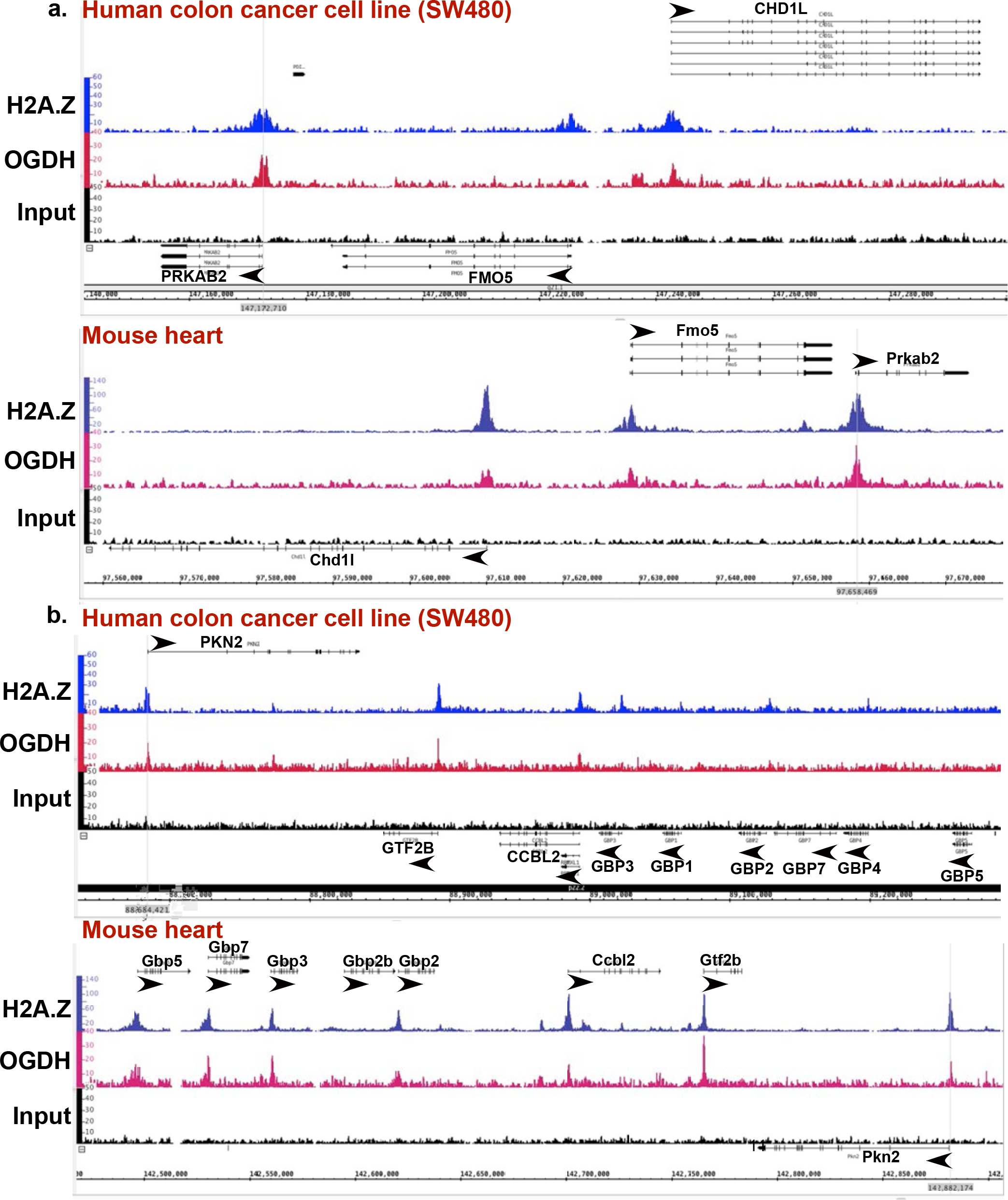
Conserved, selective, binding of OGDH to H2A.Z-bound TSSs. Both mouse heart tissue and human SW480 colon cancer cell line, were subjected to H2A.Z and OGDH ChIP-Seq using the same antibodies and chromatin concentration. The resulting sequence Tags form both reactions were aligned across the coordinates for the same genes in the mouse and human genomes, as indicated. Two regions are shown, **a.** the first showing the PRKAB2, FMO4, and CHD1L genes in the human cells and mouse tissue, where OGDH co-localizes with H2A.Z at the TSS in all three genes in the latter, however, OGDH is absent in from the FMO5 in the human genome, **b.** the second region encompasses PKN2, GTF2B, CCBL2, GBP1-5,7 genes that show conserved co-localization of OGDH and H2A.Z at the TSS of the former 3 genes in the mouse and human, but differs between species for the GBP genes, which have no OGDH in of the human cells, with a relatively small peak of H2A.Z at the TSS of GBP1-4. These data reveal that the co-localization of H2A.Z and OGDH at TSSs of key specific genes are conserved between species.

**Fig. 9S-c.**
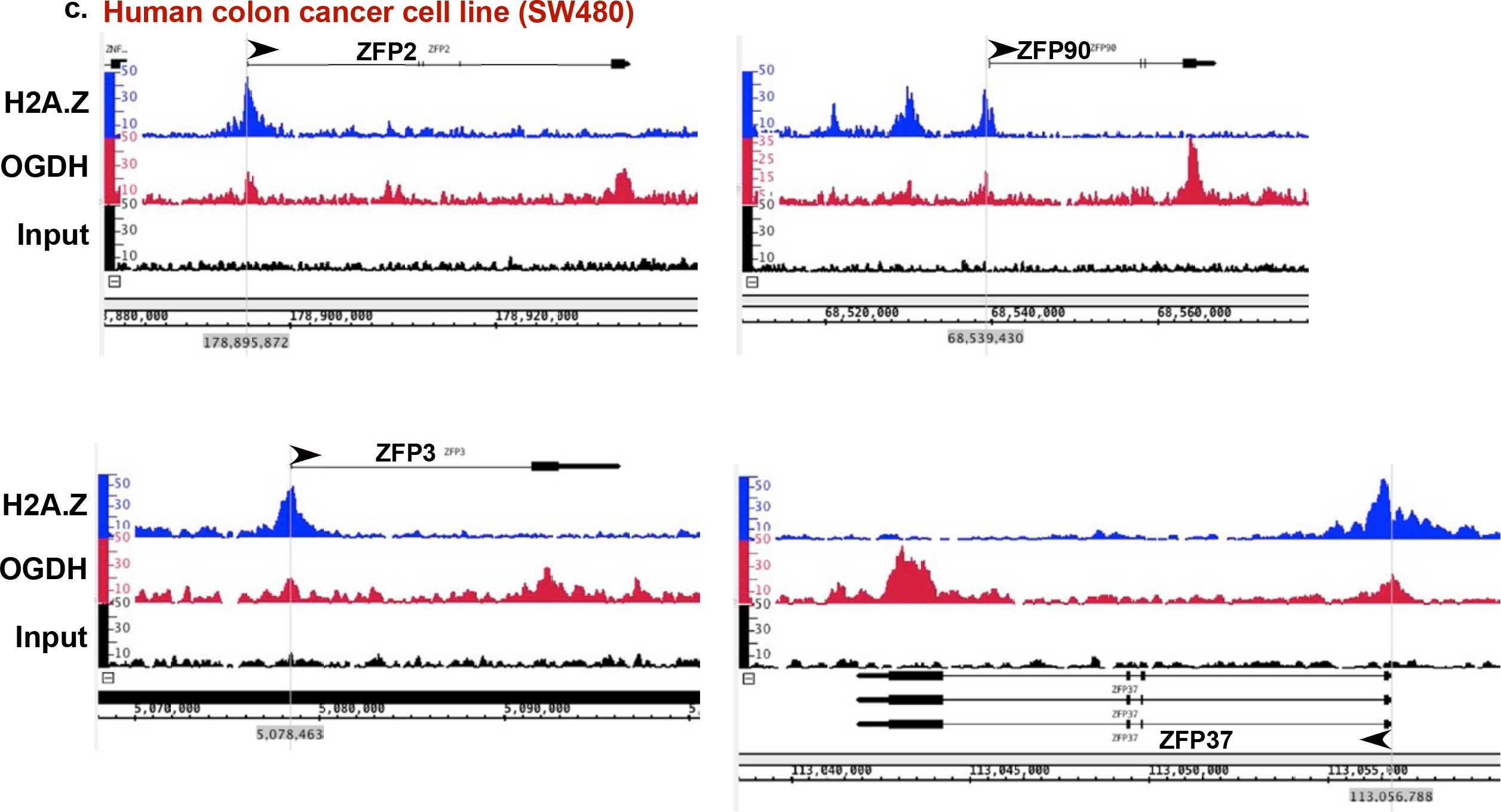
H2A.Z-independent binding of OGDH to the terminal exon in ZFP genes in humans. **The** human SW480 colon cancer cell line, was subjected to H2A.Z and OGDH ChIP-Seq using the same antibodies and chromatin concentration applied in the mouse heart tissue ChIP-Seq. The resulting sequence tags form both reactions (Y-axis) were aligned across the genome’s coordinates (X-axis). The results show 4 examples of ZFP genes in which OGDH is present in their terminal exon in the absence of H2A.Z, similar to what we observed in the mouse tissue (see Fig. 8S).

**Figure 10S.**
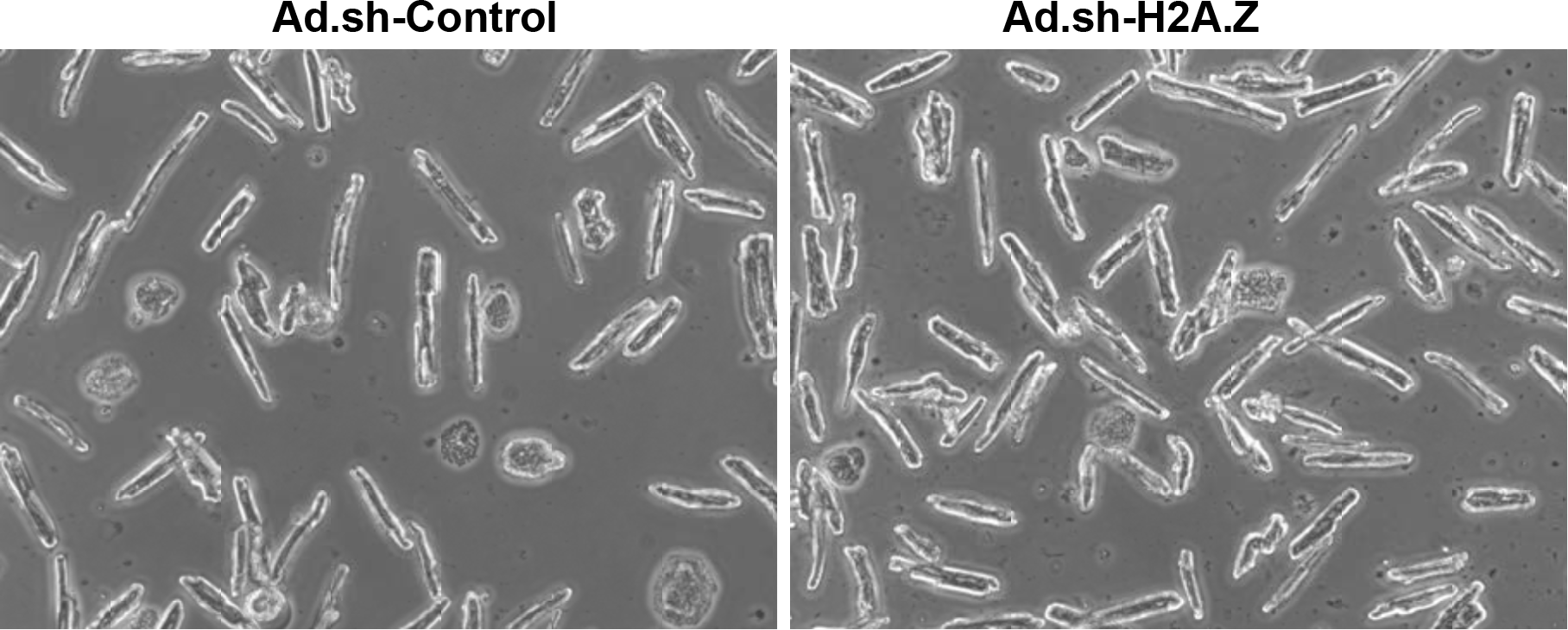
H2A.Z knockdown does not impact the viability of mouse adult cardiac myocytes. Mouse adult cardiac myocytes were isolated from the heart of 8 wk old male C57/Bl mice. They were then infected with 30 moi of adenoviruses harboring a nonsense shRNA control or one targeting H2A.Z. After 24 h, the cells were imaged before organelles were extracted for the Western blot analysis reported in Fig. 8. At this time point, H2A.Z knockdown did not affect cell viability, as evidenced by the maintenance of rod shape morphology of the cells.

**Figure 11S.**
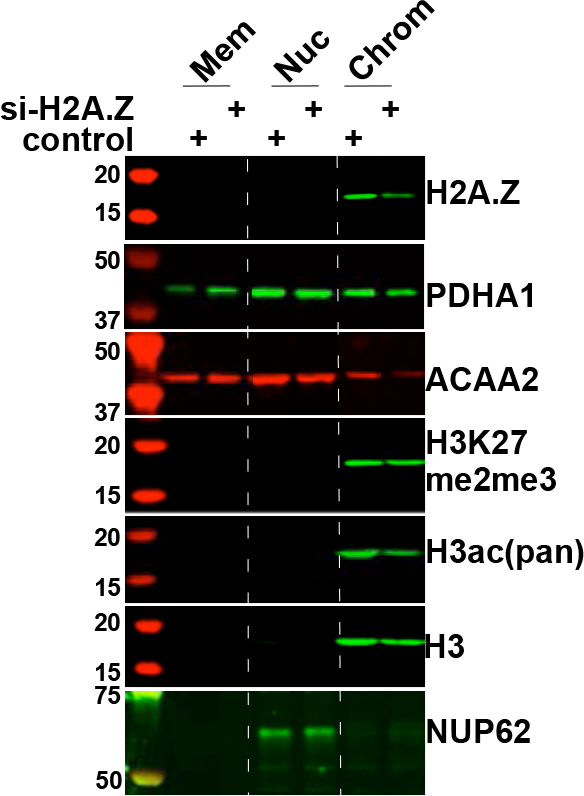
Knockdown of H2A.Z inhibits chromatin binding of metabolic enzymes and reduces histone modifications. Rat neonatal cardiac myocytes were isolated from the hearts of 1 day old Sprague Dawley pups. They were then infected with 30 moi of adenoviruses harboring a nonsense shRNA control or one targeting H2A.Z. After 24 h, organelles were isolated and fractionated into membrane/mitochondrial (Mem), nuclear (Nuc), and chromatin-bound (Chrom, no crosslinking applied), using a combination of differential lysis and sequential centrifugation. The proteins extracted from each of these fractions were analyzed by Western blotting for the genes indicated on the right of each panel.

**Figure 12S.**
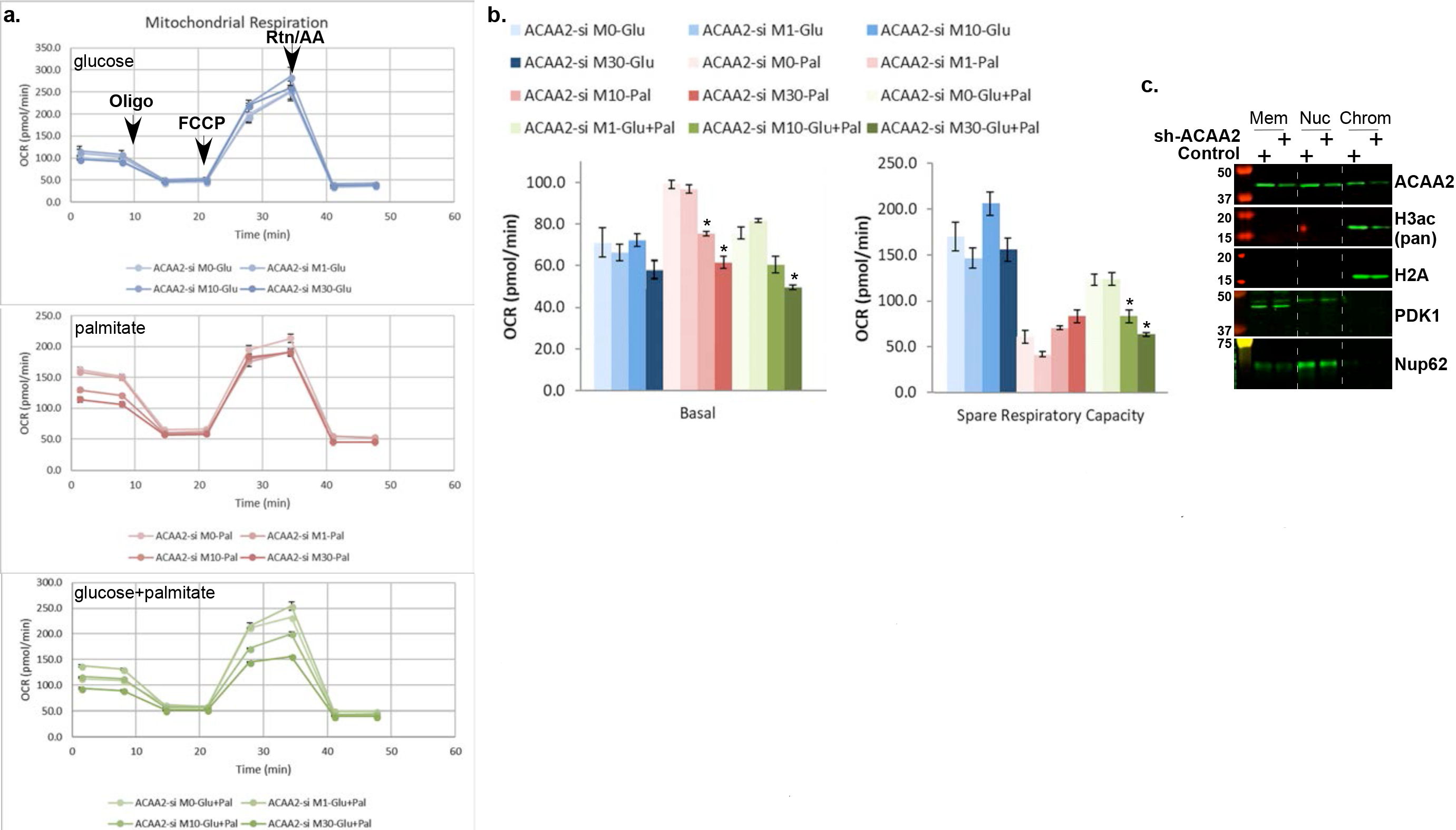
Knockdown of ACAA2 decreases fatty acid-dependent basal oxygen consumption rates (OCR) and H3 acetylation. Neonatal myocytes were infected with increasing doses of Ad harboring shRNA ACAA2 (ACAA2-si) or a control, in the presence of glucose, palmitate-BSA, or both. After 24h, cells were analyzed by the Seahorse analyzer for oxygen consumption rates (OCR). **a.** The results are graphed in real time showing basal, ATP-linked (after oligomycin injection), maximum (after FCCP injection), and mitochondrial [after rotenone (Rtn) / antimycin A (AA) injection] OCR. **b**. Basal and spare mitochondrial respiratory capacities are calculated plotted, \**p*<0.01 v. control, n=10, each. **c.** Neonatal myocytes were infected with 20 moi of the Ad.sh-ACAA2 or a control. After 24h, organelles were extracted and fractionated into membrane/mitochondrial (Mem), nuclear (Nuc), and chromatin-bound (Chrom) protein fractions, and analyzed by Western blotting for the indicated antibodies (right).

**Figure 13S.**
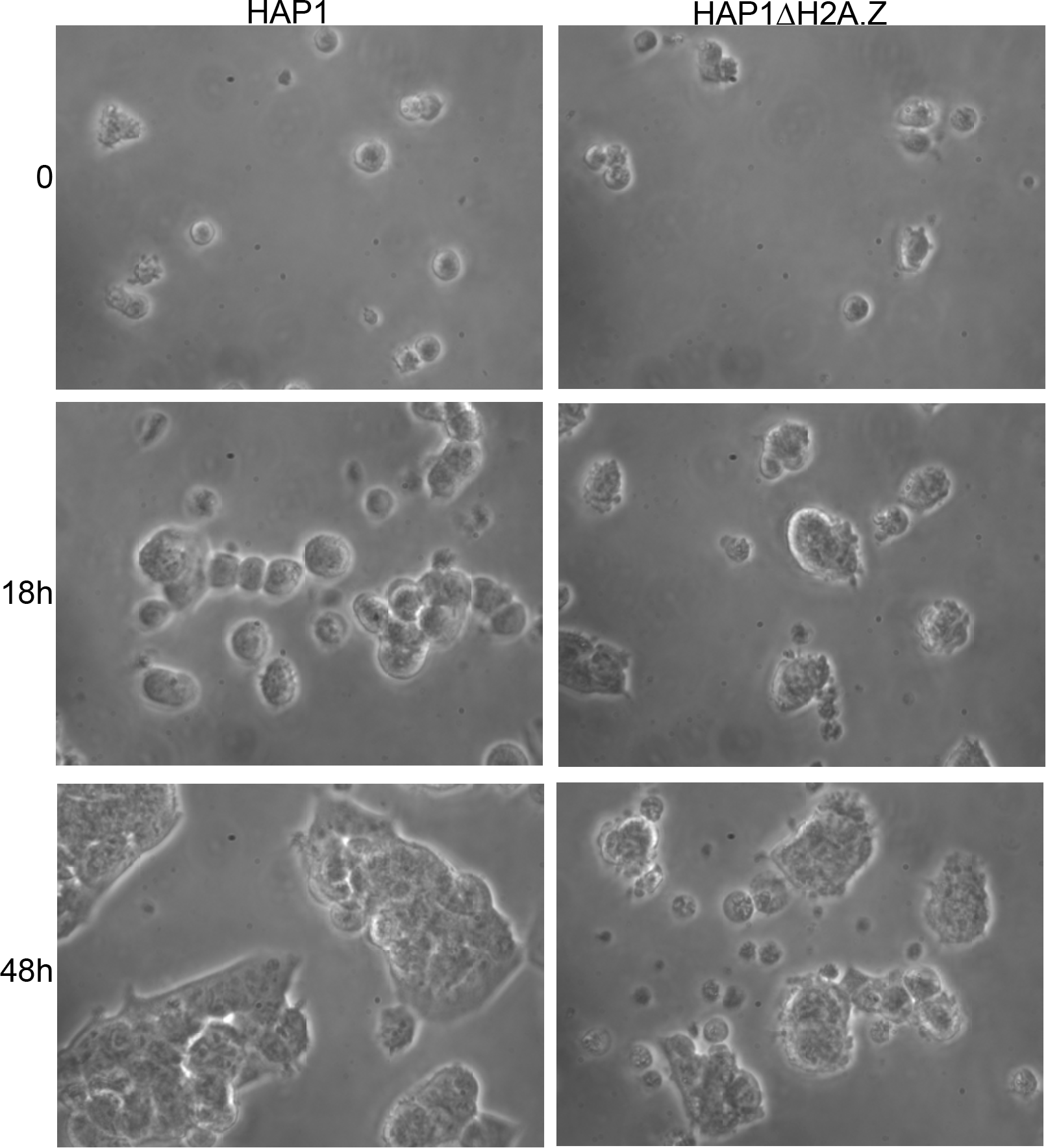
HAP1∆H2A.Z are viable but proliferate at a ~1/4 the rate of the parent cells. HAP1 and HAP1∆H2A.Z were purchased from Horizon Discovery and cultured according to the company’s protocol on gelatin coated glass slides. On day 0 equal number of cells were seed and imaged live at 18 and 48 h after that. Cells were counted at each time point in 3 fields revealing a ~4:1 ratio of parent:∆H2A.Z cell numbers.

**Figure 14S.**
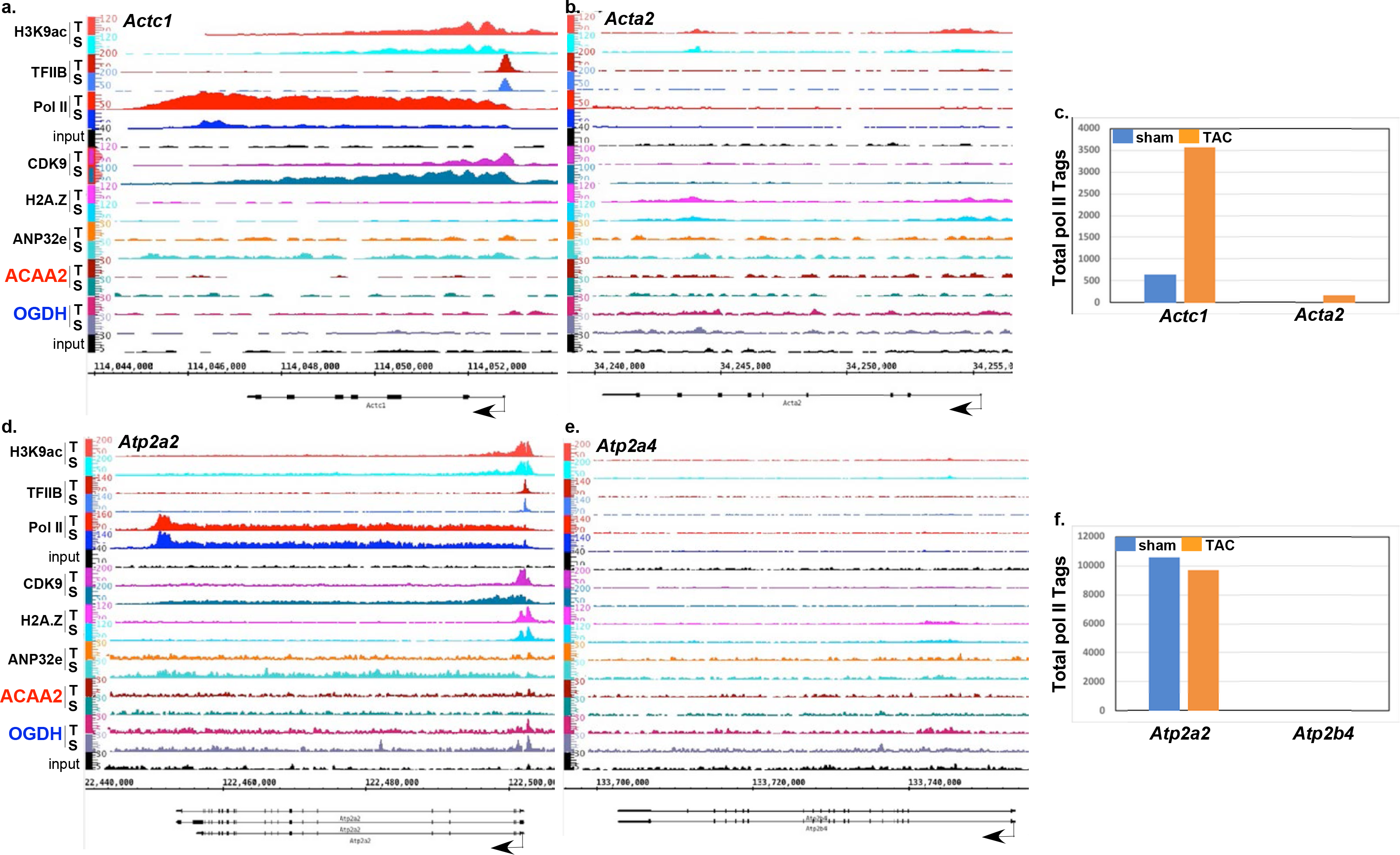
ChIP-Seq tags from the heart tissue are predominantly derived from cardiac myocytes v. myofibroblasts, endothelial, or smooth muscle cells. Mice were subjected to a sham or TAC operation. One-week post-TAC, the hearts were isolated and analyzed by ChIP-Seq for the indicated proteins (left). The alignment of the ChIP-Seq sequence Tags (Y-axis) for H3K9ac, TFIIB, pol II, Cdk9, H2A.Z, ANP32E, ACAA2, and OGDH across the genomic coordinates of **a.** *Actc1,* **b.** *Acta2,* **d.** *Atp2a2,* and **e.** *Atp2a4* genes (Y-axis). **c.** and **f.** are graphs of the total number of pol II Tags for each of those genes. Smooth muscle actin (*Acta2*), which is expressed in smooth muscle cells and myofibroblasts in the heart, and ATPase plasma membrane Ca^2+^ transporting 4 (*Atp2b4*), which is ubiquitously expressed, including in epithelial cells, have no dectable binding of pol II compared to its high abundance in the corresponding cardiac genes, cardiac actin (*Actc1*) and ATPase sarcoplasmic/endoplasmic reticulum Ca^2+^ transporting 2 (*Atp2a2*).

**Figure 15S.**
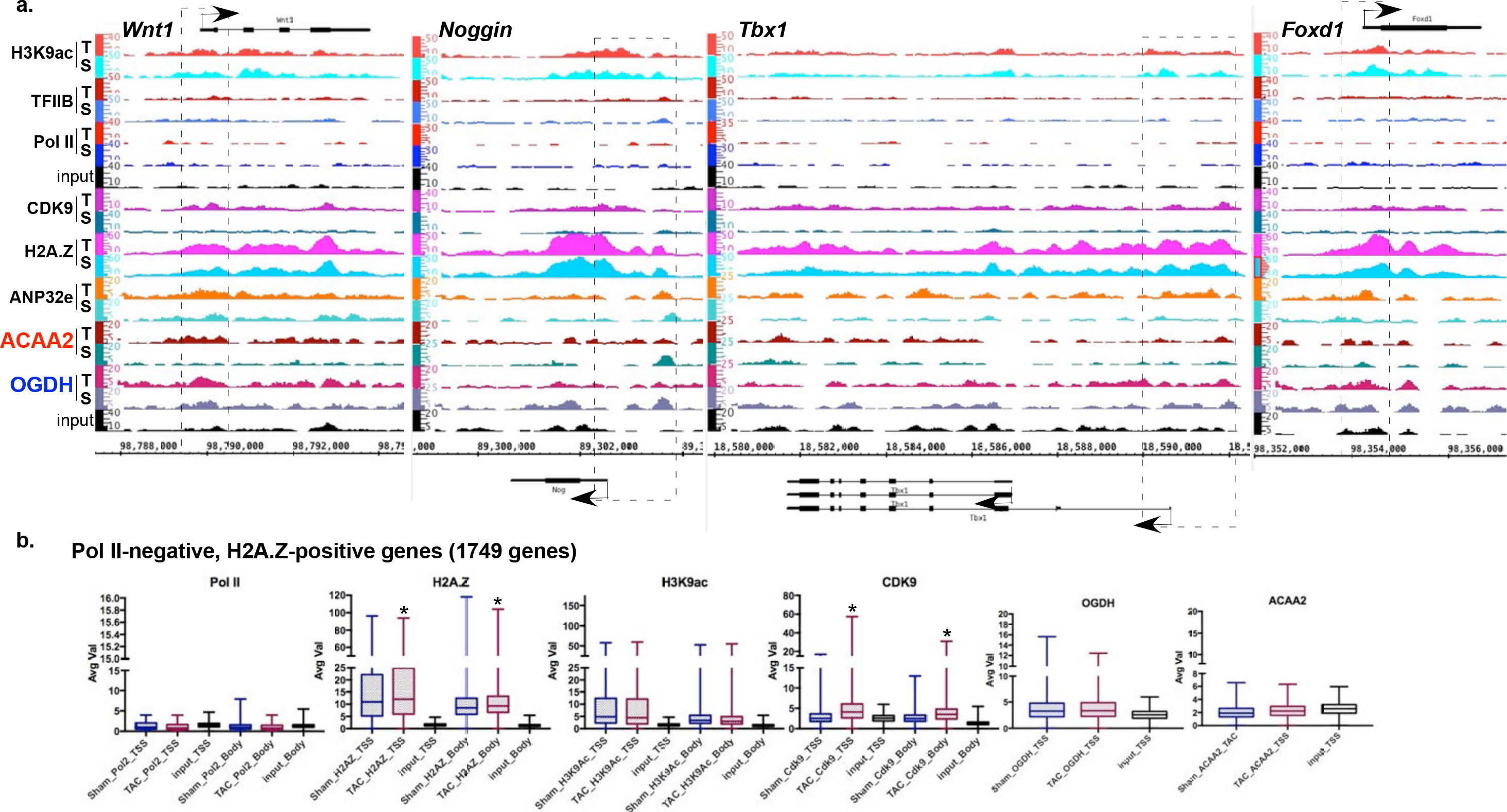
H2A.Z associates with the suppressed development genes. Mice were subjected to a sham or TAC operation. One-week post-TAC, the hearts were isolated and analyzed by ChIP-Seq for ACAA2. **a.** The alignment of the ChIP-Seq sequence Tags for H3K9ac, TFIIB, pol II, Cdk9, H2A.Z, ANP32E, ACAA2, and OGDH across the genomic coordinates of *Wnt1*, *Noggin*, *Tbx1*, and *Foxd1*, developmental genes. **b.** Genes were sorted for those that were pol II-negative genes and H2A.Z-posirive. The ChIP-Seq sequence Tags for ACAA2, H2A.Z, H3K9ac, OGDH, and pol II, at the TSS (−1000 to +1000) and across the gene body (+1000-gene end), were plotted as box plots (median and quartiles).

## REFERENCES

1. Marzluff WF, Gongidi P, Woods KR, Jin J and Maltais LJ. The human and mouse replication-dependent histone genes. Genomics. 2002;80:487–98.

2. Meneghini MD, Wu M and Madhani HD. Conserved histone variant H2A.Z protects euchromatin from the ectopic spread of silent heterochromatin. Cell. 2003;112:725–36.

3. Li B, Pattenden SG, Lee D, Gutierrez J, Chen J, Seidel C, Gerton J and Workman JL. Preferential occupancy of histone variant H2AZ at inactive promoters influences local histone modifications and chromatin remodeling. Proc Natl Acad Sci U S A. 2005;102:18385–90. Epub 2005 Dec 12.

4. Zhang H, Roberts DN and Cairns BR. Genome-wide dynamics of Htz1, a histone H2A variant that poises repressed/basal promoters for activation through histone loss. Cell. 2005;123:219–31.

5. Raisner RM, Hartley PD, Meneghini MD, Bao MZ, Liu CL, Schreiber SL, Rando OJ and Madhani HD. Histone variant H2A.Z marks the 5' ends of both active and inactive genes in euchromatin. Cell. 2005;123:233–48.

6. Leach TJ, Mazzeo M, Chotkowski HL, Madigan JP, Wotring MG and Glaser RL. Histone H2A.Z is widely but nonrandomly distributed in chromosomes of Drosophila melanogaster. J Biol Chem. 2000;275:23267–72.

7. Weber CM, Ramachandran S and Henikoff S. Nucleosomes are context-specific, H2A.Z-modulated barriers to RNA polymerase. Mol Cell. 2014;53:819–30. doi: 10.1016/j.molcel.2014.02.014.

8. Creyghton MP, Markoulaki S, Levine SS, Hanna J, Lodato MA, Sha K, Young RA, Jaenisch R and Boyer LA. H2AZ is enriched at polycomb complex target genes in ES cells and is necessary for lineage commitment. Cell. 2008;135:649–61. doi: 10.1016/j.cell.2008.09.056. Epub 2008 Nov 6.

9. van der Knaap JA and Verrijzer CP. Undercover: gene control by metabolites and metabolic enzymes. Genes Dev. 2016;30:2345–2369. doi: 10.1101/gad.289140.116.

10. Wellen KE, Hatzivassiliou G, Sachdeva UM, Bui TV, Cross JR and Thompson CB. ATP-citrate lyase links cellular metabolism to histone acetylation. Science. 2009;324:1076–80. doi: 10.1126/science.1164097.

11. Yoshii Y, Furukawa T, Yoshii H, Mori T, Kiyono Y, Waki A, Kobayashi M, Tsujikawa T, Kudo T, Okazawa H, Yonekura Y and Fujibayashi Y. Cytosolic acetyl-CoA synthetase affected tumor cell survival under hypoxia: the possible function in tumor acetyl-CoA/acetate metabolism. Cancer Sci. 2009;100:821–7.

12. Sutendra G, Kinnaird A, Dromparis P, Paulin R, Stenson TH, Haromy A, Hashimoto K, Zhang N, Flaim E and Michelakis ED. A nuclear pyruvate dehydrogenase complex is important for the generation of acetyl-CoA and histone acetylation. Cell. 2014;158:84–97. doi: 10.1016/j.cell.2014.04.046.

13. Nagaraj R, Sharpley MS, Chi F, Braas D, Zhou Y, Kim R, Clark AT and Banerjee U. Nuclear Localization of Mitochondrial TCA Cycle Enzymes as a Critical Step in Mammalian Zygotic Genome Activation. Cell. 2017;168:210–223.e11. doi: 10.1016/j.cell.2016.12.026. Epub 2017 Jan 12.

14. Wang Y, Guo YR, Liu K, Yin Z, Liu R, Xia Y, Tan L, Yang P, Lee JH, Li XJ, Hawke D, Zheng Y, Qian X, Lyu J, He J, Xing D, Tao YJ and Lu Z. KAT2A coupled with the alpha-KGDH complex acts as a histone H3 succinyltransferase. Nature. 2017;552:273–277. doi: 10.1038/nature25003. Epub 2017 Dec 6.

15. Haraguchi CM, Mabuchi T and Yokota S. Localization of a mitochondrial type of NADP-dependent isocitrate dehydrogenase in kidney and heart of rat: an immunocytochemical and biochemical study. J Histochem Cytochem. 2003;51:215–26. doi: 10.1177/002215540305100210.

16. Qattan AT, Radulovic M, Crawford M and Godovac-Zimmermann J. Spatial distribution of cellular function: the partitioning of proteins between mitochondria and the nucleus in MCF7 breast cancer cells. J Proteome Res. 2012;11:6080–101. doi: 10.1021/pr300736v. Epub 2012 Nov 8.

17. Shin H, He M, Yang Z, Jeon YH, Pfleger J, Sayed D and Abdellatif M. Transcriptional regulation mediated by H2A.Z via ANP32e-dependent inhibition of protein phosphatase 2A. Biochim Biophys Acta. 2018;1861:481–496. doi: 10.1016/j.bbagrm.2018.03.002. Epub 2018 Mar 8.

18. Mohammed H, Taylor C, Brown GD, Papachristou EK, Carroll JS and D’Santos CS. Rapid immunoprecipitation mass spectrometry of endogenous proteins (RIME) for analysis of chromatin complexes. Nat Protoc. 2016;11:316–26. doi: 10.1038/nprot.2016.020. Epub 2016 Jan 21.

19. Uhlen M, Fagerberg L, Hallstrom BM, Lindskog C, Oksvold P, Mardinoglu A, Sivertsson A, Kampf C, Sjostedt E, Asplund A, Olsson I, Edlund K, Lundberg E, Navani S, Szigyarto CA, Odeberg J, Djureinovic D, Takanen JO, Hober S, Alm T, Edqvist PH, Berling H, Tegel H, Mulder J, Rockberg J, Nilsson P, Schwenk JM, Hamsten M, von Feilitzen K, Forsberg M, Persson L, Johansson F, Zwahlen M, von Heijne G, Nielsen J and Ponten F. Proteomics. Tissue-based map of the human proteome. Science. 2015;347:1260419. doi: 10.1126/science.1260419.

20. Stark C, Breitkreutz BJ, Reguly T, Boucher L, Breitkreutz A and Tyers M. BioGRID: a general repository for interaction datasets. Nucleic Acids Res. 2006;34:D535–9. doi: 10.1093/nar/gkj109.

21. Wratting D, Thistlethwaite A, Harris M, Zeef LA and Millar CB. A conserved function for the H2A.Z C terminus. J Biol Chem. 2012;287:19148–57. doi: 10.1074/jbc.M111.317990. Epub 2012 Apr 9.

22. Huttlin EL, Ting L, Bruckner RJ, Gebreab F, Gygi MP, Szpyt J, Tam S, Zarraga G, Colby G, Baltier K, Dong R, Guarani V, Vaites LP, Ordureau A, Rad R, Erickson BK, Wuhr M, Chick J, Zhai B, Kolippakkam D, Mintseris J, Obar RA, Harris T, Artavanis-Tsakonas S, Sowa ME, De Camilli P, Paulo JA, Harper JW and Gygi SP. The BioPlex Network: A Systematic Exploration of the Human Interactome. Cell. 2015;162:425–440. doi: 10.1016/j.cell.2015.06.043.

23. Cai Y, Jin J, Yao T, Gottschalk AJ, Swanson SK, Wu S, Shi Y, Washburn MP, Florens L, Conaway RC and Conaway JW. YY1 functions with INO80 to activate transcription. Nat Struct Mol Biol. 2007;14:872–4. doi: 10.1038/nsmb1276. Epub 2007 Aug 26.

24. Latrick CM, Marek M, Ouararhni K, Papin C, Stoll I, Ignatyeva M, Obri A, Ennifar E, Dimitrov S, Romier C and Hamiche A. Molecular basis and specificity of H2A.Z-H2B recognition and deposition by the histone chaperone YL1. Nat Struct Mol Biol. 2016;23:309–16. doi: 10.1038/nsmb.3189. Epub 2016 Mar 14.

25. Mao Z, Pan L, Wang W, Sun J, Shan S, Dong Q, Liang X, Dai L, Ding X, Chen S, Zhang Z, Zhu B and Zhou Z. Anp32e, a higher eukaryotic histone chaperone directs preferential recognition for H2A.Z. Cell Res. 2014;24:389–99. doi: 10.1038/cr.2014.30. Epub 2014 Mar 11.

26. Kristensen AR, Gsponer J and Foster LJ. A high-throughput approach for measuring temporal changes in the interactome. Nat Methods. 2012;9:907–9. doi: 10.1038/nmeth.2131. Epub 2012 Aug 5.

27. Brickner DG, Cajigas I, Fondufe-Mittendorf Y, Ahmed S, Lee PC, Widom J and Brickner JH. H2A.Z-mediated localization of genes at the nuclear periphery confers epigenetic memory of previous transcriptional state. PLoS Biol. 2007;5:e81.

28. Light WH, Brickner DG, Brand VR and Brickner JH. Interaction of a DNA zip code with the nuclear pore complex promotes H2A.Z incorporation and INO1 transcriptional memory. Mol Cell. 2010;40:112–25. doi: 10.1016/j.molcel.2010.09.007.

29. Zovkic IB, Paulukaitis BS, Day JJ, Etikala DM and Sweatt JD. Histone H2A.Z subunit exchange controls consolidation of recent and remote memory. Nature. 2014;515:582–6. doi: 10.1038/nature13707. Epub 2014 Sep 14.

30. Shen T, Ji F, Wang Y, Lei X, Zhang D and Jiao J. Brain-specific deletion of histone variant H2A.z results in cortical neurogenesis defects and neurodevelopmental disorder. Nucleic Acids Res. 2018;46:2290–2307. doi: 10.1093/nar/gkx1295.

31. Mews P, Donahue G, Drake AM, Luczak V, Abel T and Berger SL. Acetyl-CoA synthetase regulates histone acetylation and hippocampal memory. Nature. 2017;546:381–386. doi: 10.1038/nature22405. Epub 2017 May 31.

32. Jiang Y, Qian X, Shen J, Wang Y, Li X, Liu R, Xia Y, Chen Q, Peng G, Lin SY and Lu Z. Local generation of fumarate promotes DNA repair through inhibition of histone H3 demethylation. Nat Cell Biol. 2015;17:1158–68. doi: 10.1038/ncb3209. Epub 2015 Aug 3.

33. Coleman-Derr D and Zilberman D. Deposition of histone variant H2A.Z within gene bodies regulates responsive genes. PLoS Genet. 2012;8:e1002988. doi: 10.1371/journal.pgen.1002988. Epub 2012 Oct 11.

34. Nesvizhskii AI, Keller A, Kolker E and Aebersold R. A statistical model for identifying proteins by tandem mass spectrometry. Anal Chem. 2003;75:4646–58.

35. Sayed D, Yang Z, He M, Pfleger J and Abdellatif M. Acute Targeting of General Transcription Factor IIB Restricts Cardiac Hypertrophy via Selective Inhibition of Gene Transcription. Circ HF. 2015;8:138–148.

36. Ackers-Johnson M, Li PY, Holmes AP, O’Brien SM, Pavlovic D and Foo RS. A Simplified, Langendorff-Free Method for Concomitant Isolation of Viable Cardiac Myocytes and Nonmyocytes From the Adult Mouse Heart. Circ Res. 2016;119:909–20. doi: 10.1161/CIRCRESAHA.116.309202. Epub 2016 Aug 8.

37. Han M, Yang Z, Sayed D, He M, Gao S, Lin L, Yoon SH and Abdellatif M. GATA4 Expression is Primarily Regulated via a miR-26b-Dependent Posttranscriptional Mechanism During Cardiac Hypertrophy. Cardiovasc Res. 2012;93:645–54.

38. Sayed D, Hong C, Chen IY, Lypowy J and Abdellatif M. MicroRNAs play an essential role in the development of cardiac hypertrophy. Circ Res. 2007;100:416–24.

39. Chen IY, Lypowy J, Pain J, Sayed D, Grinberg S, Alcendor RR, Sadoshima J and Abdellatif M. Histone H2A.z is essential for cardiac myocyte hypertrophy but opposed by silent information regulator 2alpha. J Biol Chem. 2006;281:19369–77.

40. Lypowy J, Chen IY and Abdellatif M. An alliance between Ras GTPase-activating protein, filamin C, and Ras GTPase-activating protein SH3 domain-binding protein regulates myocyte growth. J Biol Chem. 2005;280:25717–28.

41. Rane S, He M, Sayed D, Vashistha H, Malhotra A, Sadoshima J, Vatner DE, Vatner SF and Abdellatif M. Downregulation of MiR-199a Derepresses Hypoxia-Inducible Factor-1{alpha} and Sirtuin 1 and Recapitulates Hypoxia Preconditioning in Cardiac Myocytes. Circ Res. 2009;104:879–886.

42. Sayed D, He M, Yang Z, Lin L and Abdellatif M. Transcriptional regulation patterns revealed by high resolution chromatin immunoprecipitation during cardiac hypertrophy. J Biol Chem. 2013;288:2546–58. doi: 10.1074/jbc.M112.429449. Epub 2012 Dec 10.

43. Lerdrup M, Johansen JV, Agrawal-Singh S and Hansen K. An interactive environment for agile analysis and visualization of ChIP-sequencing data. Nat Struct Mol Biol. 2016;23:349–57. doi: 10.1038/nsmb.3180. Epub 2016 Feb 29.

44. Nicol JW, Helt GA, Blanchard SG, Jr., Raja A and Loraine AE. The Integrated Genome Browser: free software for distribution and exploration of genome-scale datasets. Bioinformatics. 2009;25:2730–1.

